# The Metabolic Scope Theory of Aging: Rising Mitochondrial Impedance Compresses Metabolic Reserve to Constrain Lifespan

**DOI:** 10.64898/2026.06.18.730063

**Authors:** Gilad Lehmann, Yona Greenman, Izhar Shtrom, Yossi Anis, Joseph Lehmann, Naftali Stern, Gabi Shefer

**Affiliations:** Tel Aviv-Sourasky Medical Center (Ichilov), Tel Aviv, Israel; Independent Researcher, Atlit, Israel; The Sagol Center for Epigenetics and Institute of Endocrinology, Metabolism and Hypertension, Tel Aviv-Sourasky Medical Center, Tel Aviv, Israel; Tel-Aviv University, Tel Aviv, Israel

## Abstract

Maximum lifespan varies more than 100-fold across vertebrates, yet within each species aging emerges as a coordinated syndrome spanning metabolism, immunity, endocrine signaling, cognition, and regeneration. We propose the Metabolic Scope Theory of Aging (MSTA), which treats longevity as the time required to exhaust mitochondrial bioenergetic reserve rather than as a consequence of resting metabolic rate alone.

The framework decomposes lifespan into three physical axes: Scope, the reserve capacity that buffers cumulative damage; Stability, the resistance of mtDNA-linked OXPHOS architecture to erosion; and Pace, the temperature-dependent kinetics of lesion accumulation. Using body mass as a Scope proxy, mtDNA GC content as a Stability proxy, and body temperature as Pace, the resulting Scope–Stability–Pace relation, lnMLS = *α* lnBM + *β* GC% − *γ* Tb + c, explains ∼69% of mammalian maximum-lifespan variance across 379 species. Cross-class comparisons reinforce the same constraint structure: birds offset high thermal Pace through elevated mtDNA Stability, and the SSP temperature coefficient derived from mammals matches the temperature dependence of lifespan observed in ectotherms. The framework further connects comparative lifespan scaling to Gompertz-like mortality acceleration through progressive reserve erosion and threshold crossing.

Mechanistically, MSTA models cumulative mtDNA-linked damage as rising impedance within OXPHOS. Increasing internal resistance drives mitochondria toward a high-redox-pressure, low-current regime that preserves basal ATP while restricting NAD^+^ regeneration, CoQ acceptor availability, and Δp-dependent work. The earliest failure is therefore not energetic collapse but loss of regenerative scope: NAD^+^-gated TCA flux, aspartate and nucleotide synthesis, one-carbon metabolism, and redox-buffered repair become progressively harder to sustain, and diverse age- related pathologies emerge as tissue-specific projections of this shared upstream constraint. MSTA separates a reversible, operational impedance (redox poise, membrane potential, endocrine tone) from a fixed, informational one (accumulated mtDNA damage) that sets the hard ceiling on lifespan. Because the informational layer cannot be reversed by regulatory means, the framework predicts that until therapies can directly restore mitochondrial conductance, interventions will be most effective when they relieve redox pressure or bypass constrained biosynthetic gates.

## Introduction

> “Food restores what is lost, but it does not restore the principle by which restoration itself is governed.” —Aristotle, On Longevity and Shortness of Life (paraphrase; 350 B.C.)

Aging is an enduring paradox: metabolic turnover ceaselessly renews the material fabric of life, yet the template orchestrating this renewal succumbs to irreversible decay. Across the mammalian radiation—despite sharing orthologous protein-coding genes, conserved epigenetic machinery, and the same fundamental cellular enzymes—maximum lifespan varies 100-fold, from scarcely two years in shrews to over two centuries in bowhead whales. This reveals that the tempo of decay is exquisitely tunable. Within each species, however, late-life frailty emerges as a synchronized cluster: diverse tissues, despite disparate origins and turnover rates, decline in concert. This dual puzzle—radical variation across species, tight coordination within them—demands a unified explanation^1,2^.

In humans, the most intensively studied case, this synchronization is particularly dramatic: sarcopenia accompanies insulin resistance, cognitive decline parallels immune senescence, and frailty precipitates across systems with remarkable predictability. The erosion of bioenergetic reserve captures this trajectory quantitatively. In young, fit adults, peak factorial aerobic scope—the ratio of maximal to resting metabolism—reaches 8–10 fold^3^; this is a transient stress-response capacity, distinct from the sustained alimentary ceiling that caps day-to-week throughput at roughly 2.5-fold above basal expenditure^4^. Aging compresses the peak reserve and narrows the margin between basal maintenance, sustained throughput, and acute stress capacity, until maintenance and stress responses can no longer be funded together. The decline is not linear: aerobic capacity falls from ∼0.5%/year in the fourth decade to *>*2%/year by the seventh, consistent with progressive, self-reinforcing reserve erosion rather than a fixed chronological rate^5^.

Existing theories struggle to explain both synchronization and variation. Evolutionary and signaling frameworks account for decline’s existence but not its coordinated biochemical signature. Hallmark taxonomies and information-theoretic accounts of epigenetic change identify recurrent features and proximate drivers of aged tissue, but not the upstream physical constraint from which they follow. Mitochondrial-centric models historically faltered on bulk sequencing data suggesting mutation loads too low for causality. Recent single-cell analyses weaken this objection: cryptic mutations—variants unique to individual cells, masked by averaging—undergo clonal expansion, reaching heteroplasmies sufficient to impair local respiration^6^. Still, direct proof that physiological mtDNA loads pace normal aging is lacking. MSTA therefore treats this not as a finding but as a founding premise—cumulative OXPHOS impedance as the rate-limiting constraint on lifespan—and stands on the predictions that follow.

The mitochondrial genome is also physically exposed to a distinctive erosion process. Unlike nuclear DNA’s symmetric duplication, vertebrate mtDNA replicates asymmetrically: the heavy strand (H-strand) is displaced and remains single-stranded for extended intervals^7^. Exposed cytosine (and to a lesser degree adenine) undergoes spontaneous hydrolytic deamination at Arrhenius-governed rates. At the organismal level the fitted temperature dependence of lifespan corresponds to an effective acceleration of ∼2.5-fold per 10 *^◦^*C (activation energy ∼73 kJ/mol; §2.4), while the underlying chemical reactions may carry steeper *Q*_10_ values^8,9^. Unrepaired lesions fix as directional C→T and A→G transitions, progressively depleting the H-strand of deamination-prone bases. In humans, for example, the H-strand is depleted of cytosine (13.2%) and adenine (24.8%) while enriched in guanine (31.2%) and thymine (30.8%). The strong compositional asymmetry of vertebrate mtDNA is therefore the cumulative imprint of this replication-associated, temperature-sensitive chemical pressure^10^.

Because mtDNA erosion cannot be eliminated over long post-mitotic lifetimes, longevity must depend not on preventing damage but on tolerating it. This shifts the question from damage avoidance to damage buffering.

Post-mitotic tissues maintain oxidative capacity far exceeding basal requirements, and it is this reserve—not the absolute rate of energy turnover—that determines how much erosion can be absorbed before regenerative function fails. The respiratory chain can be idealized as a current-limited circuit, in which the NADH-to-O_2_ redox span provides the driving voltage *V_redox_*, electron flux is the current, and accumulated lesions raise the effective internal resistance *R_int_*. This is not a claim that any single lesion behaves as an identical resistor; it is a reduced description that collapses heterogeneous defects—point mutations, altered subunit geometry, impaired supercomplex assembly—into one impedance variable. Because these defects sit in series along the electron path, their contributions add: each lesion raises *R_int_* by a roughly fixed increment, so internal resistance grows approximately linearly as damage accumulates.

Writing normalized impedance as *Z_R_* = *R_int_/R_int,_*_0_, irreversible frailty occurs when *Z_R_* reaches a critical value *Z_R,crit_* at which respiration-gated biosynthesis can no longer be sustained above basal demand. The interval the system can traverse, *S_R_* = *Z_R,crit_* − 1, is the factorial aerobic reserve itself: maximal output corresponds to baseline resistance and the maintenance floor to critical resistance, so *R_int,crit_/R_int,_*_0_ = *MMR/BM R* ≡ *Scope* and *S_R_*≈ *Scope* − 1. With impedance rising at velocity *v_Z_* = *dZ_R_/dt*, maximum lifespan is the time required to cross the interval:

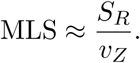

Lifespan is therefore linear in Scope—reserve divided by the rate at which it is spent—so the body-mass dependence of lifespan inherits that of Scope. Across mammals, factorial scope scales with body mass as *M* ^0^.^12–0.15^, predicting a lifespan allometric exponent of the same shallow magnitude^11,12^. What matters is not how fast an organism burns, but how much reserve it has to burn through.

### The Metabolic Scope Theory of Aging

We term this framework the Metabolic Scope Theory of Aging (MSTA). Longevity decomposes into three physical currencies under evolutionary tradeoff: Scope—the reserve capacity, indexed by body-mass scaling, that sets the traversable impedance interval; Stability—the resistance of mtDNA-linked OXPHOS architecture to erosion, indexed by mitochondrial GC content; and Pace—the temperature-dependent kinetics, indexed by body temperature, that set how fast that interval is consumed. Scope expands the damage budget; Stability and Pace set the velocity at which it is spent. These combine into the Scope–Stability–Pace (SSP) relationship,

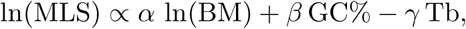

which is not an arbitrary regression but the log-linear form of a capacity term divided by a rate term, derived in full from the impedance-traversal equation in §2.6.

This same impedance picture explains why the first failure of aging is biosynthetic rather than energetic. As internal resistance rises, the chain drifts into a high-redox-pressure, low-current regime: upstream reducing equivalents accumulate as NADH and reduced CoQ even as downstream electron flux collapses^13^. Basal ATP can be defended through demand suppression and substrate rerouting, but respiration-gated biosynthesis cannot—NAD^+^-dependent TCA flux, aspartate and nucleotide synthesis, one-carbon metabolism, and redox-buffered repair become progressively harder to sustain. Diverse age-related pathologies emerge as tissue-specific projections of this single upstream constraint, resolving the paradox of synchronized senescence.

In the sections that follow, we test the SSP relationship across mammals and other vertebrate classes, examine exceptionally long-lived lineages as axis-optimization cases, connect reserve erosion to Gompertz-like mortality acceleration, and derive the circuit-level mechanisms by which rising OXPHOS impedance produces the ordered metabolic, endocrine, immune, and regenerative failures of aging—showing that mammalian, ectotherm, and interventional evidence converge as a consilience of independent observations.

## Results

### 2.1 The Scope–Stability–Pace (SSP) law explains ∼69% of mammalian lifespan variation

Maximum lifespan (MLS) scales across mammals as lnMLS∼0.148 lnBM, close to the theoretical exponent for factorial aerobic scope (∼0.15, derived from the allometric divergence between maximal and basal metabolic rate; Fig. 1).

**Figure 1.**
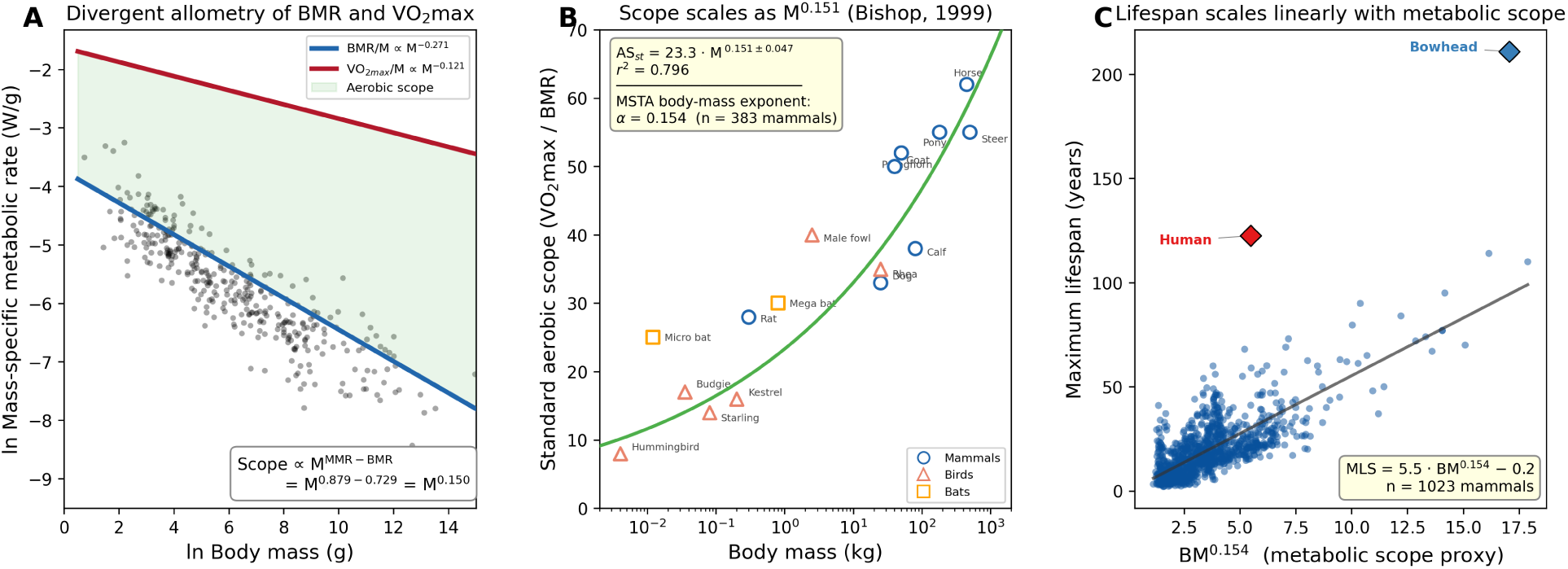
The body-mass exponent of lifespan derives from the allometric divergence between basal and maximal metabolic rate. **(A)** Mass-specific basal metabolic rates (BMR/M; black dots; n=350 mammalian species; AnAge database) plotted against ln body mass. Overlaid are the canonical allometric regression lines from Bishop^11^: mass-specific BMR ∝M^-0.271^ (blue) and mass-specific VO_2_max ∝M^-0.121^ (red), derived from whole-organism scalings of BMR ∝M^0.729^ and VO_2_max∝M^0.879^, respectively. Because the two rates scale with different exponents, per-gram maintenance cost declines steeply with body size while per-gram maximal capacity declines gently; their ratio—the factorial aerobic scope (green shaded area)—therefore widens with increasing body mass, scaling as M^0.879–0.729^=M^0.150^. The exact values of the BMR and MMR exponents vary modestly across studies depending on data filtering criteria and sample composition: White and Seymour^14^ report a stricter BMR exponent of 0.686±0.014 (n=571 species), while Weibel et al.^15^ report an MMR exponent of 0.872±0.029 (n=34 species). These alternative exponents yield a scope exponent of ∼0.18, bracketing the directly measured value in panel B. **(B)** Standard aerobic scope (VO_2_max/BMR) plotted against body mass for 9 mammalian species (blue circles), 6 bird species (red triangles), and 2 bat species (orange squares), redrawn from Bishop (1999), Figure 3b. The regression line (green) follows ASst=23.3·M^0.151^±0.047 (r^2^=0.796). This directly measured scope exponent of 0.151 is statistically indistinguishable from the MSTA body-mass exponent *α*=0.148 derived independently from longevity data (n=379 mammals), consistent with an allometric parallel between body size, reserve capacity, and lifespan. We interpret VO_2_max/BMR here as a whole-organism aerobic proxy for conductance reserve—the cellular OXPHOS headroom that buffers rising impedance (§3.1)—not as a direct measurement of that reserve; because the ratio is dominated by locomotor muscle, its athletic component need not track maximum lifespan (§3.10). **(C)** Maximum lifespan (MLS) plotted against BM^0.148^ (the MSTA body-mass exponent) for 1,018 mammalian species. If lifespan scales as a power law of body mass with exponent *α*, then plotting MLS against BM*α* should yield a linear relationship—as observed. The regression line follows MLS=6.13·BM^0.148^ − 1.03. Human (*Homo sapiens*, red diamond) and bowhead whale (*Balaena mysticetus*, blue diamond) are highlighted as positive outliers whose extended longevity is accounted for by the additional SSP pillars (GC content and body temperature, respectively).

To test whether mtDNA Stability and thermal Pace capture residual variance, we indexed Stability by mitochondrial GC content (GC%) and Pace by core body temperature (Tb). A log-linear model with all three coefficients freely fitted,

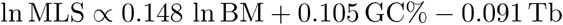

explained ∼69% of MLS variance (n=379 species, p<10^-90^; Fig. 2)^16,17,18,19^. Adding the SSP terms (GC% and Tb) to a body-mass-only model raised R^2^ from 0.56 to 0.69 (ΔR^2^≈0.13; joint F(2,375)=75.3, p<10^-27^), recovering ∼29% of the lifespan variance left unexplained by body mass alone.

**Figure 2.**
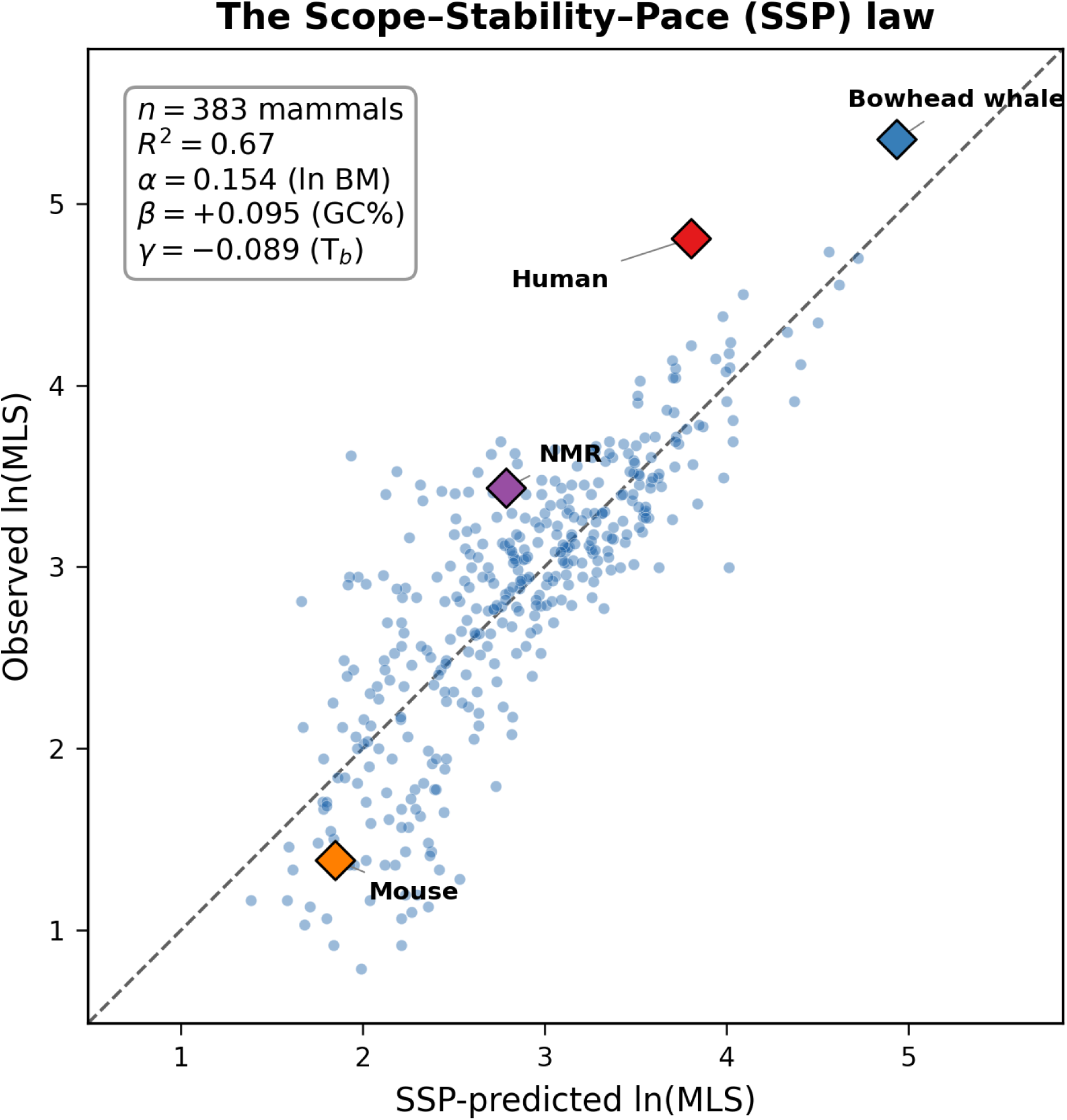
The Scope–Stability–Pace (SSP) law explains ∼69% of mammalian lifespan variance. A log-linear model of the form lnMLS=*α* lnBM+*β* GC%−*γ* Tb+c was fitted by ordinary least squares to n=379 mammalian species with complete data for body mass (BM), heavy-strand mitochondrial DNA guanine–cytosine content (GC%), and core body temperature (Tb). The plot shows SSP-predicted lnMLS (abscissa) versus observed lnMLS (ordinate); the dashed line is the identity (y=x). Fitted coefficients: Scope *α*=0.148 (ln BM), Stability *β*=+0.105 (GC%), Pace *γ*=+0.091 (Tb); R^2^=0.69, p=10^-90^. Diamond markers highlight four reference species: mouse (*Mus musculus*; orange), naked mole rat (NMR, *Heterocephalus glaber* ; purple), human (*Homo sapiens*; red), and bowhead whale (*Balaena mysticetus*; blue). Human and bowhead whale are positive outliers (observed > predicted), consistent with additional longevity mechanisms beyond the three SSP axes^20^.

All coefficient signs aligned with the mechanistic logic of MSTA: positive for Scope (larger BM extends lifespan via greater reserve), positive for Stability (higher GC% slows erosion), and negative for Pace (higher Tb accelerates lesion chemistry). Stability and Pace effects strengthened mutually in the full model (each coefficient’s magnitude increased when the other term was included), confirming they are coupled rather than redundant: duplex stabilization mitigates thermally activated damage. These coefficients are robust to phylogenetic non-independence: their signs are preserved in all 100 posterior mammalian trees under three models of trait evolution (Brownian motion, Pagel’s *λ*, and Ornstein–Uhlenbeck). Magnitudes attenuate under correction — least for Scope (*α*), and by roughly half for Stability (*β*) and Pace (*γ*), as deep-clade structure such as the endotherm body-temperature split absorbs part of their univariate variance — yet all three remain significant and correctly signed under every model (Methods; Supplementary Note S2.3).

Thus, the SSP law provides a compact quantitative framework for mammalian lifespan variation: Scope determines cumulative tolerable erosion, while Stability–Pace sets erosion rate. Because species occupy SSP coordinates shaped by selection rather than populating the space uniformly, the fitted coefficients recover the local geometry of the mammalian solution space — local longevity elasticities within an evolved constraint manifold, not arbitrary regression weights.

### 2.2 Co-evolution of Scope and Stability in endotherms

According to the asynchronous replication constraint described in the Introduction, the mtDNA H-strand is single-stranded for extended periods and vulnerable to spontaneous deamination—hydrolytic lesions that rise exponentially with temperature. Under this constraint, enriching heavy-strand guanine (reflected in whole-genome GC%) should provide thermal protection: GC-enriched H-strands carry fewer deamination-prone bases, reducing the target for thermally accelerated lesions during single-strand exposure.

Within-mammal relationships are confounded by body mass. Simple correlation between body temperature and mtDNA GC% in mammals is weak but significant (r=+0.16, p=0.002; n=379). However, this relationship disappears when controlling for body mass (partial r=+0.09, p=0.08), because larger mammals tend to be both warmer and GC-enriched. The robust relationship is between body mass and GC%—not temperature and GC% (partial r=+0.30, p<10^-6^ controlling for Tb; Fig. 3A). This suggests that within the narrow mammalian temperature range (∼30–41 °C), Stability co-evolves primarily with Scope rather than directly with Pace.

**Figure 3.**
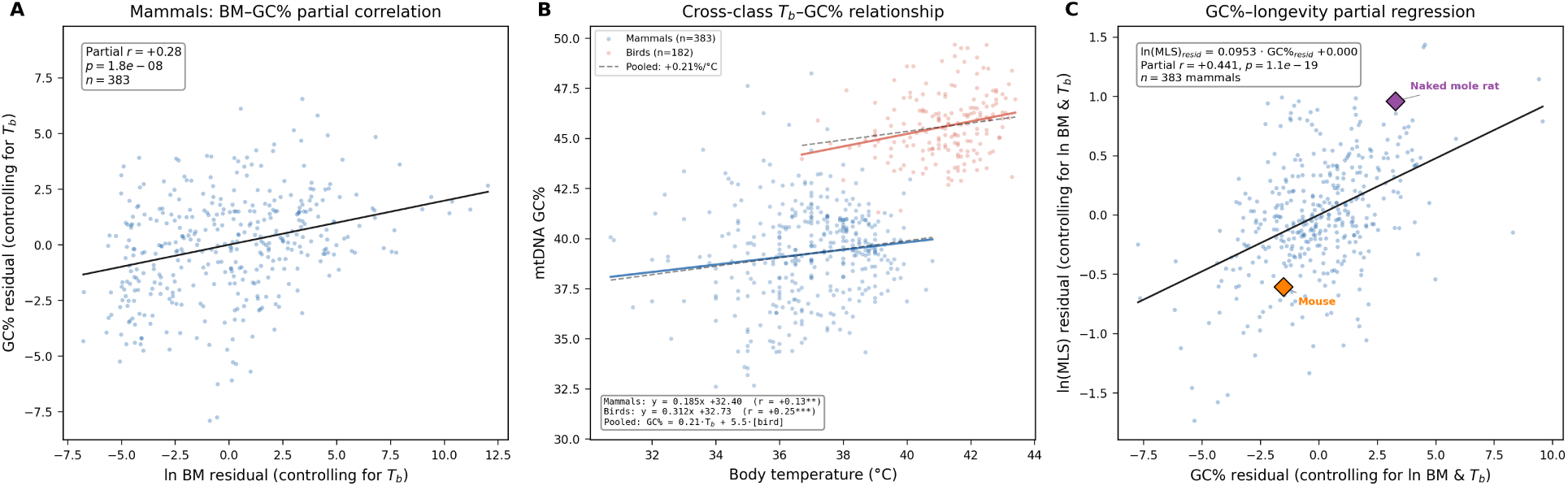
Pace–Stability co-evolution across endotherms. **(A)** Partial regression of ln BM residuals versus GC% residuals after controlling for Tb in mammals (n=379). The positive partial correlation (r=+0.30, p<10*^−^*^6^) indicates that larger mammals maintain higher mtDNA GC% independently of temperature, consistent with Scope–Stability co-selection. **(B)** Cross-class relationship between Tb and mtDNA GC%. Mammals (n=379): GC%=0.225 Tb+30.90 (r=+0.16, p<0.01); birds (n=182): GC%=0.312 Tb+32.73 (r=+0.25, p<0.001). Pooled model controlling for class: +0.21%/° C with a +5.5% avian offset, supporting a universal thermal adaptation mechanism. **(C)** Partial regression of GC% versus lnMLS after removing the effects of both ln BM and Tb in mammals (n=379). The Stability–longevity association remains strong after full confound control (partial r=+0.485, p=1.1×10^-23^; slope =0.105 ln MLS per GC percentage point). Diamond markers: NMR (purple) is a strong positive outlier; mouse (orange) falls below the regression line. This partial effect of GC% matches the SSP *β* coefficient in the full model, confirming the independent contribution of Stability to longevity.

Cross-class comparison reveals the universal thermal signal. In birds, where Tb–body mass confounding is weaker, the temperature–GC% relationship persists after controlling for body mass (*β*=+0.24% per °C, p=0.014). Importantly, mammalian and avian Tb–GC% slopes are statistically indistinguishable (interaction p=0.47), supporting a common thermal adaptation mechanism. Pooling both endotherm classes yields a robust universal estimate: each 1 °C increase in body temperature associates with +0.23% higher mtDNA GC content (p<0.001; Fig. 3B). The class-level difference—birds maintain ∼5.4% higher GC% than mammals at the same body temperature— may reflect their longer evolutionary exposure to elevated thermal stress or additional avian-specific constraints.

H-strand adenine content drives the Stability–longevity relationship. Examination of individual base correlations reveals that H-strand adenine (reported as L-strand thymine in sequence databases) shows the strongest association with mammalian longevity (r=-0.46, p<10^-20^; Fig. 3C). Long-lived mammals maintain low H-strand adenine content; short-lived species retain high adenine. This pattern implicates adenine deamination (A → hypoxanthine → G) as the primary mutagenic pathway, consistent with the known vulnerability of unpaired adenine to oxidative and hydrolytic attack during single-strand exposure. The GC%–longevity correlation thus reflects selection against H-strand adenine—the deamination substrate—rather than for cytosine retention per se.

Saturated Stability in birds explains their absent GC%–longevity correlation. Despite higher thermal pressure than mammals (Tb≈41 °C vs 37 °C), birds show no GC%–MLS correlation (r=-0.01, p=0.85; n=182). This apparent paradox resolves upon examining H-strand adenine: avian H-A content averages 23.9% with compressed variance (SD =1.0%), compared to mammalian 27.7% (SD =1.8%). The avian 95th percentile for H-A% (25.6%) approximates the mammalian 25th percentile (26.6%)—even high-adenine birds sit at the mammalian low end. Birds have driven H-strand adenine to its functional floor, exhausting variance for further optimization. Selection cannot improve what is already minimized.

#### Summary

The Pace–Stability co-evolution proposed by MSTA operates across endotherms but is obscured within mammals by body-mass confounding and within birds by compositional saturation. The universal Tb–GC% relationship (+0.23% per °C) emerges clearly only when classes are pooled, providing cross-taxonomic support for thermal adaptation. The mechanistic driver is minimization of H-strand adenine—the thermally vulnerable deamination substrate—explaining both the positive GC%–longevity correlation in mammals (where variance remains) and its absence in birds (where optimization is complete).

Notably, the GC%–longevity slope is steepest not for any protein-coding gene but for the 12S rRNA (0.242 lnMLS per GC percentage point, n=107 mammals), which encodes the small ribosomal subunit responsible for translational decoding fidelity. Because a single deamination lesion in a decoding-critical rRNA position can corrupt translation of all 13 mtDNA-encoded proteins for that ribosome’s remaining lifetime, the per-nucleotide leverage of rRNA protection exceeds that of any individual coding gene. The Stability pillar thus operates at least in part through preservation of translational fidelity — a multiplicative layer that compounds the coding-gene-specific effects discussed below (Section 2.6).

### 2.3 Exceptional longevity via SSP axis optimization: volant species and the naked mole rat

The SSP framework resolves apparent paradoxes in lineages with extreme lifespans.

#### Volant species: decomposing the bird–mammal longevity gap

At equivalent body mass, birds live ∼63% longer than mammals (*β*=+0.49 lnMLS, SE 0.05, p<10*^−^*^20^; n=561 pooled). The advantage is robustly real, but its attribution across the SSP axes is not uniquely identified.

Birds face elevated thermal pressure (mean Tb=41.2 °C vs mammalian 37.0 °C), which should accelerate mtDNA damage. However, they maintain substantially higher mtDNA GC% (45.6% vs 39.2%), providing enhanced thermal stability. Because these two axes co-vary across the contrast (birds are both hotter and more GC-rich), Table 1 reports how the avian residual moves with model specification rather than a single decomposition:

**Table 1.** The bird–mammal longevity gap is robust but its SSP partition is not identified. Coefficient on an Aves indicator in regressions of lnMLS on ln BM plus the listed predictor(s), pooled mammals + birds (n=561). The matched-mass advantage (+0.49) is highly significant, but the residual after attribution ranges from non-significant to +0.83 depending on whether GC% or Tb is conditioned on, because the two predictors are collinear across the contrast (birds are both +6.4 GC% points and +4.2 °C hotter). Adding GC% alone renders the avian advantage non-significant—Stability is statistically sufficient.

| Model (added to ln BM) | Bird residual (lnMLS) | *t* | Interpretation |
| --- | --- | --- | --- |
| — (body mass only) | +0.49 | 10.0 | Raw matched-mass advantage |
| + GC% | +0.02 | 0.3 | Stability alone absorbs the gap (n.s.) |
| + Tb | +0.83 | 11.3 | Conditioning on Pace inflates the residual |
| + GC% + Tb | +0.36 | 4.1 | Joint model: residual partly remains |

*Joint-model coefficients on this set are +0.078 lnMLS per GC% point and −0.092 per °C; applied to the class means they give a Stability gain of +0.50 ( $\approx+65\%$ MLS) and a Pace penalty of −0.39 ( $\approx-32\%$ ), net +0.11, with a remaining class residual of +0.36. Because that residual swings from +0.02 to +0.83 across specifications, it is treated as unexplained class/volant variance, not as a measurement of Scope/conductance reserve (§3.1).*

**Table 2.** Two-criterion test for the GC%–MLS correlation across vertebrate classes. The correlation is strongest where both thermal pressure and residual H-strand adenine variance are present; it attenuates where either condition is absent (birds at the A_H floor; ectotherms at low Tb).

| Class | Thermal pressure | GC% variance | H-A% status | GC%–MLS $r$ |
| --- | --- | --- | --- | --- |
| Mammals | High ($37^\circ\text{C}$) | High (SD =2.3) | 28% | +0.48* |
| Birds | Very high ($41^\circ\text{C}$) | Low (SD =1.6) | 24% | -0.01 |
| Fish | Low ($16^\circ\text{C}$) | Low (SD =1.8) | 26% | -0.16* |

The dominant message is that Stability is statistically sufficient: GC% alone reduces the avian advantage to +0.02 lnMLS (t=0.3), so birds’ elevated mtDNA GC% accounts for their matched-mass longevity before Pace is even considered. In the joint model the Stability gain (+0.50) largely offsets the Pace penalty (−0.39) for a small net mtDNA effect (+0.11), leaving a class residual of +0.36 that is not uniquely attributable.

Earlier framings assigned this residual to expanded avian aerobic Scope; that attribution is not supported here. Aerobic scope is not measured for the avian dataset, and the residual is specification-dependent (Table 1); moreover, whole-organism aerobic scope is a locomotor-dominated proxy for cellular reserve rather than a direct measure of it (§3.10). The avian residual is therefore best read as unexplained class/volant variance—candidate contributors including reduced extrinsic mortality, life-history remodeling, flight-related physiology, and unmeasured conductance architecture—rather than as a quantified Scope term.

Bats follow similar logic but employ an additional strategy: torpor and hibernation lower effective annualized Pace by reducing cumulative thermal exposure, achieving longevity extension through this Pace reduction, a route independent of aerobic capacity (bat SSP residual: +0.37 ± 0.53 lnMLS; Fig. 4B)^21^. The anomaly is concentrated in the hibernating families (Vespertilionidae and Rhinolophidae), which carry lower mtDNA GC% (∼37% versus ∼41% in non-hibernating bats) yet live longest—months of torpor lower their effective thermal dose well below the euthermic Tb the SSP formula uses, a Pace discount the model does not capture.

**Figure 4.**
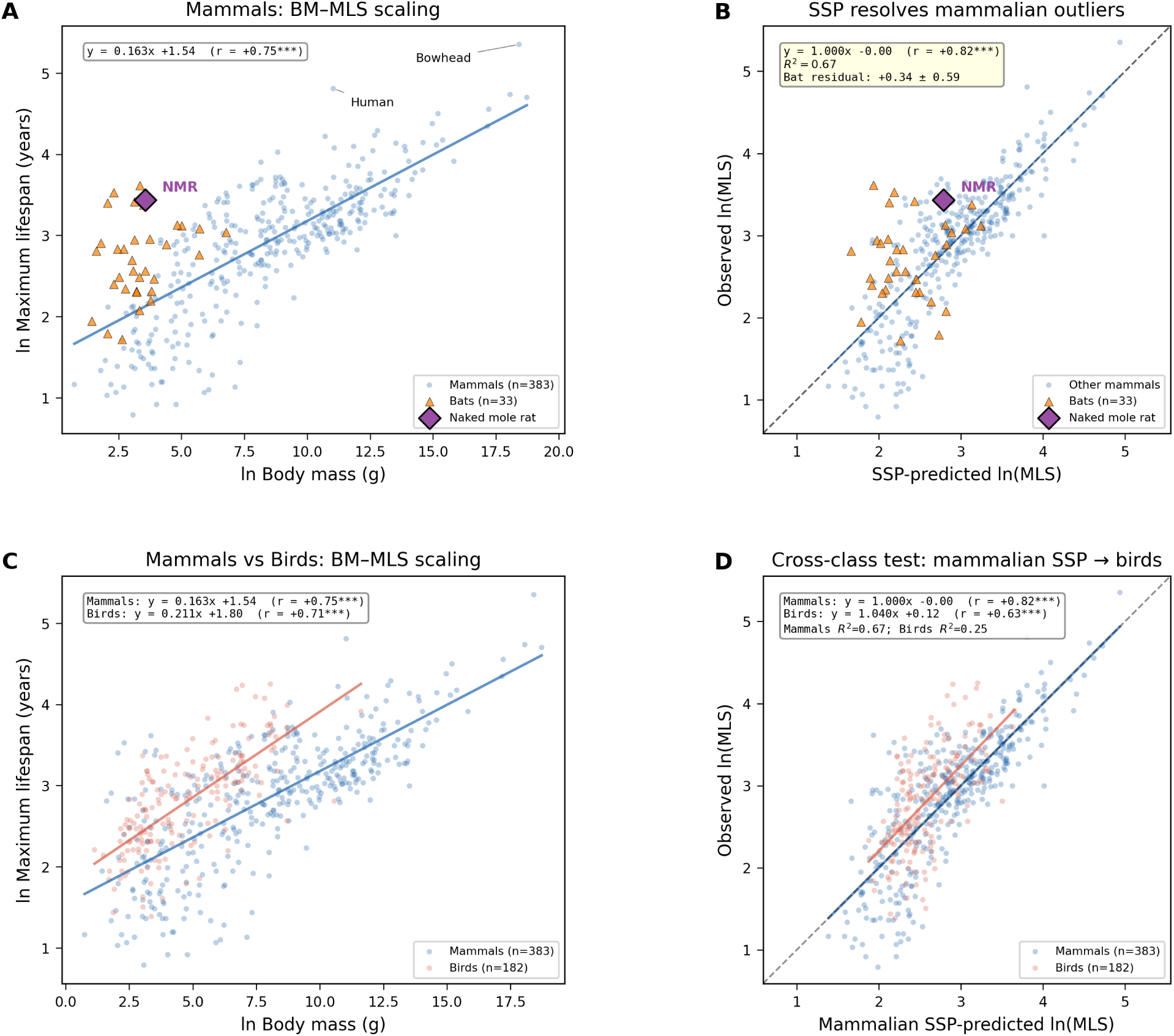
SSP resolves mammalian outliers and extends to birds. **(A)** Body-mass–lifespan scaling in mammals (n=379). lnMLS=0.161 lnBM+1.57 (r=+0.75, p<0.001). Bats (triangles, n=31; filled = hibernating Vespertilionidae/Rhinolophidae, n=11; open = non-hibernating, n=20) and NMR (purple diamond) are conspicuous positive outliers from the allometric trend (bat residual +0.64±0.45). Human (red diamond) and bowhead whale (blue diamond), like NMR, are marked in both panels A and B. **(B)** SSP predicted versus observed lnMLS for the same 379 mammals. The regression of observed on predicted values yields y=1.000x-0.00 (r=+0.83, R^2^=0.69). Bat residual: +0.37±0.53 (mean ± SD); adding GC% and Tb reduces the bat excess from +0.64 (panel A) to +0.37 ln units — from ∼1.9× to ∼1.45× longer-lived than predicted — so SSP substantially but not fully absorbs the anomaly. **(C)** Comparison of body-mass–lifespan allometry between mammals (n=379) and birds (n=182). Mammals: slope =0.161, r=+0.75; birds: slope =0.211, r=+0.71. The steeper avian slope reflects elevated Stability (GC%) together with additional class-specific variance not captured by the mammalian SSP fit. **(D)** Cross-class validation: the mammalian SSP model (coefficients from panel B) applied to predict avian lifespans. Mammals: r=+0.83, R^2^=0.69; birds predicted by mammalian model: slope =1.03, r=+0.61, R^2^=0.37. The mammalian SSP model retains significant predictive power for birds, though the reduced R^2^ suggests additional class-specific variance consistent with the +5.5% GC offset (Fig. 3B) and unmeasured avian-specific factors.

#### Naked mole rat (Heterocephalus glaber)

This small rodent achieves rodent-record lifespan via complementary axis shifts. Relative to similar-sized mammals, it maintains low resting Tb (∼32 °C), exponentially slowing hydrolytic deamination (Pace suppression), and profoundly depressed BMR (∼75% below allometric expectation), which would expand effective factorial Scope if maximal aerobic output is preserved—a point requiring direct measurement. Combined with elevated GC% (high Stability), the NMR occupies a rare SSP configuration: low Pace + high Stability + high effective Scope (Fig. 4)^22,23^.

These cases illustrate evolutionary levers for exceptional lifespan—raising Scope (flight, relative BMR suppression), raising Stability (GC enrichment), and/or lowering Pace (hypothermia, torpor)—all generating coherent SSP balances that extend MLS without violating thermodynamic limits.

### 2.4 Ectotherm comparisons: Pace dominance at temperature extremes

#### 2.4.1 Fish achieve mammalian longevity despite ∼30-fold lower absolute metabolic power

If absolute aerobic capacity determined longevity, cold ectotherms should live shorter than endotherms of equivalent mass. At the same body mass, ectothermic fish have ∼24-fold lower resting metabolic rates and ∼30-fold lower maximal metabolic rates than endotherms^24^. Yet factorial aerobic scope—the ratio MMR/BMR—is broadly similar across vertebrate classes (∼5–8× in both groups). Contrary to an absolute-power penalty prediction, fishes live as long as or slightly longer than mammals at equivalent body mass (*β*=+0.06 lnMLS, p=0.07; n=1,438 species; Fig. 5A). Remarkably, some fish achieve extraordinary longevity: the rougheye rockfish (*Sebastes aleutianus*; deep-water Tb ≈ 5°C) lives 205 years at only 0.5 kg body mass—approximately 15-fold longer than predicted for a mammal of equivalent size. Lake sturgeon (*Acipenser fulvescens*; Tb ≈ 10–15°C) reach 152 years at 70 kg; orange roughy (*Hoplostethus atlanticus*; deep-water Tb ≈ 4°C) achieve 149 years at 3.8 kg.

**Figure 5.**
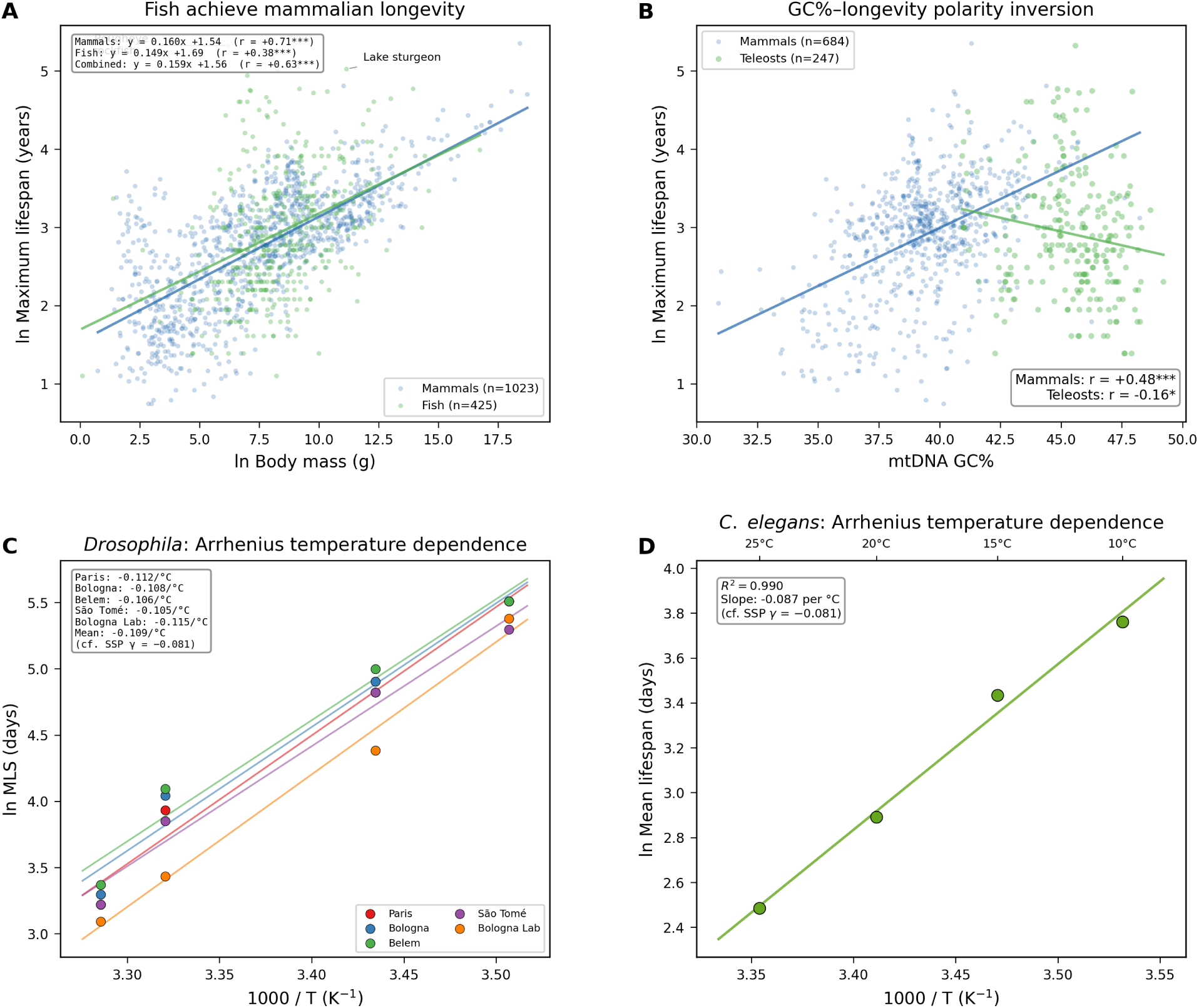
Ectotherm comparisons validate Pace dominance at temperature extremes. **(A)** Body-mass–lifespan scaling in mammals (n=1,018) and fish (n=420, including teleosts, chondrichthyans, chondrosteans, and holosteans). Mammals: y=0.159x+1.55 (r=+0.71); fish: y=0.144x+1.73 (r=+0.37); combined: y=0.158x+1.57 (r=+0.63). Despite ∼30-fold lower absolute metabolic power, fish achieve mammalian-grade longevity at equivalent body mass, consistent with MSTA’s prediction that factorial aerobic scope—not absolute metabolic rate—determines lifespan. Lake sturgeon (*Acipenser fulvescens*) and rougheye rockfish (*Sebastes aleutianus*) are annotated as extreme positive outliers. **(B)** GC%–longevity polarity inversion. In mammals (n=679), GC% correlates positively with lnMLS (r=+0.50, p<0.001), whereas in teleost fish (n=247) the correlation is weakly negative (r=-0.16, p=0.01). This inversion is predicted by MSTA: at low ambient temperature (Tb≈16 ° C), thermal deamination is negligible, eliminating the selective premium for GC enrichment while its metabolic costs persist. **(C)** Arrhenius temperature dependence of lifespan in *Drosophila melanogaster*. Five geographically distinct strains (Paris, Bologna, Belem, São Tomé, Bologna Lab)^26^ show near-identical ln(MLS)-vs-1000/T slopes (range: -0.105 to -0.115 ° C-1; mean =-0.109 ° C-1), bracketing the SSP Pace coefficient (−*γ*=−0.091). The convergent slopes across strains acclimated to tropical and temperate regimes demonstrate that temperature acts through a universal thermodynamic mechanism rather than genotype-specific adaptation. **(D)** Arrhenius temperature dependence of mean lifespan in *Caenorhabditis elegans* (10–25 ° C)^28^. The slope (-0.087 ° C-1, R^2^=0.990) closely matches the SSP −*γ*=−0.091, extending kinetic validation to a second ectotherm phylum (Nematoda) and confirming that the Pace coefficient reflects a phylogenetically conserved Arrhenius-like thermal dependence of the lesions that limit lifespan.

This pattern resolves within the MSTA framework: longevity depends on the ratio of regenerative capacity to damage rate, not absolute metabolic throughput. Fish have ∼30-fold lower absolute metabolic power but ∼4-fold slower thermally driven damage accumulation (Arrhenius kinetics at Tb≈16 °C vs 37 °C, Q10≈2). When damage is sufficiently slow, even modest repair capacity suffices. The fish comparison validates that Pace reduction can compensate for low absolute Scope, provided the ratio of repair to damage remains favorable^25^.

This finding has important implications for the Strehler–Mildvan formulation (Section 2.5): it is factorial scope (the dimensionless ratio MMR/BMR), not absolute aerobic power, that determines the “vitality” buffer—initial reserve capacity relative to average challenge—against mortality. Fish and mammals share similar factorial scope (∼5–8×) and achieve similar body-mass-adjusted longevity—despite operating at vastly different absolute metabolic scales.

#### 2.4.2 Thermal removal collapses the Stability–longevity signal

MSTA predicts that GC% enrichment confers longevity benefit primarily by mitigating temperature-accelerated deamination during single-strand exposure. In fish (mean Tb≈16 °C), this selective pressure should weaken substantially.

Consistent with prediction, the GC%–MLS correlation in teleosts is weakly negative (r=-0.16, p=0.01; n=247)—a polarity inversion from the strong positive correlation in mammals (r=+0.48; Fig. 5B). Notably, fish maintain high GC% (∼45%, similar to birds) yet derive no longevity benefit—and may incur a slight cost—from this composition. This supports the conditional Pace–Stability interaction: at low temperatures, the “stability premium” for high GC% disappears because damage accumulation is already sufficiently slow. The weak negative correlation may reflect metabolic or structural costs of GC-rich genomes that are outweighed by stability benefits only under thermal pressure.

##### Two criteria for GC%–longevity correlation

Cross-class comparison reveals that the GC%–MLS correlation is strongest when both conditions are met: (1) sufficient thermal pressure to drive deamination, and (2) sufficient compositional variance for selection to operate.

Birds face intense thermal pressure but have already minimized H-strand adenine to ∼24% (the apparent functional floor)—leaving no variance for selection. Fish face minimal thermal pressure—making GC% optimization unnecessary despite available variance. Only mammals satisfy both criteria, explaining why the GC%–MLS correlation emerges strongly in this class alone.

#### 2.4.3 Kinetic validation of Pace: interspecies coefficient matches within-species thermal manipulation

Endotherms fixed Tb within species, precluding direct experimental tests of thermal effects on MLS. Ectotherms enable manipulation: rearing temperature varies while genotype and morphology remain constant, isolating Pace’s causal impact.

We tested whether the mammalian SSP-derived temperature coefficient (−*γ* = −0.091 °C*^−^*^1^) matches the observed temperature–lifespan slope in ectotherms reared across thermal regimes. Five independent datasets, spanning Insecta, Branchiopoda, Nematoda, and Actinopterygii, yield near-identical Arrhenius slopes for mean lifespan as a function of rearing temperature (Table 3, Fig. 5C–D).

**Table 3.** Within-species temperature–lifespan slopes in five ectotherms. All five slopes bracket the mammalian SSP Pace coefficient (−*γ*_SSP = −0.091 °C*^−^*^1^).

| Species (clade) | T range<br>($^{\circ}\text{C}$) | Cohort | d ln(MLS)/dT<br>($^{\circ}\text{C}^{-1}$) | R<sup>2</sup> | Ref |
| --- | --- | --- | --- | --- | --- |
| *Drosophila melanogaster*<br>(Insecta) | — | 5 geographic<br>strains | −0.105 to −0.115<br>(mean −0.109) | — | 26 |
| *Musca domestica*<br>(Insecta) | 20–35 | n = 150 ♀ per<br>T | −0.094 | 0.9998 | 27 |
| *Caenorhabditis elegans*<br>(Nematoda) | 10–25 | — | −0.087 | 0.990 | 28 |
| *Daphnia magna*<br>(Branchiopoda) | 5–30 | 4 clonal lines | −0.093* | — | 29 |
| *Nothobranchius rachovii*<br>(Actinopterygii) | 20–30 | n = 300 ♂ | −0.091 | 0.999 | 30 |

The five within-species slopes bracket the mammalian −*γ*SSP = −0.091 tightly, and the mammalian 95% confidence interval encompasses every ectotherm point estimate. This is quantitative consistency between the comparative mammalian regression and five independent experimental manipulations performed in different species, different laboratories, and different decades — evidence that the Pace coefficient reflects a universal thermodynamic mechanism rather than organism-specific metabolic programming.

Convergence is specific to the slope, not the intercept. Absolute lifespans at matched Tb differ by orders of magnitude across mammals, flies, and nematodes — Scope and Stability, which set the intercept, are phylum-specific. What converges is the *sensitivity* of lifespan to temperature, dln(MLS)/dTb, and this is precisely what MSTA’s axis decomposition predicts: Pace is the only SSP axis whose effect flows through Arrhenius chemistry, so it is the only axis for which cross-phylum invariance is expected. Consistent with this, the SSP temperature coefficient (*γ* = +0.091 °C*^−^*^1^) corresponds to an observed activation energy of Ea≈73 kJ/mol (0.75 eV). This value sits in the center of the canonical biochemical window (60–85 kJ/mol) and is consistent with the central parameter of the Metabolic Theory of Ecology (MTE), which predicts a fundamental biochemical Ea of ∼0.63 eV (∼61 kJ/mol) across taxa. The C. elegans slope yields Ea ≈ 61 kJ/mol, matching the MTE prediction precisely; N. rachovii and M. domestica yield Ea ≈ 67–68 kJ/mol; D. melanogaster and D. magna 66–82 kJ/mol. Every value falls within the canonical biochemical window. The five cross-phylum ectotherm Arrhenius slopes, the mammalian SSP coefficient, and the MTE universal activation energy therefore converge on a single thermal-chemistry parameter — strong evidence that the Pace term captures a phylogenetically invariant property of the chemistry that limits lifespan (Supplementary Note S1.1).

### 2.5 Linking SSP to Gompertz mortality dynamics

Strehler and Mildvan showed^31^ that Gompertzian mortality emerges when physiological reserve declines against stochastically distributed environmental challenges. Their theory defines a dimensionless “vitality ratio” (V0/*ε*D)—initial reserve capacity relative to average demand—that determines mortality kinetics. MSTA identifies this ratio with factorial aerobic scope:

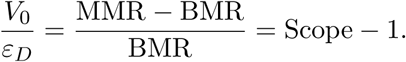

Strehler and Mildvan derived V0/*ε*D≈7–11 from Gompertz parameters across 32 national mortality tables, and validated this against physiological measurements (cardiac index: 7.9–18; heat output ratio: 6.6–13.7). Modern doubly labeled water studies confirm human factorial scope of 8–10× during sustained maximal exertion (Ruby et al., 2015; Thurber et al., 2019), yielding Scope-1≈7–9—quantitative agreement across demographic, classical physiological, and modern calorimetric methods.

#### Longitudinal validation

The 47-year SPAF cohort^32^ reveals patterns predicted by stochastic scope erosion. Mass-specific VO_2_max declined exponentially from 42 mL/kg/min at age 34 to 26 mL/kg/min at age 63 (∼1.3%/year at the median). Since basal metabolic rate declines more slowly (∼0.5%/year), factorial scope contracts at approximately 1%/year—matching Strehler–Mildvan’s demographically derived vitality erosion rate.

Critically, relative variability doubles with age (IQR/median: 0.17 at the youngest cohort ages → 0.38 by age 63): the population “spreads out” as individuals erode at different rates. This expanding variance is characteristic of stochastic damage accumulation rather than programmed decline. Mortality emerges from the lower tail: by age 63, the lowest-decile individuals (VO_2_max≈11.9 mL/kg/min) have already fallen below the ∼18 mL/kg/min threshold associated with loss of functional independence^33^, retaining only ∼3.4× resting metabolic rate. These tail-crossing dynamics—where the lower edge of the reserve distribution increasingly overlaps with the challenge distribution—are precisely what generates Gompertzian mortality acceleration in the Strehler–Mildvan framework.

This decline has a circuit origin in the concurrent erosion of three power-delivery parameters. As impedance accumulates, the effective driving voltage falls (V_redox drops from ∼1.14 V toward ∼0.77 V as electron entry shifts from the high-voltage NADH/respirasome route to the CoQ route), the proton stoichiometry falls (n_H from ∼10 toward ∼6 H^+^ per 2e*^−^*), and the internal resistance R_total rises. Because deliverable power scales as the product P ∝ V_redox × n_H / R_total, modest concurrent changes compound—a ∼20% loss in each variable removes roughly two-thirds of deliverable power—so reserve contracts faster than any single parameter, approximately exponentially: V(t) ≈ V_0_·exp(−kt), with V_0_ the factorial aerobic scope (MMR/BMR − 1) and k the composite erosion rate. An exponentially declining reserve crossing a stochastic challenge distribution yields the Gompertz hazard h(t) = h_0_·exp(*α*t), recovering Strehler–Mildvan mortality acceleration from first principles; the full circuit derivation is given in Supplementary Note S3.1a. Crucially, the Scope formulation depends on the dimensionless ratio MMR/BMR, not absolute metabolic power. This explains why fish achieve mammalian longevity despite ∼30-fold lower absolute metabolic throughput (Section 2.4.1): with similar factorial scope (∼8×) and dramatically reduced Pace, the damage-to-repair ratio remains favorable.

The SSP law is not merely correlative. Its three terms map onto a causal structure: Scope (indexed by body mass) sets the initial reserve; Stability (indexed by GC%) and Pace (indexed by Tb) together set the rate at which somatic mtDNA lesions accumulate — GC-rich heavy strands presenting fewer deamination-prone bases, and lower body temperature slowing the Arrhenius kinetics of their loss. If this interpretation is correct, then the pattern of functional decline should be predictable from the architecture of the machinery those lesions damage. Thirteen mtDNA-encoded proteins constitute the core subunits of Complexes I, III, IV, and ATP synthase; cumulative mutations in these subunits should manifest as rising internal resistance in the electron transport chain. To track the expected consequences of this progressive impedance growth, we model the respiratory chain as three coupled circuits: electron, proton, and carbon.

### 2.6 Molecular basis of Scope erosion: circuit-derived predictions of metabolic impedance

The SSP law can be written as an impedance-traversal equation. Let Rint(t)denote the effective internal resistance of the OXPHOS network, and define normalized impedance as:

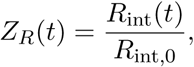

where *R*_int_,_0_ is the young-adult baseline and *R*_int_,_crit_ is the critical resistance at which regenerative biosynthetic throughput falls below maintenance demand. The dimensionless interval

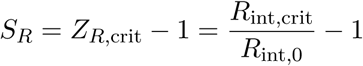

defines the available impedance scope: the normalized distance the electron, proton, and carbon circuits can traverse before failure becomes irreversible. For scaling purposes, *S_R_* is proportional to the multiplicative dynamic range *R*_int_,_crit_*/R*_int_,_0_. The rate

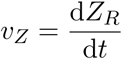

defines the velocity at which this scope is consumed. Maximum lifespan is therefore approximated by:

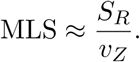

This expression separates lifespan into a **capacity term** and a **rate term**. Body mass enters primarily through the capacity term. Allometric divergence between basal and maximal metabolic rates produces factorial metabolic scope,

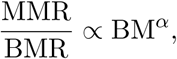

so larger animals carry greater respiratory-chain headroom relative to baseline demand and can tolerate a larger normalized rise in *R*_int_ before reaching *R*_int_,_crit_. Thus:

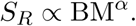

Mitochondrial base composition and body temperature enter primarily through the rate term. During asymmetric mtDNA replication, single-stranded exposure of the H-strand creates a vulnerable substrate pool for hydrolytic deamination. The lesion-generation rate can be written schematically as:

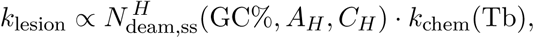

where 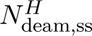 represents the burden of deamination-prone single-stranded bases, including adenine and cytosine, and *k*_chem_(Tb) follows Arrhenius temperature dependence. Higher mtDNA GC content, as used in the SSP regression, indexes a lower vulnerable-base burden, while lower body temperature slows the chemical rate constant. Therefore:

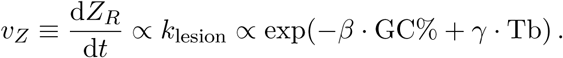

Substituting the scope and rate terms into the traversal equation gives:

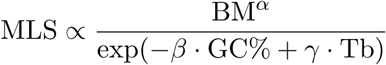

and therefore:

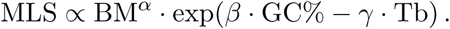

Taking logarithms recovers the SSP form:

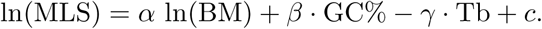

Thus, the SSP regression is a reduced physical form of the MSTA mechanism. The body-mass coefficient *α* captures allometric expansion of impedance scope; the GC% coefficient *β* captures sequence-dependent reduction of mtDNA lesion velocity; and the temperature coefficient *γ* captures thermal acceleration of the same lesion chemistry. The structural form of the SSP law is therefore mechanistically informative: BM behaves as a static scope determinant, whereas GC% and Tb behave as rate determinants integrated across the lifespan. The remainder of this section derives how progressive increases in *R*_int_ are translated by the electron, proton, and carbon circuits into the ordered metabolic failures of aging.

#### 2.6.1 Circuit architecture and output law

Aging in this framework is a single lesion read out three ways. The pathology is one variable—rising internal resistance (R_int) in the electron-transport chain, the cumulative imprint of mtDNA-linked conductance loss—but its consequences are partitioned across three coupled circuits, each gating a different currency under a different engagement mode (Fig. 6). Because the lesion sits in the charge pump between a source and a load, its effect propagates in both directions: upstream, throttling the carbon circuit that feeds it, and downstream, starving the proton load it charges. Failure is therefore directional—metabolites accumulate upstream of throttled gates while downstream products decline—and we trace it in the direction current flows, from source to load: the carbon circuit (substrate oxidation, the source current), the electron circuit (the charge pump, where R_int accumulates), and the proton circuit (the load drawn against Δ*p*).

**Figure 6.**
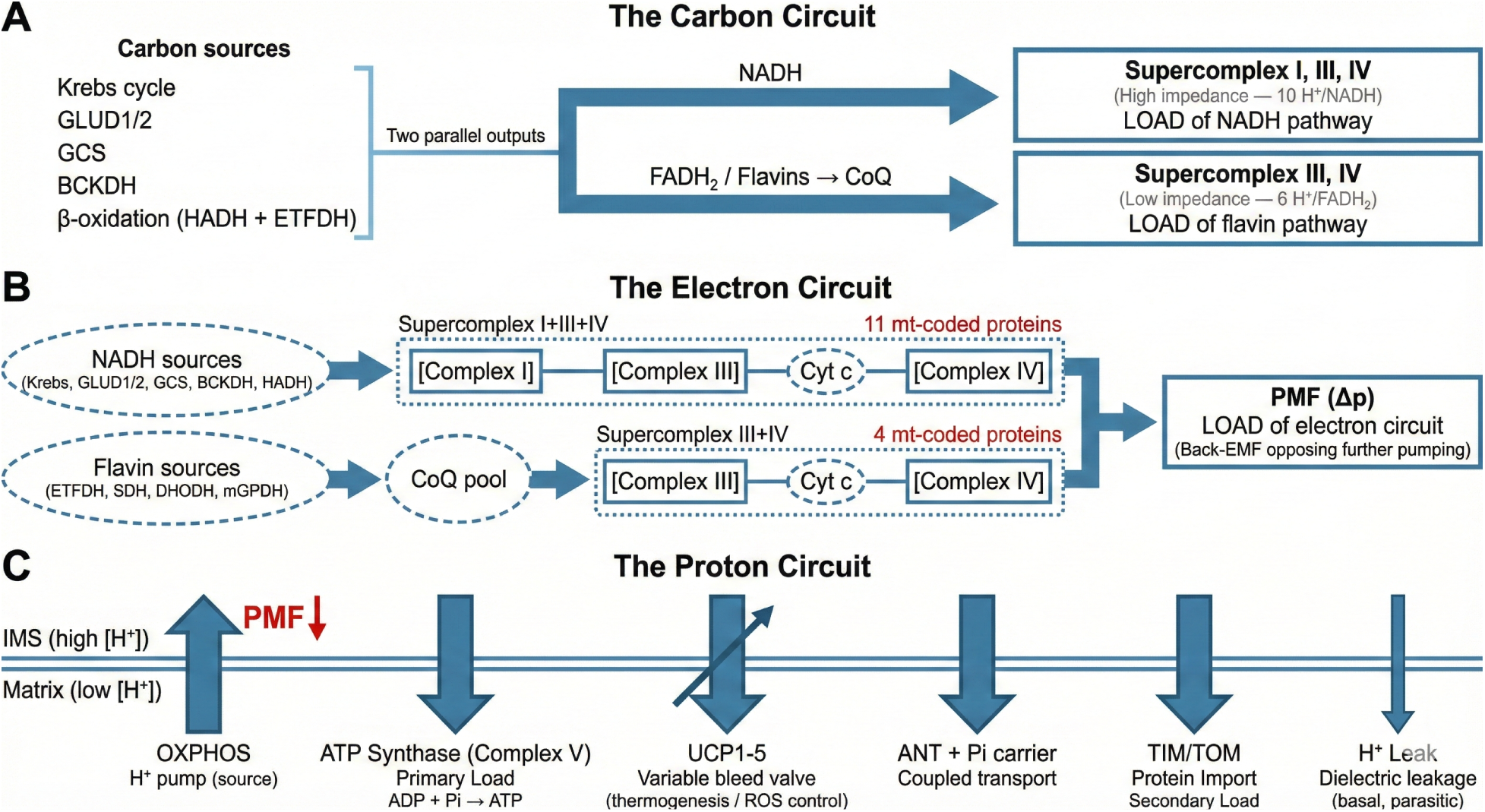
Three nested circuits of mitochondrial energy transduction and the origin of selective NADH vulnerability. **(A)** The carbon circuit. NAD^+^-dependent dehydrogenases (Krebs cycle, GLUD1/2, GCS, BCKDH, *β*-oxidation) generate two parallel outputs: NADH, which must pass through Supercomplex I+III+IV (11 mtDNA-encoded subunits), and FADH_2_/flavins (from SDH, ETFDH, DHODH, mGPDH), which reduce the CoQ pool and enter only through Supercomplex III+IV (4 mtDNA-encoded subunits). **(B)** The electron circuit. The NADH path traverses 11 mtDNA-encoded subunits in series versus 4 for the flavin/CoQ path; both pathways accumulate impedance, but NADH back-pressure dominates the aging phenotype because Complex I is the sole matrix NADH exit and its downstream sensors (DLD/E3, IDH3, MDH2) operate near thermodynamic equilibrium, reporting constitutively in every cell at rest. Flavin-linked pathways that bypass Complex I continue to reduce CoQ and sustain the PMF; their bottleneck becomes rate-limiting only under proliferative or catabolic demand. **(C)** The proton circuit. The PMF generated by OXPHOS is itself a load — back-EMF opposing further proton pumping. Lowering PMF (e.g., via uncoupling) relieves upstream electron flow but reduces ATP synthesis, the primary load of the proton circuit. Secondary loads include the ANT/P_i_ carrier, ΔΨ-dependent protein import (TIM23/PAM, after TOM entry), regulated uncoupling (UCP1–5), and passive proton leak.

In the **carbon circuit**, the Krebs cycle is the source: it draws carbon substrate (entering via PDH and anaplerotic feeds) and delivers two outputs—the reducing equivalents (NADH, FADH_2_) that drive the electron circuit downstream, and the cataplerotic precursors that supply biosynthesis (citrate → lipogenesis, *α*-KG → 2-oxoglutarate-dependent oxygenases, OAA → aspartate and nucleotides, succinyl-CoA → heme). Its work is therefore dual—powering the chain and renewing the cell—and we define *regenerative scope* as the spare cataplerotic throughput sustainable above maintenance turnover, the cellular correlate of the factorial aerobic scope measured at the organismal level (§2.1). The NAD^+^-dependent dehydrogenases^36^—IDH3, *α*KGDH, MDH2, and the five DLD-gated complexes—are its gating resistors, their conductance set by matrix NAD^+^ availability (Cambronne, X. A. & Kraus, W. L)^37^, coupling the circuit to the electron circuit downstream. Because matrix NADH has a single oxidative outlet—Complex I—the circuit is constitutively engaged: every resting cell reports rising R_int as an elevated NADH/NAD^+^ ratio that dims these near-equilibrium dehydrogenases before any demand is imposed.

In the **electron circuit**, the impedance-bearing redox charge pump carrying this current—an nDNA-encoded electron conductor coupled to an mtDNA-encoded proton-pumping core, so that mtDNA-linked damage raises R_int in the pump while the conducting framework is spared—two parallel routes converge on a shared terminal oxidase. The respirasome (I–III_2_–IV) spans the NADH-to-O_2_ redox potential (V_redox ≈ 1.14 V); the smaller III_2_–IV supercomplex spans CoQH_2_-to-O_2_ (∼0.77 V), fed by Complex II, ETFDH, DHODH, and mGPDH^34,35^. Both routes terminate in no-bypass series nodes: Complex I is the sole matrix NADH outlet—mammals lack the alternative NADH dehydrogenase (NDH-2, e.g. Ndi1) that in plants, fungi, and protists routes NADH electrons around it to CoQ—and Complex III is the sole CoQ-pool oxidase, mammals likewise lacking the alternative oxidase (AOX) that branches from the CoQ pool to vent reduced quinone directly to O_2_ in those lineages, so electrons cannot route around either lesion. Electron flux (I_e) is the circuit current; lesions in mtDNA-encoded subunits act as series resistors, raising R_int with Complexes I, III, and IV in line. The redox driving force is thermodynamically fixed, so as R_int rises the current—and with it the rate of NAD^+^ and oxidized-CoQ regeneration on which the carbon circuit depends—falls. Because the flavin/CoQ route bypasses Complex I, it supplies the alternative entry the NADH arm lacks; its bottleneck is exposed only under high demand, making the CoQ gate demand-revealed rather than constitutive.

In the **proton circuit**, the load, proton pumping by the electron circuit charges the inner-membrane capacitance, generating the proton-motive force (Δ*p*, dominated by Δ*ψ*)^13^. ATP synthase and proton-leak pathways discharge it. ATP synthase occupies a dual role: it is a **load** (consuming Δ*p* to synthesize ATP) but simultaneously a **relief valve** for the electron circuit—by dissipating Δ*p* it lowers the back-EMF opposing further pumping and permits greater electron flux, as demonstrated during State 3 (ADP-stimulated) respiration. Because deliverable power to a load of fixed resistance scales as Δ*ψ*^2^, the proton circuit sheds its highest-threshold work first while basal ATP is defended—the threshold logic that makes Δ*ψ* the third gate.

The three circuits are coupled, and that coupling is what turns a single charge-pump lesion into a system-wide syndrome. Rising R_int slows NAD^+^ regeneration, raising the effective resistance of the carbon circuit’s gating dehydrogenases and collapsing biosynthetic throughput—the electron-to-carbon link that dominates the aging phenotype. The same impedance reduces proton pumping and lowers Δ*ψ*, eroding the work the proton circuit can deliver. A third coupling runs the other way: nicotinamide nucleotide transhydrogenase (NNT) uses Δ*p* to drive NADH + NADP^+^ → NAD^+^ + NADPH, transmitting reductive pressure from the electron circuit into the NADPH pool (Rydström, J.)^38^ and tying proton-circuit voltage to carbon-circuit recovery. One further coupling is acute rather than cumulative, and we bracket it here: Δ*ψ* also gates electrogenic Ca^2+^ entry through the uniporter, and matrix Ca^2+^ activates the carbon circuit’s gating dehydrogenases—pyruvate dehydrogenase, NAD^+^-linked isocitrate dehydrogenase, and *α*-ketoglutarate dehydrogenase—matching flux to demand within seconds.^40^ This feedforward arm draws on the proton-circuit capacitor but tunes throughput, not the cumulative lesion that sets pace.

Each circuit obeys a distinct output law (Table 4), and in every case the aging-relevant deliverable is regenerative throughput, not the basal ATP that textbook treatments foreground. The carbon circuit runs at constant substrate and enzyme levels, so its biosynthetic flux is linear in cofactor availability (J ∝ [NAD^+^] or [CoQ_ox]) with no intrinsic buffer—the reason it fails first. The electron circuit runs at fixed V_redox, so the current that regenerates those cofactors falls as I_e ∝ 1/R_int (power likewise as P ∝ 1/R), a linear decline. The proton circuit runs at approximately constant load resistance, so its deliverable work falls as P ∝ Δ*ψ*^2^—quadratically sensitive to voltage loss. This asymmetry—linear and unbuffered for the carbon circuit, quadratic but self-relieving for the proton circuit—is why biosynthetic failure precedes energy failure.

**Table 4.**
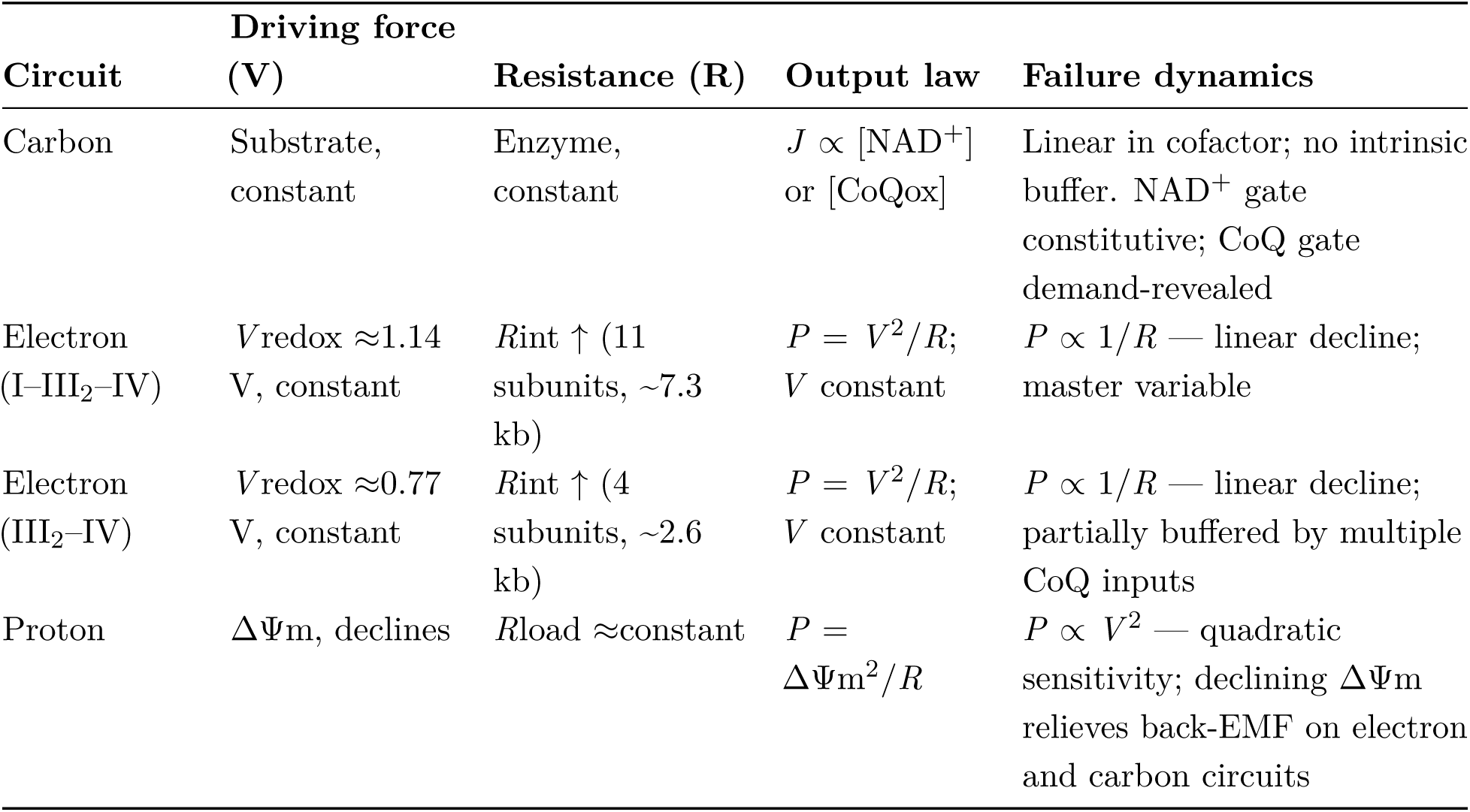
Three-circuit failure modes and dynamics of mitochondrial energy transduction. The carbon circuit declines linearly with cofactor availability, with no intrinsic buffer. The electron circuit declines linearly with impedance (P ∝ 1/R at constant redox driving force). The proton circuit declines quadratically with membrane potential (P ∝ ΔΨm^2^), amplifying sensitivity to voltage loss; however, declining ΔΨm relieves back-EMF against proton pumping, partially rescuing the electron and carbon circuits. SC, supercomplex.

| Circuit | Driving force (V) | Resistance (R) | Output law | Failure dynamics |
| --- | --- | --- | --- | --- |
| Carbon | Substrate, constant | Enzyme, constant | $J \propto [\text{NAD}^+]$ or $[\text{CoQ}_{\text{ox}}]$ | Linear in cofactor; no intrinsic buffer. $\text{NAD}^+$ gate constitutive; CoQ gate demand-revealed |
| Electron (I–III<sub>2</sub>–IV) | $V_{\text{redox}} \approx 1.14$<br>V, constant | $R_{\text{int}} \uparrow$ (11 subunits, ~7.3 kb) | $P = V^2/R$;<br>$V$ constant | $P \propto 1/R$ — linear decline; master variable |
| Electron (III<sub>2</sub>–IV) | $V_{\text{redox}} \approx 0.77$<br>V, constant | $R_{\text{int}} \uparrow$ (4 subunits, ~2.6 kb) | $P = V^2/R$;<br>$V$ constant | $P \propto 1/R$ — linear decline; partially buffered by multiple CoQ inputs |
| Proton | $\Delta\Psi_{\text{m}}$, declines | $R_{\text{load}} \approx \text{constant}$ | $P = \Delta\Psi_{\text{m}}^2/R$ | $P \propto V^2$ — quadratic sensitivity; declining $\Delta\Psi_{\text{m}}$ relieves back-EMF on electron and carbon circuits |

The two electron-entry routes carry unequal mutational targets yet erode in parallel. The respirasome (I–III_2_–IV) channels matrix NADH through eleven mtDNA-encoded subunits in series, while the III_2_–IV supercomplex accepts CoQ-pool electrons—fed by Complex II, ETFDH, DHODH, and mGPDH—through only four^39,40^. The NADH route therefore presents a larger target and a longer series chain, yet target size, series depth, and driving voltage approximately cancel, so both pathways accumulate impedance at the same fractional rate (1 + *µ*t; Supplementary Note S3.1b). The asymmetry between the NAD^+^ and CoQ gates is therefore one of *visibility, not rate*—parallel conductance erosion read out at two thresholds, not a chronological ordering of lesions. The three sections below trace each circuit—carbon, electron, proton—from this shared impedance to its distinct downstream failure.

#### 2.6.2 Carbon-circuit compression: NADH back-pressure and the NAD^+^ gate

Rising Complex I impedance progressively slows matrix NADH → NAD^+^ recycling. The steady-state oxidized fraction of the NAD pool equals *k_ox_*/(*k_red_* + *k_ox_*) and shifts continuously as Complex I conductance (*k_ox_*) declines—the most sensitive enzymes lose throughput while the bulk ratio remains heavily oxidized, so early manifestations may be subclinical. Because NADH/NAD^+^ propagates across compartments via the malate–aspartate and glycerol-3-phosphate shuttles (LaNoue, K. F. & Schoolwerth, A. C)^41^, mitochondrial back-pressure propagates systemically. The NAD^+^ pool is not only a redox couple but is also drained non-redoxly and continuously—by sirtuins, PARPs, and CD38 (Supplementary Table S8b)—a consumption that rises with age and that the purely recyclable CoQ pool never bears. This drain, layered on the redox load of maintenance catabolism, is what makes the carbon gate constitutive: it is reported at rest by every metabolically active cell, even though proliferative aspartate synthesis loads it further.

The resulting contraction is structured rather than diffuse. Because the gates are the NAD^+^- dependent steps, each Krebs intermediate gates a defined client set of downstream enzymes (Supplementary Note S3.2.1), and intermediates accumulate immediately upstream of each gated step relative to that step’s products even as total cataplerotic flux falls—a directional, redox-gated metabolite signature, not a uniform loss of TCA flux. Erosion of regenerative scope is what compresses the physiological reserve developed in §2.1.

The NNT coupling established in §2.6.1—with auxiliary routes through malic enzyme and isocitrate/IDH cycles—creates a **double-lock**: simultaneous NAD^+^ depletion and NADPH elevation from a single upstream lesion.

##### Proteome-wide consequences

Multiple domain families bind NAD(P)(H)—Rossmann-fold dehydrogenases, SDRs, AKRs, ALDHs, and P450 reductases (236 human proteins that bind NAD(P)(H) as a redox cofactor, spanning 16 structural fold families; Supplementary Table S8a)—so the cofactor shift reprograms the broader proteome, not just metabolic flux. Each NAD^+^-dependent enzyme responds according to its own *K_m_* for NAD^+^ and its sensitivity to NADH product inhibition, creating an ordered load-shedding hierarchy (the reaction and stalling consequence of each enzyme are tabulated in Table S9). The shared E3/DLD node sits near the top: PDH, *α*KGDH, BCKDH, DHTKD1, and the glycine cleavage system stall coordinately under direct NADH product inhibition (Reed, L. J.; Ambrus, A. & Adam-Vizi, V)^36,42^, producing a deterministic fingerprint—Warburg shift (PDH), BCAA accumulation with mTORC1-driven insulin-resistance signaling (BCKDH; Newgard, C. B. et al)^43^, acylcarnitine accumulation from *β*-oxidation mismatch at HADH/HADHA and ETFDH (Mihalik, S. J. et al)^44^, *α*-KG depletion with epigenetic drift via TET/JmjC starvation, and one-carbon stress (Bao, X. R. et al.)^45^ (GCS). Beyond DLD, parallel NAD^+^-dependent steps fail—serine synthesis (PHGDH), GAG precursors (UGDH), carnitine (ALDH9A1), membrane desaturation (CYB5R3)—and, through aspartate and NAD^+^-dependent purine synthesis, the carbon arm of the nucleotide checkpoint that gates cell division (§3.7). Because the shedding order is set by enzyme kinetics rather than the magnitude of Complex I loss, the earliest biosynthetic deficits are predicted decades before clinical thresholds.

#### Endocrine forks illustrate the cofactor grammar

At many branch points, NAD^+^- and NADPH-dependent enzymes catalyze antagonistic reactions. Under the double-lock the NAD^+^ arm slows down while the NADPH arm accelerates. The cortisol fork is paradigmatic: HSD11B2 (NAD^+^) deactivates cortisol to cortisone while HSD11B1 (NADPH) reactivate it, predicting tissue-level hypercortisolism in aging that coexists with declining circulating levels (Seckl, J. R. & Walker, B. R)^46^. The same grammar repeats at sex-steroid nodes and across aldehyde-detoxification, osmotic, and lipid-storage systems (Table 5); the prostaglandin and retinoid forks, with the full eight-system catalog of 72 cofactor-opposed forks, are in Supplementary Table S6, and the endocrine syndrome these forks write is developed in §3.6 (Fork inversion).

**Table 5.** Representative cofactor-opposed forks across physiological systems. Rising OXPHOS impedance generates NADH back-pressure, transduced through NNT and allied redox cycles into elevated NADPH/NADP^+^. At each fork the oxidative NAD^+^-dependent arm decelerates under impedance while the reductive NADPH-dependent arm persists; the physiological consequence is set by the product type — toxin retention, lipid storage, or a shift toward the higher-potency hormone — rather than a uniform activation, and synthesis-gated forks (e.g. progesterone) instead lose production (§3.6). The endocrine set is shown in full; one representative is drawn from each of five further systems. The complete catalog appears in Supplementary Table S6.

| System | NAD<sup>+</sup>-dependent enzyme (oxidative; decelerates) | NADPH-dependent enzyme (reductive; accelerates) | Predicted aging phenotype |
| --- | --- | --- | --- |
| Endocrine | HSD11B2 (cortisol → cortisone, inactivation) | HSD11B1 (cortisone → cortisol, reactivation) | Tissue cortisol excess despite normal or declining plasma — visceral obesity, sarcopenia, insulin resistance, hypertension |
| Endocrine | HSD17B2 (estradiol → estrone, high → low potency) | HSD17B1/3 (estrone → estradiol, low → high potency) | Local estradiol excess despite low plasma → post-menopausal hormone-sensitive breast cancer |
| Endocrine | HSD17B2 (testosterone → androstenedione, high → low potency) | AKR1C3 (androstenedione → testosterone, low → high potency) | Local androgen supply independent of gonads → castration-resistant prostate cancer |
| Endocrine | HSD3B2 (pregnenolone → progesterone, obligate synthesis) | AKR1C1 (progesterone → 20α-DHP, reductive clearance) | Double-hit — synthesis blocked while catabolism accelerates → functional progesterone deficiency, unopposed estrogen |
| Fuel storage | HADH/HADHA (β-oxidation, NAD<sup>+</sup>) | FASN/KAR (acetyl-CoA → palmitate) | Carbon forced into storage despite energy deficit → NAFLD, ectopic lipid |
| Neurotransmitter aldehyde | ALDH1A1 (DOPAL → DOPAC, detoxified) | AKR1A1 (DOPAL → DOPET, retained) | DOPAL accumulation → α-synuclein modification → Parkinson's disease |
| Neurometabolic | SSADH/ALDH5A1 (succinic semialdehyde → succinate) | AKR7A2 (semialdehyde → GHB) | Endogenous GHB accumulates → sundowning, cognitive fog, sleep-wake disruption |
| Osmotic / sugar | SORD (sorbitol → fructose, clearance) | AKR1B1 (glucose → sorbitol) | Osmotic polyol trap → cataracts, diabetic neuropathy, nephropathy |
| Genotoxic aldehyde | ADH5/GSNOR (formaldehyde → formate) | CYP demethylases (continuous formaldehyde production) | DNA-protein crosslinks and interstrand-crosslink damage → hematopoietic failure |

The NADPH the double-lock elevates is not merely pathological—it is vented through reductive sinks the cell can no longer discharge through impeded OXPHOS. Lipogenesis is the paradigmatic carbon-circuit vent:

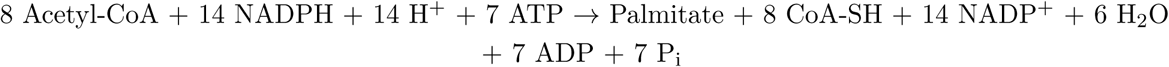

Each palmitate consumes 14 NADPH, a massive electron drain and the citrate that feeds it accumulates upstream of the stalled IDH3 gate—the one cataplerotic export that *increases* under NADH back-pressure. Additional NADPH-consuming sinks are catalogued in Supplementary Note S3.2.3; the glutathione and thioredoxin systems are developed as a reductive-stress vent in the Discussion (§3.4).

##### In vivo validation

^31^P-MRS of human brain shows intracellular NAD^+^ declining while NADH rises with age^48^, and circulating signatures of NADH reductive stress track mitochondrial-disease severity (Sharma, R. et al.)^49^. Hepatic expression of LbNOX, a bacterial NADH oxidase that recycles NAD^+^ without making ATP or pumping protons, lowers NADH/NAD^+^ and reverses insulin resistance in mice (Goodman et al., 2020)^50^—extending to whole-organism physiology the foundational finding that the essential biosynthetic function of the respiratory chain is NAD^+^ regeneration, not oxidative phosphorylation per se (Titov et al., 2016)^51^.

#### 2.6.3 Electron-circuit saturation: CoQ over-reduction and the CoQ gate

As Complexes III and IV accumulate lesions, CoQH_2_ clearance slows and the quinone pool shifts toward a more reduced steady state; CoQH_2_ generation need not chronically exceed clearance for this to occur, because the pool redistributes continuously as the balance between reduction and oxidation changes. The CoQ pool obeys the same steady-state logic as the NAD pool: with *k_red_* the rate constant for CoQ reduction and *k_ox_* that for CoQH_2_ oxidation through Complexes III–IV, the oxidized fraction is *k_ox_*/(*k_ox_* + *k_red_*)—∼91% at a tenfold ratio, ∼67% at twofold, 50% when the two equalize. Progressive over-reduction also creates the thermodynamic conditions for reverse electron transport (RET) through Complex I, a potent superoxide source that links the CoQ gate to accelerated oxidative damage (Murphy, M. P.; Chouchani, E. T. et al)^52,53^.

Each CoQ-dependent enzyme has its own affinity for oxidized CoQ, so activities fall in order of substrate sensitivity as the oxidized fraction contracts. DHODH—which supplies the pyrimidine ring and has the lowest maximal flux among the major CoQ-dependent dehydrogenases—is predicted to be among the earliest casualties, the pyrimidine arm of the dual checkpoint (§3.7), consistent with the proliferative failures (immune senescence, impaired wound healing) that characterize demand-dependent aging.

The over-reduced pool also drives the electron circuit’s own relief valve. RET-generated superoxide is vented through the glutathione cycle, which consumes one NADPH (two electrons) per pair of leaked electrons—coupling this vent back to the NADPH pool elevated by the carbon-circuit double-lock and draining OXPHOS and NADPH simultaneously, which is why concurrent oxidative and reductive stress in aged tissue are causally linked rather than coincidental. CRISPR synthetic-lethality data and natural models of complete Complex I loss (renal oncocytoma, Hürthle-cell carcinoma) independently corroborate these sinks; the carrier-linked dependencies underlying this vulnerability are detailed in Supplementary Note S3.2.2 and Table S5. The CoQ gate thus broadens the lesion from the selective NADH-clearance defect of the NAD^+^ gate into a second front of biosynthetic failure at the shared CoQ node.

#### 2.6.4 Proton-circuit brownout: Δ*ψ* decline and selective load-shedding

The proton circuit fails on a third logic. Δ*ψ* is a capacitor voltage that gates work, not a consumable cofactor pool, so as it drifts downward the failure appears as a brownout: by the threshold logic established in §2.6.1, voltage-dependent work is selectively shed before basal ATP content falls.

Mitochondrial protein import is the leading casualty. Nuclear-encoded precursors cross the outer membrane through TOM independently of voltage, but presequence-driven import across the inner membrane requires Δ*ψ*-dependent translocation through TIM23 and the PAM motor. Declining Δ*ψ* therefore slows import of the very nuclear-encoded respiratory-chain subunits, assembly factors, and carriers needed to renew the chain—a feedback loop in which rising impedance lowers Δ*ψ*, lower Δ*ψ* impairs renewal, and impaired renewal further erodes conductance reserve. Fe–S cluster export and the Δ*ψ*-dependent mitochondrial steps of steroidogenesis (StAR-mediated cholesterol import, CYP11A1) are early casualties on the same threshold logic: their reserve falls before resting ATP, so the deficit surfaces first as blunted stimulated output rather than resting deficiency, with the age-related decline of DHEA-S—the most age-sensitive steroidogenic readout—as its recognizable signature, even while resting ATP appears normal^72^.

Basal ATP synthesis occupies the opposite, low-Δ*ψ*-threshold end of the same spectrum and is therefore defended longest—the more so because cells can suppress demand and reroute substrate long before energy charge falls. The resulting prediction is preserved resting ATP with impaired ATP *kinetics*: slowed recovery and a reduced capacity to fund repair, contraction, immune activation, and biosynthesis at once, rather than acute energetic failure. Basal ATP is thus the defended floor across all three circuits; what erodes first in each is regenerative, not energetic, capacity.

The proton circuit is coupled to the other two through its voltage, and the coupling cuts both ways. Declining Δ*ψ* relieves the back-EMF opposing proton pumping, partially rescuing electron and carbon flux; the same decline deprives NNT of the driving force for its NADH → NAD^+^ reaction, narrowing the one route that links proton-circuit capacity to carbon-circuit recovery. Regulated uncoupling (UCP1–5) and passive leak serve as the proton circuit’s own vent, dissipating Δ*ψ* to relieve back-EMF at the cost of voltage-dependent work. No internal redistribution satisfies all three circuits at once—the trade-off the interventional discriminator in §3.9 exploits. Beyond these three regressors, the framework implies further, largely nuclear-encoded determinants of encoded chain quality and chain maintenance; their relation to the residual lifespan variance is taken up in §3.10.

##### Carbon circuit (left; constitutive, NAD^+^ gate) — compression

Rising NADH back-pressure throttles the NAD^+^-gated dehydrogenases of the Krebs rheostat in every resting cell, contracting biosynthetic carbon flux: *α*-ketoglutarate depletion starves 2-oxoglutarate oxygenases (epigenetic maintenance via TET/JmjC; collagen/ECM hydroxylation), while one-carbon, anaplerotic, and aspartate-derived nucleotide outputs decline.

##### Electron circuit (centre; demand-revealed, CoQ gate) — saturation

Chain impedance over-reduces the CoQ pool; CoQ-dependent enzymes fail in order of acceptor affinity—DHODH-gated pyrimidine synthesis earliest, then ETF/ETFDH-dependent fatty-acid and amino-acid oxidation—so the constraint is revealed only under proliferative or catabolic demand.

##### Proton circuit (right; threshold-gated, Δ*ψ*) — brownout

Falling electron flux lowers the proton-motive force (Δ*ψ*_m_, ΔpH); because deliverable power scales as Δ*ψ*^2^, high-threshold loads are shed first—TIM23/PAM protein import, StAR/CYP11A1 steroidogenesis, Fe–S cluster export, metabolite/ion transport—while basal ATP synthesis, the lowest-threshold load, is the defended floor across all three circuits.

##### Dual checkpoint

The carbon (aspartate/purine) and electron (DHODH/pyrimidine) arms converge on a single shared node, de novo nucleotide synthesis; its failure imposes replicative arrest, the reason dividing compartments (immune expansion, wound healing, stem-cell mobilization) are doubly hit.

The circuits run in series—the carbon canopy’s reducing equivalents feed the electron roots, the electron canopy’s proton pumping feeds the proton roots—over a shared foundation of metabolic scope / bioenergetic reserve. A dashed feedback marks declining Δ*ψ* relieving back-EMF, partially rescuing upstream flux. Across the figure the biosynthetic canopy thins while basal ATP is preserved: biosynthetic failure precedes energy failure. Solid arrows, direct consequences and series flow; dashed arrow, regulatory feedback. Leaves denote functional classes; representative enzymes and phenotypes are shown.

## Discussion

Aging is progressive biosynthetic failure. The three-circuit analysis of §2.6 traces this to rising mito-chondrial impedance, read out through three asymmetric gate modes: constitutive **carbon-circuit compression**, as NADH back-pressure restricts NAD^+^-gated dehydrogenases and biosynthetic carbon flux; demand-revealed **electron-circuit saturation**, as an over-reduced CoQ pool limits oxidized acceptor capacity for DHODH, ETFDH, SDH, and mGPDH during proliferative, catabolic, or stress load; and threshold-gated **proton-circuit brownout**, as declining Δ*ψ* selectively sheds voltage-dependent mitochondrial work. Resting ATP is defended longest; what erodes first is not energy content but exergy — the usable free-energy headroom that drives anabolic, repair, immune, endocrine, and regenerative work.

MSTA therefore makes four linked claims. First, aging is driven by progressive biosynthetic contraction rather than resting ATP deficiency. Second, the sequence of age-related phenotypes is ordered by the engagement modes of three mitochondrial gates — constitutive NAD^+^-gate compression, demand-revealed CoQ-gate saturation, and threshold-gated Δ*ψ* brownout. Third, the Scope–Stability–Pace (SSP) law specifies the timescale over which this deterioration unfolds across species. Fourth, the SSP axes constitute the physical substrate of disposable soma — the mitochondrial conductance reserve that classical evolutionary theories posited as “maintenance” without specifying its molecular implementation. Species differ in longevity because they occupy different viable solutions on a reserve–erosion manifold; individuals age because their inherited solution is progressively consumed. Selection does not optimize lifespan but fitness, and high-SSP coordinates are favored only where reproductive and ecological context makes the cost worth paying. These claims place MSTA upstream of, rather than in competition with, the descriptive frameworks of aging biology. Hallmark taxonomies catalogue recurrent features of aged tissue without specifying the constraint that orders them^54,55^; MSTA proposes a circuit-level ordering. Carbon-circuit compression links NAD^+^ restriction, *α*-KG depletion, aspartate limitation, and one-carbon stress to epigenetic drift, impaired DNA repair, ECM decline, altered oxygen sensing, and stem-cell exhaustion. Electron-circuit saturation links CoQ acceptor limitation to DHODH-dependent pyrimidine restriction, impaired proliferative immunity, delayed wound healing, *β*-oxidation mismatch, and RET-linked ROS. Proton-circuit brownout links Δ*ψ* loss to impaired TIM23/PAM-dependent import, OXPHOS-renewal failure, steroidogenic decline, proteostatic stress, and slowed recovery kinetics. In this framing, hallmarks are not independent causes arranged in parallel but partially overlapping tissue projections of constrained mitochondrial conductance. Information-theoretic accounts, which attribute progressive dysfunction to loss of epigenetic information^56^, become one major readout of the same thermodynamic lesion: the *α*-KG and NAD^+^ contraction above is itself the upstream constraint that starves the enzymatic machinery preserving chromatin state.

The neglect of Scope as a longevity axis has deep historical roots. Longevity biology inherited a rate-centric lens from Rubner (1908) and Pearl (1928), mechanized through Harman’s free-radical theory (1956), which made resting metabolic rate the default allometric variable for cross-species lifespan analysis. The allometric signal of Scope is genuinely shallow: factorial scope scales as approximately *BM* ^0^.^148^, so a doubling of body mass raises scope by only ∼10%, and the full mammalian range spans only ∼4-fold — readily obscured by within-species measurement variability in small, taxonomically heterogeneous compilations. In that context, Schmidt-Nielsen’s influential synthesis — built on ∼22 mostly domesticated species — concluded that factorial aerobic scope (VO_2_max/BMR) is ∼10 and approximately mass-invariant, reframing structured species-level variation as “athletic” idiosyncrasy rather than a systematic axis of design^57^. Because VO_2_max is also far harder to standardize across taxa than BMR, datasets stayed sparse for decades and reserve capacity remained underweighted — precisely the gap MSTA closes.

What that axis measures deserves to be stated precisely, because the empirical proxy and the underlying construct are not the same quantity. The life-limiting variable is *conductance reserve*: the dimensionless headroom between young-adult OXPHOS conductance and the minimum conductance required to sustain regenerative biosynthesis above basal maintenance—the cellular *regenerative scope* of §2.6.2 and Fig. 8. Aging consumes this reserve as rising impedance lowers conductance toward the maintenance floor, while Stability and Pace set the rate of erosion. Body mass is its allometric proxy in the SSP law, and whole-organism factorial scope (VO_2_max/BMR; Fig. 1B) is its organismal shadow; the two share a body-mass exponent (∼0.15), which is the parallel Fig. 1B documents. But factorial scope is dominated by locomotor muscle, and the structure–function series of Weibel et al.^15^ shows that the oxidative rate per unit mitochondrial volume is invariant across mammals while VO_2_max tracks mitochondrial quantity—so aerobic-scope differences between athletic and sedentary species at matched mass are differences of muscle quantity, not of the cellular conductance reserve that limits lifespan. This is consistent with the observation that elite aerobic performers do not dominate extreme-longevity demographics (§3.10): adding locomotor mitochondrial quantity raises VO_2_max without enlarging the cellular reserve, so locomotor scope and cellular conductance reserve are expected to diverge along the athletic axis. The construct is therefore best tested not by VO_2_max but by its direct cellular correlates—tissue-level spare respiratory capacity, the maximal-to-basal conductance ratio, and cristae/respirasome architecture—whose decline with age is well documented and whose cross-species ceiling the SSP body-mass term is taken to index.

### 3.1 MSTA operates across scales

Across species, SSP coordinates set the inherited trajectory: Scope determines how much impedance can be absorbed before regenerative throughput falls below maintenance demand, while Stability and Pace determine how quickly that impedance accumulates. Within species, individuals vary around this trajectory through stochastic mtDNA damage, mitochondrial quality control, endocrine operating point, and tissue loading — thyroid tuning Pace, GH/IGF-1 tuning growth-versus-maintenance allocation, insulin tuning substrate pressure, sex steroids tuning regenerative demand.

Exceptional longevity is therefore predicted to combine favorable inherited SSP coordinates with low-impedance endocrine set-points, consistent with lifespan extension in Laron syndrome, Ames dwarf mice, and FOXO3-variant centenarians^58–60^. At the cellular level, the same impedance state is read out through three gates: constitutive NAD^+^-gate compression in all metabolically active cells, demand-revealed CoQ-gate saturation in proliferating or catabolically stressed cells, and threshold-gated Δ*ψ* brownout in voltage-dependent mitochondrial functions. Tissue specificity therefore does not require a separate aging mechanism for each organ; it reflects different local loading of one three-gate architecture, even as decline remains synchronized across tissues at the decadal scale of the clinical syndrome.

Reversibility is bounded by the impedance layer targeted. Operational impedance — NADH/NAD^+^ ratios, CoQ redox state, Δ*ψ* set-point, supercomplex distribution, substrate pressure, and endocrine tone — is rapidly reversible; caloric restriction, exercise, NAD^+^ precursors, and hormetic stressors act at this layer and produce the measurable but capped lifespan extensions observed across model organisms^61^. Architectural impedance — cristae geometry, cardiolipin composition, respirasome as-sembly — is partially reversible on longer timescales through biogenesis and turnover. Informational impedance — accumulated mtDNA point mutations, clonal heteroplasmy expansion, deletion load, and the somatic cell and fiber losses they drive — defines the hard ceiling: what has crossed from operational dysregulation into informational loss cannot be restored by regulatory intervention alone. Somatic-cell nuclear transfer illustrates the partition: Dolly the sheep developed normally from a six-year-old donor nucleus^62^ because the oocyte cytoplasm reset the regulatory and architectural layers while supplying a fresh mitochondrial environment — nuclear transfer succeeds precisely because it bypasses the informational layer rather than repairing it. MSTA thus predicts a ceiling to rejuvenation set not by thermodynamics in general but by the fraction of accumulated impedance that has become informationally fixed.

This framework supplies a theoretical definition of biological age: cumulative exposure to rising OXPHOS impedance — the integrated history of conductance loss, cofactor contraction, CoQ saturation, and Δ*ψ* erosion that chronological time only approximates. Methylation clocks, frailty indices, and reserve-based measures such as VO_2_max decline, handgrip, and gait speed can be read as different empirical projections of this state variable, which is why they correlate with one another and with mortality without any of them fully capturing the others.

A first-principles clinical signature follows: aging tissues should exhibit critical slowing down — slower return to baseline after perturbation — even when resting state variables appear preserved^63^. Heart-rate recovery after exercise, post-prandial glucose clearance, lactate disposal, and post-illness immune and cognitive resolution should therefore outperform resting biomarkers as indices of cumulative impedance, and durable interventions should improve recovery kinetics, not merely resting levels.

### 3.2 The Krebs Rheostat

The Krebs Rheostat is the tissue-level expression of carbon-circuit compression (Fig. 8). The coordinated dimming of the near-equilibrium and E3/DLD-gated dehydrogenases derived in §2.6.2 converts a continuous redox variable — NADH/NAD^+^ — into graded loss of biosynthetic throughput,

producing selective contraction at NAD^+^-gated steps rather than uniform pathway decay. Beyond the enzyme-level gating, extrinsic NAD^+^ consumption by CD38 and PARPs further contracts the oxidized cofactor pool^64,65^. In acute stress this is adaptive; in aging it becomes chronic restriction. This rheostat produces a two-sided squeeze on the carbon circuit. On the entry side, NADH product inhibition of PDH and the oxidative dehydrogenases means the cycle cannot accept new carbon skeletons even amid nutrient abundance—an *anaplerotic bottleneck* in which the cycle is saturated with electrons, not carbon. On the exit side, intermediates accumulating upstream of the stalled NAD^+^ steps are forced out as *cataplerotic escape*—citrate export to ATP-citrate lyase driving de novo lipogenesis, which recasts hepatic steatosis, intramyocellular lipid, and insulin resistance as overflow valves under a redox ceiling rather than primary defects (§3.4; Supplementary Note S5.1). Between a blocked entry and a leaking exit, the intermediate pool shrinks, imposing a passive kinetic brake.

Consistent with this prediction, untargeted metabolomics in aging mouse hippocampus shows TCA intermediates (cis-aconitate, succinate) declining 25–47% despite abundant lipid and glutamine inputs, with a 2.3–2.5-fold shift specifically at NAD^+^/NADH-dependent dehydrogenase sites^66^ — the pattern expected from redox gating, not random TCA collapse. Because *α*-ketoglutarate is the obligate co-substrate for over 60 Fe(II)/2-oxoglutarate-dependent oxygenases^67,68^ — JmjC/TET demethylases, HIF-prolyl hydroxylases, collagen hydroxylases — its contraction links the same upstream throttle to epigenetic drift, altered oxygen sensing, and impaired collagen maturation in parallel^69^. The familiar metabolic fingerprint of aging — Warburg shift, BCAA accumulation, *α*-KG depletion, disrupted glycine metabolism — is therefore interpretable as one rheostatic program rather than multiple independent failures.

### 3.3 Gate engagement, not lesion order, determines phenotype order

Aging phenotypes appear in a recognizable order because downstream functions engage the three mitochondrial gates differently. Phenotype timing does not report which OXPHOS component was damaged first; it reports which downstream gate has the lowest reserve margin in a given tissue under a given load.

The carbon/NAD^+^ gate is constitutive: every resting cell reports NADH back-pressure through NAD^+^-dependent dehydrogenases and biosynthetic nodes, so insulin resistance, BCAA accumulation, lactate pressure, *α*-KG depletion, impaired collagen maturation, and epigenetic drift emerge gradually and systemically, requiring no overt respiratory failure. The CoQ gate is demand-revealed: electron-circuit saturation becomes visible only when tissues attempt high-throughput proliferation, *β*-oxidation, immune activation, wound healing, hematopoiesis, or stress recovery — a resting tissue may appear adequate while an activated T cell, healing epithelium, fasting liver, or regenerating stem-cell compartment fails because DHODH, ETFDH, SDH, or mGPDH cannot access sufficient oxidized CoQ. The proton gate is threshold-gated: TIM23/PAM import, Fe–S export, steroidogenic mitochondrial steps, carrier exchange, and stress recovery have different voltage thresholds and become unreliable while resting ATP remains defended.

This gate logic resolves an apparent paradox: the same proton circuit supports both ATP synthesis and non-ATP work, yet steroidogenesis, import, and recovery kinetics can decline long before basal energy charge falls, because each is a higher-threshold load than basal ATP synthesis (§2.6.4). Phenotype timing therefore reports the vulnerability of the local gate, not the rank of the upstream lesion: a steroidogenic cell reveals proton brownout as falling pregnenolone and DHEA-S; skeletal muscle reveals it as delayed phosphocreatine recovery; an activated T cell reveals the dual NAD^+^/CoQ nucleotide checkpoint; liver reveals carbon compression as lipogenesis and insulin resistance. The shared lesion is rising impedance; the visible phenotype is set by local gate threshold. Independent support for this ordered cascade comes from temperature manipulation in a non-mammalian vertebrate. Rearing *Nothobranchius rachovii* across 20–30 °C accelerates aging in proportion to temperature, and the accompanying molecular changes track the circuit architecture developed here: Δ*ψ*m falls as the proton circuit loses voltage; mitochondrial density rises in compensation; and the NAD^+^-dependent SirT1–FoxO axis and its downstream antioxidant targets attenuate in step with mitochondrial decline^30^. The same ordered pattern inferred from mammalian aging is recovered, under controlled thermal manipulation, in a teleost fish.

### 3.4 Compensations as metabolic debt

The containment responses derived in §2.6 share a common character: they preserve short-term viability by creating downstream liability. They are not independent pathologies layered onto aging; they are the costs of keeping basal ATP defended while regenerative scope contracts.

Carbon-circuit compression vents through lipogenesis. Citrate accumulating upstream of the IDH3 bottleneck exits to support acetyl-CoA and fatty-acid synthesis, consuming NADPH and storing reductant as lipid — a biochemical electron sink when OXPHOS cannot discharge reducing equivalents. Short-term this vents redox pressure; chronically it produces ectopic lipid, hepatic steatosis, and substrate inflexibility.

The electron valve is ROS, and it draws on two distinct redox-pressure sites that should not be conflated. Carbon-circuit compression promotes ROS at stalled E3/DLD flavin centers, where NADH product pressure accumulates at the 2-oxoacid dehydrogenases^70^; electron-circuit saturation promotes reverse electron transport through Complex I when an over-reduced CoQ pool coexists with maintained Δ*ψ*^52,53^. Detoxification of both through the glutathione and thioredoxin systems consumes NADPH, which is why oxidative and reductive stress in aged tissue are causally coupled rather than coincidental: the same congestion that raises NADH and CoQH_2_ drives the leak, while clearing it drains the NADPH pool the carbon double-lock has already elevated. This also explains why antioxidant supplementation has shown limited lifespan benefit^71^ — it blocks the vent without relieving the impedance that necessitated it. Quantitatively the vent is larger than its reputation. Clearing one superoxide costs ∼0.5 NADPH — two radicals dismutate to one H_2_O_2_, reduced by two glutathiones and regenerated by one NADPH — so leak and clearance together reduce O_2_ to water by four electrons, two bled from the over-reduced chain and two from NADPH: an improvised terminal oxidation that discharges reducing equivalents around an impeded Complex IV. The NADPH it spends is the matrix pool that NNT regenerates from the very NADH back-pressure driving the leak, making the antioxidant network a principal fate of the double-lock’s matrix surplus — an in-compartment loop distinct from cytosolic lipogenesis, and one whose flux can match the lipogenic vent because it rides on the entire respiratory electron flux. It also scales with the lesion: leak fraction rises with Δ*ψ* and CoQ over-reduction, so the vent widens as impedance worsens. Oxidative damage is thus buffered while the surplus lasts and surfaces only when reductive pressure outruns NADPH-limited containment, placing oxidative stress downstream and late in life.

This temporal logic is sharply illustrated by mitochondria-targeted catalase. Catalase clears H_2_O_2_ to water at no NADPH cost, but it is peroxisomal and cytosolic and nearly absent from the matrix, where the low-Km glutathione and thioredoxin peroxidases—not the high-Km catalase—are kinetically matched to physiological H_2_O_2_. Forcing catalase into the matrix (MCAT mice) raises median lifespan by ∼20% (∼5 months) and maximum by ∼5.5 months, reduces mitochondrial deletions, and delays cardiac and oxidative pathology, whereas peroxisomal or nuclear targeting does little—and the benefit accrues in old animals, not young.^84^ MSTA reads this not as antioxidant rescue but as cleaner disposal of a product the saturated glutathione vent can no longer contain: an NADPH-independent sink at the source, engaged only once impedance has driven H_2_O_2_ past containment, and idle earlier, given catalase’s high Km, so the native vent and low-level redox signaling are spared. Its limit is equally diagnostic—the mortality curve shifts later without a change in slope, delaying the threshold crossing rather than slowing impedance accumulation: the bounded benefit expected of a compensation that contains the product without restoring conductance.

Proton-circuit brownout recruits uncoupling and leak: dissipating Δ*ψ* relieves back-EMF and reduces electron pressure, but spends the same voltage required for import, carrier exchange, steroidogenesis, and ATP synthesis under demand. Glycolytic compensation likewise defends ATP while raising lactate burden and substrate pressure. Each valve buys time by shifting cost elsewhere, so aging progresses not only through primary lesions but through accumulated compensation costs — which is why blocking any single downstream phenotype, without restoring conductance, routes pressure through another valve and fails to extend lifespan.

### 3.5 Symptoms trace thresholds, not lesions

The sequence of aging phenotypes is governed by two orderings that are often conflated. The first is backbone-bottleneck order — the sequence in which the three circuits lose conductance (§2.6). The second is phenotype order — the sequence in which dysfunction becomes clinically visible — governed not by which backbone is damaged first but by the currency threshold at each downstream gate. The two can diverge sharply: high-threshold, Δ*ψ*-dependent work fails clinically decades before resting ATP. The cleanest signature is kinetic—slowed restoration of Δ*ψ* and ATP after a load, seen as delayed phosphocreatine and heart-rate recovery while basal energy charge appears normal. Voltage-dependent biosynthetic loads follow the same threshold logic: the reserve of mitochondrial steroidogenesis falls early too, surfacing as blunted stimulated output with declining DHEA-S as its recognizable readout.

A direct diagnostic implication follows: because basal ATP is defended and demand is suppressible, resting measurements often appear normal while reserve is already eroded, so challenge responses should outperform them. Carbon-circuit compression should be most visible in NADH/NAD^+^-sensitive ratios and post-load recovery — lactate/pyruvate, BCAA/ketoacid balance, *α*-KG-linked metabolites, aspartate availability, one-carbon flux, ^31^P-MRS NAD redox, and post-prandial clearance. Electron-circuit saturation should surface under proliferative or catabolic challenge — uridine-rescuable proliferation, acylcarnitine accumulation on fasting or exercise, and immune-expansion and wound-healing kinetics. Proton-circuit brownout should surface in voltage-dependent recovery — phosphocreatine and heart-rate recovery, Δ*ψ*-sensitive import assays, steroidogenic stimulation tests, and delayed recovery after illness or exertion. MSTA therefore predicts that the most informative biomarkers of biological age will be dynamic, gate-specific, and challenge-based; Tables S2 and S3 map each predicted phenotype to its parent gate, required currency, expected engagement mode, and informative ratio biomarker. These challenge tests are gate-resolved; a complementary systemic integrator is needed to report cumulative impedance across all three gates from a single sample. Circulating GDF15 — induced by the integrated stress response that reductive NADH back-pressure is sufficient to trigger, and rising with FGF21 under the same constraint — is that integrator: it is the leading circulating marker of mitochondrial energy resistance, scales with the severity of mtDNA defects, and tracks human longevity, falling in the offspring of long-lived parents. MSTA therefore adopts GDF15, with FGF21, as the resting-state readout of accumulated OXPHOS impedance, complementary to the dynamic, gate-specific markers above^83^.

### 3.6 Fork inversion

Fork inversion is not a fourth circuit; it is the endocrine, inflammatory, and signaling expression of the carbon-circuit double-lock established in §2.6.2. As rising impedance raises NADH and restricts NAD^+^ regeneration, reductive pressure is transmitted into the NADPH pool through NNT, isocitrate/IDH cycling, and malic-enzyme exchange. Because NNT runs forward only while Δ*ψ* powers it, the NADPH arm of the double-lock is Δ*ψ*-contingent: elevation while the proton gate holds, starvation as it fails. While Δ*ψ* is intact the coupling is initially protective, chronically leaving NADPH-dependent reductive arms favored relative to their stalled NAD^+^-dependent partners, imposing directional bias on cofactor-opposed enzyme pairs. As Δ*ψ* declines, NNT forward transhydrogenation slows while NADH/NAD^+^ remains high, and the NADPH arm inverts from surplus to deficit — so the late lock is the loss of NADPH-dependent reductive and antioxidant capacity rather than its excess, the crossover at which the oxidative stress of aged tissue surfaces. Read across the endocrine forks of Table 5 (§2.6.2), the consequence is not simply more or less hormone but tissue-level signal distortion: clearance arms fail while activation arms persist, so local hormone action can rise even as circulating production declines. This is the mechanistic bridge from mitochondrial impedance to the signal–output decoupling characteristic of aging physiology — insulin without glucose disposal, gonadotropins without steroidogenesis, anabolic drive without protein synthesis, inflammatory activation without resolution.

The progesterone row illustrates why fork inversion need not always raise active hormone: some branches are double-hit, where NAD^+^ is required for synthesis while NADPH-supported clearance persists, so the prediction is collapse of production with altered shunting rather than local amplification. The 11-oxygenated androgen pathway is similarly layered (Table S6), with sex-specific consequences^47^. Fork inversion is therefore a grammar, not a single endocrine direction: cofactor architecture determines which arm prevails when the NAD^+^/NADPH balance shifts.

### 3.7 OXPHOS integrity constitutes a dual checkpoint for cell division

De novo nucleotide synthesis lies at the intersection of constitutive NAD^+^-gate compression and demand-revealed CoQ-gate saturation; every dividing cell must clear both gates. The carbon/NAD^+^ arm supplies aspartate, mitochondrial one-carbon units, and purine support: purine synthesis requires NAD^+^ at IMPDH and at MTHFD2-dependent formate supply, and both purines and pyrimidines require aspartate, whose production is respiration- and NAD^+^-gated^74,75^. The CoQ/saturation arm supplies DHODH-dependent pyrimidine capacity: DHODH oxidizes dihydroorotate by passing electrons to oxidized CoQ^73^, so an over-reduced pool imposes acceptor limitation precisely in dividing cells. That this arm is gated by oxidized-CoQ availability rather than NAD^+^ status is shown directly by alternative-oxidase rescue of pyrimidine synthesis (§3.9).

Progressive impedance therefore imposes an increasingly stringent dual checkpoint on every dividing cell, predicting the convergent decline of stem-cell mobilization, immune expansion, and epithelial renewal. Aged naïve T cells, which show simultaneous defects in mitochondrial respiration and one-carbon metabolism that impair activation^76^, provide in vivo support for both arms. The model predicts that combined uridine (bypassing DHODH) and aspartate or formate supplementation should rescue proliferative capacity in aged progenitors ex vivo without restoring OXPHOS conductance — a result that would confirm, rather than refute, that the defect lies downstream of the two gates; failure of such bypasses would instead implicate proton brownout or irreversible damage.

### 3.8 Timescale across species

The SSP law specifies the timescale; the three-gate circuit map specifies the phenotype sequence. Scope sets how much impedance can be absorbed, while Stability and Pace set how fast it accumulates; Figure 7 describes what happens locally as that interval is traversed, as carbon-circuit compression, electron-circuit saturation, and proton-circuit brownout progressively erode regenerative work while basal ATP remains defended. The fitted coefficients translate into near-equivalent longevity weights:

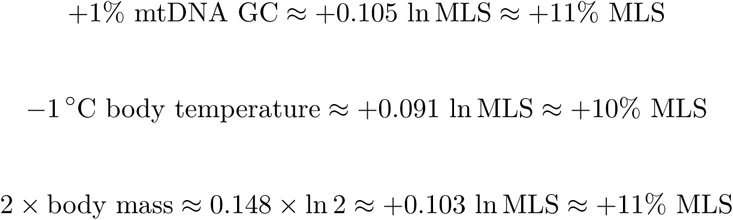

**Figure 7.**
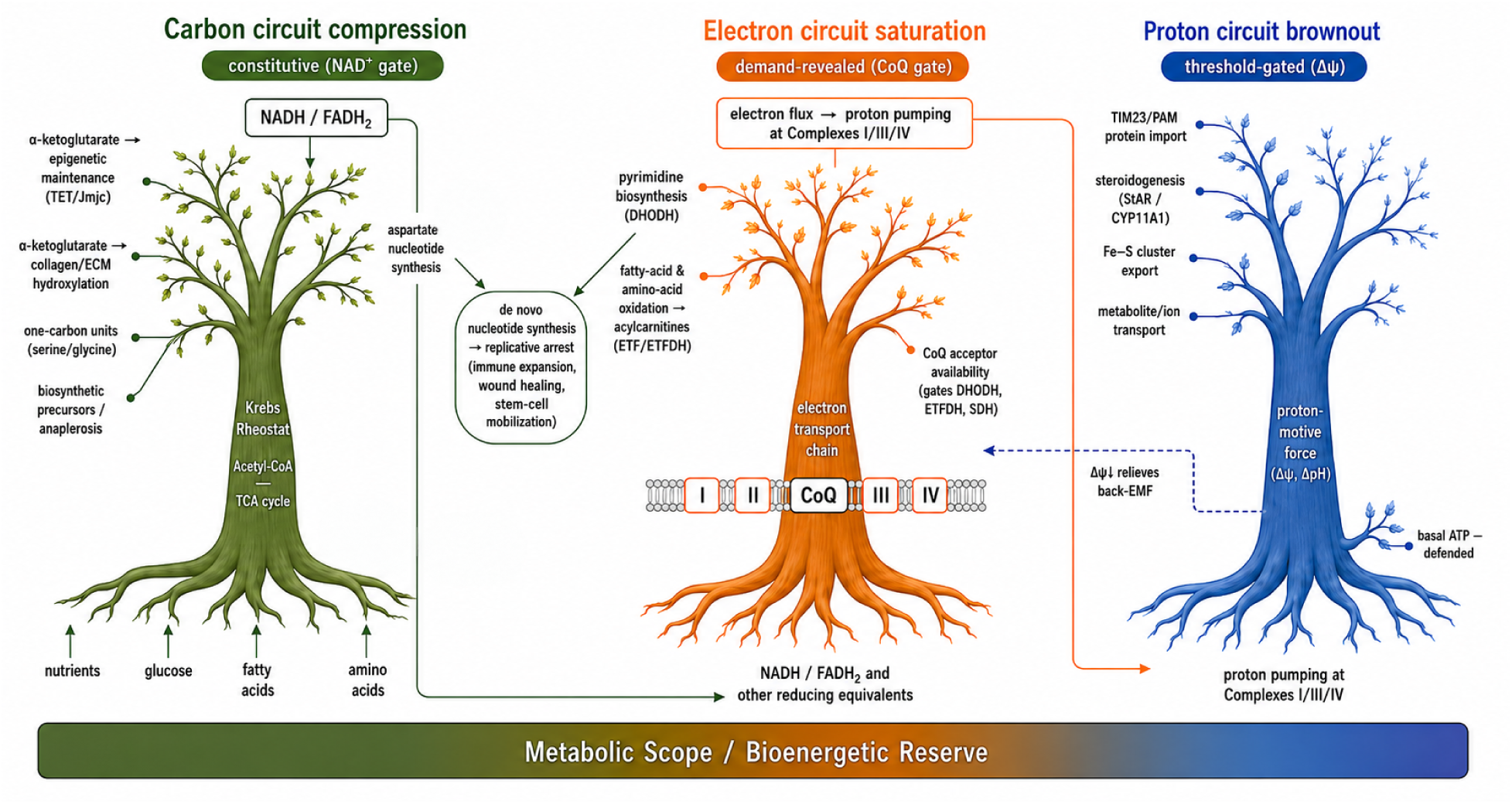
Rising OXPHOS impedance is read out through three coupled mitochon-drial circuits, each with a distinct failure mode. A single upstream lesion—the progressive, parallel rise in internal resistance (*R_int_*) across the respiratory chain—manifests as three concurrent constraints, not a temporal sequence of failures. Each circuit is drawn as a tree: substrates and upstream currency enter at the roots, the circuit core forms the trunk, and the dependent downstream processes form the canopy, which thins as the circuit is compressed.

**Figure 8.**
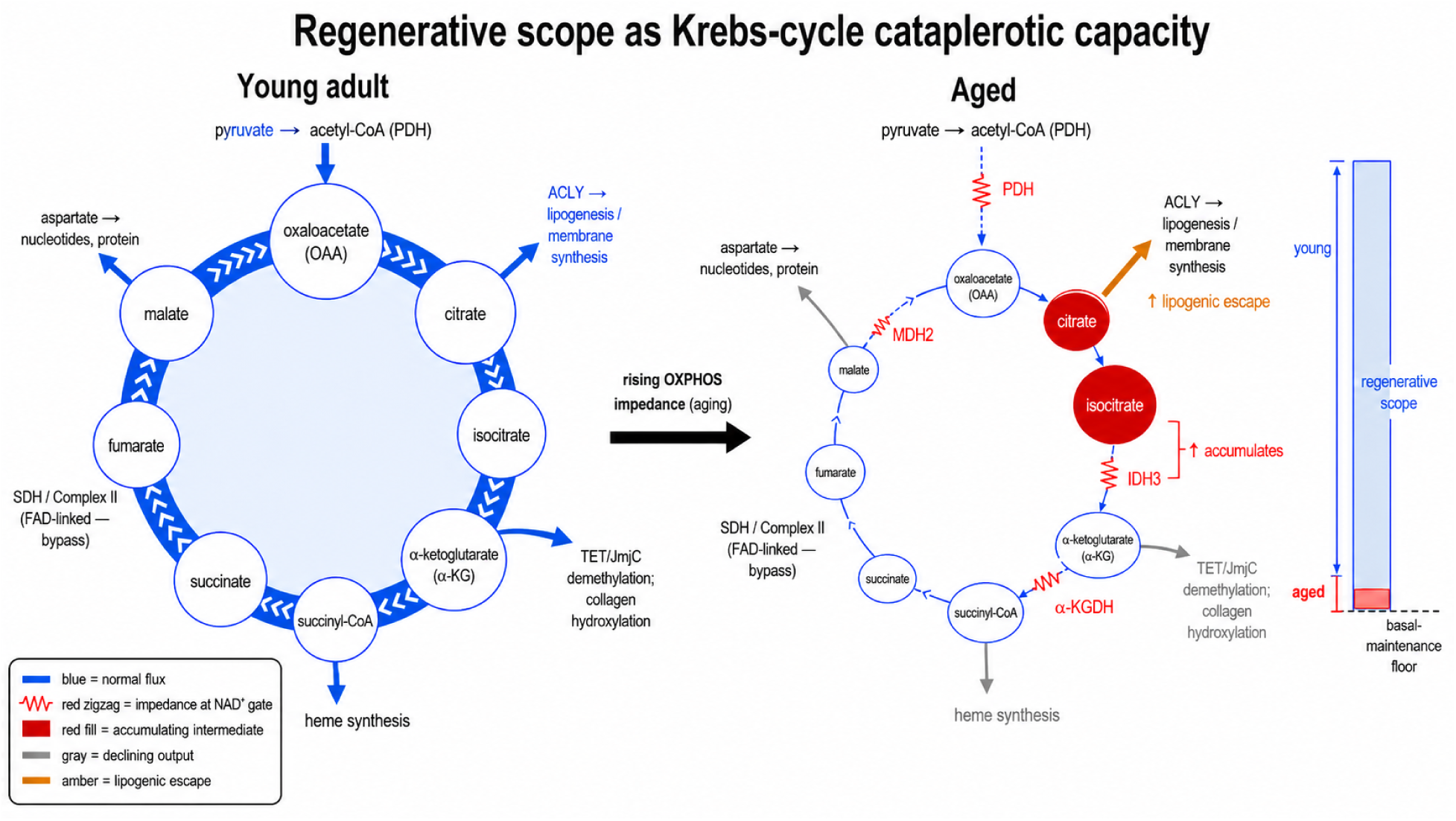
Regenerative scope as Krebs-cycle cataplerotic capacity. The Krebs/TCA cycle drawn as a redox-gated distribution network whose spare cataplerotic throughput — its regenerative scope — supports biosynthesis above the basal energetic requirement. Forward carbon flow is gated by the NAD^+^-dependent dehydrogenases (PDH, IDH3, *α*-ketoglutarate dehydrogenase [*α*-KGDH], and MDH2), collectively the Krebs rheostat; the FAD-linked succinate dehydrogenase step (SDH/Complex II) is not NAD^+^-gated and is drawn as a bypass. Left (young adult): a large, well-stocked intermediate pool (filled ring) turning over rapidly (dense flux arrowheads) sustains every cataplerotic output. Right (aged): rising Complex I impedance raises the matrix NADH/NAD^+^ ratio, and the resulting back-pressure raises resistance at each NAD^+^-gated step (red impedance symbols; dashed, thinned arrows denote reduced flux), slowing turnover and shrinking the intermediate pool (thin, depleted ring). The result is a directional metabolite signature rather than uniform pathway depletion: because IDH3 is the dominant bottleneck, citrate and isocitrate accumulate upstream (red) even as total cataplerotic flux falls, and the surplus citrate is exported — enlarging lipogenesis and membrane synthesis (amber), the one cataplerotic branch that increases, a lipogenic escape route. The remaining regenerative outputs decline (gray): oxaloacetate → aspartate and nucleotide synthesis, *α*-ketoglutarate → TET/JmjC-dependent demethylation and collagen/ECM hydroxylation, and succinyl-CoA → heme synthesis. The gauge (right) defines regenerative scope as the spare cataplerotic capacity above the basal-maintenance floor — ample in youth and compressed toward the floor with age, as rising impedance consumes the reserve.

The three axes therefore contribute almost equally per natural unit of variation, so the viable mammalian solution space is balanced rather than dominated by any single term — Scope, Stability, and Pace compensate for one another rather than acting independently. Birds remain long-lived despite high Pace because elevated Stability and expanded aerobic Scope offset the thermal penalty. Fish reach mammalian-grade longevity despite low absolute metabolic power because low Pace slows damage accumulation while factorial scope remains sufficient; and because thermal deamination pressure is itself low, GC% enrichment confers little additional benefit, so the Stability premium is conditional on Pace (§2.4).

Independent kinetic support comes from the ectotherm thermal-manipulation datasets of §2.4.3, whose Arrhenius lifespan slopes bracket the mammalian SSP Pace coefficient. The point is not the identity of any single dataset but the convergence: intercepts differ because Scope and Stability are phylum-specific, while the temperature sensitivity converges because the underlying deamination chemistry is shared.

The circuit analogy yields a further prediction: rising impedance slows charge–discharge kinetics, analogous to an increased recovery time constant. As *R_int_*grows, electron current falls, NAD^+^ regeneration falls with it, the CoQ pool becomes increasingly reduced under demand, and Δ*ψ*-dependent work loses reserve. Declining Δ*ψ* partially relieves back-EMF on the electron circuit but simultaneously deprives NNT of the driving force for NADH-to-NAD^+^ recovery, so no internal redistribution satisfies all three circuits at once (§2.6.4). Reserve compression therefore manifests clinically as metabolic inflexibility and delayed post-stress recovery, consistent with the Strehler–Mildvan relation between reserve erosion and mortality acceleration (§2.5; Supplementary Note S3.1a).

### 3.9 Causal prediction and experimental tests

MSTA generates a sharp empirical discriminator: an intervention should rescue only the gate it relieves. LbNOX provides the cleanest current test of the carbon/NAD^+^ gate — a bacterial NADH oxidase that recycles NAD^+^ without generating ATP or pumping protons. Hepatic expression in mice lowers NADH/NAD^+^ and reverses insulin resistance and hypertriglyceridemia^50^; in *Drosophila*, adult-onset expression extends lifespan, with muscle-specific targeting conferring the strongest protection^77^ — consistent with skeletal muscle as the principal scope-discharge tissue, where metabolic shifts precede clinical sarcopenia in primates^78^. Because LbNOX restores NAD^+^ while contributing nothing to proton pumping or CoQ-dependent electron transfer — and, by diverting NADH from Complex I, may even lower respiratory flux and Δ*ψ*m — improvement despite this decoupling identifies carbon-circuit redox pressure as a sufficient and, in some contexts, rate-limiting branch of the aging phenotype.

This differs from yeast Ndi1, which bypasses Complex I by passing NADH electrons to CoQ and can extend fly lifespan^79^: because the CoQH_2_ it generates still supports Complex III/IV proton pumping, its effect is compatible with multiple causal readings and does not isolate the carbon contribution. LbNOX is more discriminating because it separates carbon-circuit rescue from electron and proton work; broader NAD^+^/NADH rebalancing mitigates age-related phenotypes on the same logic^80^.

A second, already-realized discriminator separates the CoQ gate from the NAD^+^ gate. In cells lacking Complex III, restoring CoQ-pool reoxidation with the alternative oxidase (AOX) rescues DHODH-driven pyrimidine synthesis, oxidative TCA flux, and proliferation, whereas regenerating NAD^+^ with LbNOX does not^82^. Demand-revealed CoQ-gate failure is therefore relieved by oxidized-CoQ availability, not by NAD^+^ status—the mirror image of the carbon-gate result above, where LbNOX suffices. LbNOX and CoQ reoxidation thus partition the two reductive gates, just as the Δ*ψ*-dependent predictions below partition the proton gate from both. Mitochondria-targeted catalase completes the set from the opposite direction: rather than relieving a gate, it contains the H_2_O_2_ product of electron-circuit saturation once the glutathione vent is overwhelmed, and its signature—delayed onset with no change in mortality slope—marks it as product containment, not conductance restoration.

The same logic extends to the other gates as a perturbation set. If electron-circuit saturation limits proliferation through DHODH, uridine should rescue pyrimidine-limited proliferation without restoring conductance; if the carbon arm limits the same cells through aspartate or one-carbon supply, aspartate or formate should rescue only that arm, and combined uridine-plus-aspartate rescue would confirm the dual checkpoint of §3.7. Mild uncoupling, by relieving back-EMF, may improve NADH pressure and reduce RET-linked ROS but should worsen high-threshold Δ*ψ*-dependent work if pushed too far. Only restoration of upstream conductance should improve all three readouts at once — NAD^+^ regeneration, oxidized-CoQ availability under demand, and Δ*ψ*-dependent recovery kinetics — whereas a bypass therapy should improve only the branch it bypasses.

Two predictions follow symmetrically and provide the strongest falsification test. Phenotypes driven by cytosolic NADH/NAD^+^ elevation — with GAPDH as the canonical sensor and the age-related contraction of cortical aerobic glycolysis^81^ as the most direct in vivo readout — should be rescued by LbNOX. Genuinely Δ*ψ*-dependent phenotypes — steroidogenic decline (CYP11A1, StAR import) and impaired Δ*ψ*-dependent inner-membrane import through TIM23/PAM after TOM entry — should not, because no NAD^+^-restoring bypass resolves the proton-circuit trilemma described in §2.6.4.

The discriminators above are interventional; the theory also yields an observational prediction that needs no perturbation. The cleanest test of a reductive-poise mechanism is a near-equilibrium redox reader, which reports the free [NADH]/[NAD^+^] ratio directly, independent of both the size of the nicotinamide pool and its position in the network; the *β*-hydroxybutyrate:acetoacetate couple is the canonical one. A fasting fuel generated in hepatic mitochondria from fatty-acid-derived acetyl-CoA, it is interconverted only by the NAD-specific, near-equilibrium enzyme BDH1, so its ratio is pinned to the matrix free [NADH]/[NAD^+^] — the relationship Williamson, Lund & Krebs (1967) used to show the matrix is far more reduced than the cytosol (free NAD^+^/NADH ≈ 8 versus ≈ 725). The prediction is therefore specific and falsifiable — rising OXPHOS impedance should drive the ratio up with age — and, because a near-equilibrium ratio is fixed by poise rather than absolute concentration, it discriminates a genuine reductive shift from a simple contraction of the NAD pool (which would slow flux while leaving the ratio unmoved), making the scarcity of longitudinal ketone-ratio data not a weakness of the theory but the experiment it most invites. As an observed companion, 4-hydroxynonenal adducts accumulate robustly with age across tissues (Zheng et al., 2005; Negre-Salvayre et al., 2008): generated by lipid peroxidation and cleared chiefly through NAD^+^-dependent ALDH2, this boundary metabolite can rise only when its redox-gated exit is throttled, and it is self-amplifying, since the adducts inactivate the upstream lipoate-dependent 2-oxoacid dehydrogenases (Humphries & Szweda, 1998). We rest on these two — an equilibrium reader of poise and a boundary accumulator of clearance failure — and deliberately not on the other Krebs-cycle intermediate ratios, which are confounded on several counts: their terms are gated on both sides, the contracting NAD(H) pool lowers flux without shifting poise, and anaplerotic input and cataplerotic drain set their steady-state levels independently of any single gated step — *α*-ketoglutarate being the clearest case, siphoned by transamination, the 2-oxoglutarate-dependent dioxygenases, and biosynthesis. (We likewise set aside lactate:pyruvate, whose age-related rise partly reflects a lactate-dehydrogenase A/B isoenzyme shift rather than redox state; Ross et al., 2010.)

### 3.10 Limitations

MSTA is a circuit-derived theory built from comparative scaling and mechanistic inference, and several limitations follow. Heavy-strand GC content is a structural proxy for mutational vulnerability, not a direct measurement of lesion velocity; direct measurements of mtDNA lesion accumulation, repair, and impedance change across species and tissues are needed. Body mass is an allometric proxy for Scope, not a measurement of cellular reserve: whole-organism VO_2_max/BMR is hard to standardize and conflates tissue quantity with intrinsic cellular reserve, so direct tissue-level measurements of mitochondrial density, cristae reserve, and maximal-to-basal conductance ratios would provide stronger tests. Consistent with the proxy interpretation, elite athletes achieve clear healthspan benefits but do not dominate supercentenarian demographics — exercise delays threshold-crossing without altering the cellular architecture that sets maximum lifespan.

#### Encoded chain reserve and the residual axes

The three-circuit framework names more variables than the SSP regression quantifies. The ∼31% of mammalian lifespan variance unexplained by *α*·ln(BM) + *β*·GC% − *γ*·Tb is structured rather than noise: two categories of circuit variables, both largely nuclear-encoded and therefore invisible to mtDNA-based regressors, plausibly account for most of it. The first sets *R_int,_*_0_ — the encoded chain quality at birth. The amino acid sequences of OXPHOS components are determined jointly by mtDNA (13 catalytic-core subunits) and nDNA (∼80 structural subunits, all assembly factors, and Complex II in its entirety). mtDNA GC% acts on *R_int,_*_0_ only indirectly via codon-biased amino acid composition; the dominant nuclear-encoded contribution is uncorrelated with any SSP regressor and is plausibly the largest single source of the residual. This residual contribution to *R_int,_*_0_ is distinct from the SSP GC% coefficient: GC% in the SSP regression is treated primarily as a lesion-rate modifier via replication-associated deamination vulnerability (above), while any contribution of base composition to encoded chain quality at birth is a secondary residual effect. The second modifies *dR_int_/dt* beyond the deamination-and-Arrhenius core — the chain-maintenance machinery. mtDNA polymerase proofreading fidelity sets a floor on de novo mutation rate independent of base composition; base excision repair directly counters the deamination channel underlying *β*·GC%; and protein quality control — the mitochondrial unfolded protein response, mitophagy, and mitochondrial biogenesis — reduces the lifetime burden of damaged chain components without slowing the underlying damage chemistry.

The revised three-gate model carries its own caveats. CoQ-gate saturation is inferred from circuit topology and steady-state redox logic; direct longitudinal measurements of tissue-specific CoQ redox state during aging are sparse, and because the gate is demand-revealed, resting CoQ measurements may mislead. Proton-circuit brownout is a threshold model, not a claim of uniform depolarization — average Δ*ψ* may appear preserved in bulk assays while high-threshold voltage-dependent work fails in specific mitochondrial subpopulations, so tests should measure function — TIM23/PAM import, steroidogenic stimulation, carrier exchange, recovery kinetics — rather than bulk membrane-potential dyes alone. Figure 7 maps gate engagement modes and downstream vulnerabilities; it is not a fixed chronological staging system, and the visible order of NAD^+^ compression, CoQ saturation, and Δ*ψ* brownout will depend on local demand, reserve, and measurement context. The NAD(P)(H)-proteome estimate has been curated to direct molecular evidence: Supplementary Table S8 retains 236 proteins that bind NAD(P)(H) as a redox cofactor across 16 fold families (Table S8a), with NAD^+^-consuming enzymes (Table S8b) and biosynthesis, transport, and sensor proteins (Table S8c) tabulated separately, and inclusion criteria, cofactor specificity, and fold assignment stated in the table; the estimate reflects current UniProt annotation and will refine as binding evidence accrues. Finally, the dual-gate distinction itself awaits prospective longitudinal validation with challenge assays in human cohorts.

The SSP model accounts for ∼69% of mammalian maximum-lifespan variance, leaving structured residual variation whose likely contributors — phylogenetic non-independence, nuclear-encoded OXPHOS quality, mitochondrial translation fidelity, repair and quality-control capacity, endocrine set-point, and ecological mortality — define the next layer rather than weakening the model. And the manifold interpretation does not imply that the fitted coefficients are selection gradients or marginal fitness values; they estimate local longevity elasticities within an evolved constraint space, and inferring fitness gradients would require age-specific fecundity and extrinsic-mortality data beyond the present analysis.

### 3.11 Relation to the energy resistance principle

MSTA was developed independently of, but converges with, the recently proposed energy resistance principle (ERP), which frames health and aging around energy resistance, éR, defined as an organism’s energy potential divided by the square of its electron flux^83^. The two frameworks agree on the inversion that motivates MSTA: the primary lesion of mitochondrial aging is loss of transformation capacity, not ATP depletion — impaired mitochondria reduce electron-flux capacity while resting ATP is frequently preserved — and reductive stress, as electrons back-flow onto NAD^+^, precedes oxidative stress as the proximal molecular consequence. MSTA’s NAD^+^ gate, its coupling of reductive to oxidative stress through reverse electron transport, and its reading of lactate and cataplerotic export as redox pressure valves are the circuit-level realization of these same observations. Convergence from an independent framework strengthens the shared premise; the contribution of MSTA is to render it quantitative, comparative, and mechanistically resolved.

Mitochondrial éR is the same quantity as the OXPHOS internal resistance defined in §2.6. Holding the driving force fixed at the thermodynamically clamped NADH-to-O_2_ redox span, the electrical power law and the relation éR = EP/f^2^ coincide, so MSTA is the voltage-clamped, lesion-driven, time-resolved specialization of the general principle. This clamping also removes an ambiguity in the general formulation, whose numerator simultaneously denotes a voltage-like accumulation of substrate and a power-like demand: by fixing the redox driving voltage and treating substrate supply and biosynthetic demand as separate loads, MSTA gives éR a single referent — rising series resistance in the electron-transport chain — and, critically, supplies its rate of change. Three features separate MSTA from a static state-variable account. First, the energy-resistance ratio is a snapshot with no reserve term and no traversable interval, whereas the MSTA lifespan law equates maximum lifespan with the conductance-reserve interval divided by the velocity of impedance accumulation (§2.6); its explicit capacity term — the body-mass-scaled conductance reserve, or Scope — has no counterpart in the general principle. Second, MSTA specifies the cross-species law for impedance accumulation, the Scope–Stability–Pace relation, in which Stability and Pace set the rate of impedance growth through the deaminable-base burden of the heavy strand and the Arrhenius kinetics of its loss. Third, MSTA partitions the consequences of rising resistance into three ordered gates — NAD^+^, CoQ, and Δ*ψ* — rather than a single reductive–oxidative axis, which is what converts an intervention such as LbNOX from a generic flux-restorer into a gate-specific discriminator (§3.9). The general principle’s own outlook names these as open problems — what constitutes a biological resistor, and how organisms with broad thermal ranges might link temperature, metabolism, and lifespan; MSTA answers the first with the thirteen mtDNA-encoded subunits as series resistors eroded by replication-associated deamination, and the second with the ectotherm kinetic validation of the Pace coefficient (§2.4.3).

Two points require distinction rather than assimilation. The general principle treats temperature as a downstream consequence of resistance, higher resistance dissipating more heat; MSTA assigns body temperature the opposite, upstream role — a primary driver of the lesion-generation rate through the Arrhenius dependence of hydrolytic deamination, with an apparent activation energy (∼73 kJ/mol) recovered independently by the within-species ectotherm temperature–lifespan slopes (§2.4.3). Temperature in MSTA sets how fast resistance accumulates; it is not a byproduct of resistance already present. Cancer requires a similar refinement. The general principle reads the Warburg state as one of high energy resistance, on the premise that transformation requires resistance; MSTA draws the distinction more finely. Proliferating cells do run at high resistance through the oxidative chain — consistent with the slow oxidative TCA flux measured in solid tumors — but they sustain biosynthesis precisely by relieving the gates that aging closes: regenerating NAD^+^ through cytosolic lactate export, sustaining pyrimidine synthesis through DHODH, and supplying citrate through reductive carboxylation. Aging and cancer therefore share elevated oxidative-chain impedance yet diverge at the dual nucleotide checkpoint (§3.7): aging closes it, imposing managed retreat from division, whereas transformation forces it open through the same bypasses MSTA proposes as conditional geroprotective levers — which is why those levers carry an intrinsic proliferative-risk ceiling that any conductance-sparing therapy must respect.

## Conclusion

Aging is managed retreat under shrinking metabolic scope. The first crisis is biosynthetic, not energetic. Rising mitochondrial impedance is read out through three asymmetric gates: constitutive carbon-circuit compression, demand-revealed electron-circuit saturation, and threshold-gated proton-circuit brownout. Together they explain graduality — analog throttling at distributed gates rather than catastrophic failure; synchrony — a single shared upstream lesion expressed everywhere at once; and syndrome clustering — tissue-specific valves and endocrine forks responding to one constraint; while preserving the central paradox of aging: basal ATP remains defended even as regeneration becomes unaffordable. The therapeutic mandate that follows is conductance-first — not to add fuel, scavenge exhaust, or block individual downstream valves, but to restore the throughput that makes biosynthesis, repair, and regeneration thermodynamically affordable once more.

## Methods

### Data sources

Mammalian body mass (BM), basal metabolic rate (BMR), and maximum lifespan (MLS) were obtained from the Human Ageing Genomic Resources AnAge database (build 14)^1^. Heavy-strand mitochondrial DNA guanine–cytosine content (GC%) was compiled from the MitoAge database^2^. Core body temperatures (Tb) for mammals and birds were taken from Clarke & Rothery^18^ and supplementary sources detailed in Supplementary Note S1. Avian entries were integrated from the same primary sources. Teleost, chondrichthyan, chondrostean, and holostean records were compiled from AnAge with manual curation of ambient temperature from FishBase and primary literature. Body temperature, GC%, and longevity data used in this study are consolidated in Supplementary Table S1 (AnAge_Based_Table.csv).

### Species inclusion criteria

Species were included in the Scope–Stability–Pace (SSP) regression if all three predictors (BM, GC%, Tb) and MLS were available (n = 379 mammals for the main fit; n = 182 birds; n = 247 teleosts used in Fig. 5B). For allometric plots not requiring GC% or Tb, broader species sets were used where indicated in figure legends (up to n = 1,023 mammals and n = 425 fish). Domestic and captive-only species were retained only where AnAge records MLS from wild or minimally managed populations, to limit anthropogenic longevity inflation.

### SSP regression

The log-linear Scope–Stability–Pace model ln(MLS) = *α*·ln(BM) + *β*·GC% − *γ*·Tb + c was fitted by ordinary least squares. Reported coefficients are point estimates with coefficient of determination R^2^ and asymptotic p-values; 95% confidence intervals were obtained analytically from the regression covariance matrix. Partial correlations (Fig. 3) were computed from residualised variables after sequential regression on the confounder(s) indicated. The main analysis is unadjusted for phylogenetic non-independence.

### Phylogenetic sensitivity analysis

To test whether the SSP coefficients reflect shared ancestry rather than independent evolution, we re-fitted the model using phylogenetic generalized least squares (PGLS), which lets regression residuals covary in proportion to shared evolutionary history. We used 100 time-calibrated mammalian trees sampled from the posterior of the Upham et al. (2019) supertree (VertLife.org); 368 of the 379 SSP species were present on every tree and were used for all fits. On each tree we fitted three residual-covariance models — Brownian motion, Pagel’s *λ* (which estimates the strength of phylogenetic signal), and Ornstein–Uhlenbeck (which adds a restoring force toward a trait optimum) — with the phylolm package (Ho & Ané 2014), reporting medians and 95% ranges across the 100 trees. OLS is retained as the primary model for interpretability; full results are in Supplementary Note S2.3.

### Ectotherm kinetic validation

Five published ectotherm temperature–lifespan datasets were re-analysed by fitting ln(MLS) as a function of 1000/T_K (Arrhenius form) to extract apparent activation energies: Drosophila melanogaster (ref. 26), Musca domestica (ref. 27), Caenorhabditis elegans (ref. 28), Daphnia magna (ref. 29), and Nothobranchius rachovii (ref. 30). For Daphnia, slopes were computed from the two clones in which age-dependent (Gompertz) mortality dominates the signal; see Supplementary Note S1.1 for the full treatment.

#### Software

Data curation, regression, and Arrhenius fits were implemented in R version 4.x (R Core Team). Analysis scripts and compiled datasets are provided as Supplementary Material.

## Data availability

All primary data supporting the SSP analyses are deposited with this manuscript as supplementary files (Supplementary Table S1; AnAge_Based_Table.csv). Source databases are publicly available: AnAge (https://genomics.senescence.info/species/), MitoAge (http://mitoage.org/), and FishBase (https://www.fishbase.se/). R analysis scripts are provided as Supplementary Material.

## Code availability

All R code used for the analyses and figures in this study is provided as Supplementary Material.

## Author contributions

G.L. conceived the Metabolic Scope Theory of Aging, curated and analysed the data, and drafted the manuscript. J.L. clarified the behaviour of rising impedance in electrical circuits, which G.L. then mapped onto the bioenergetic system, and I.S. read the manuscript, raised questions, and helped refine the electric-circuit analogies. Y.A. worked through the Krebs-cycle reasoning with G.L. N.S. critically read the manuscript, identified flaws, and suggested improvements. Y.G. and G.S. contributed to interpretation of the endocrine and mitochondrial mechanisms and critically revised the manuscript. All authors read and approved the final version.

## Acknowledgements

The authors thank colleagues at the Department of Endocrinology, Tel Aviv-Sourasky Medical Center, for helpful discussions.

