## Supplementary information for "The Metabolic Scope Theory of Aging: Rising Mitochondrial Impedance Compresses Metabolic Reserve to Constrain Lifespan"

#### Contents


- Supplementary Reader's Guide
- Supplementary Methods (M1–M3)
- Supplementary Notes (S1–S7)
- Supplementary Tables (S1–S9)
- Supplementary References


#### Supplementary Reader's Guide

This Supplementary Information file provides methodological details, robustness checks, extended mechanistic derivations, and complete tables referenced in the Main Text.

Suggested navigation:

- Quantitative SSP model, variable definitions, and robustness: Supplementary Methods M1–M3 and Supplementary Note S2.
- Circuit model and metabolic impedance framing: Supplementary Notes S3 and S5.
- Consilience matrix (precursor/product dissociations and ratio biomarkers): Table S4.
- $NAD^+$/NADPH Fork framework and pressure-valve logic: Supplementary Note S7 and Table S6.

This supplement includes comparative methods and robustness checks, extended circuit-level derivations, a cross-pathway consilience matrix of precursor/product dissociations and ratio biomarkers, and the $NAD^+$/NADPH Fork framework whose directional shifts generate specific aging phenotypes.

### Supplementary Methods

#### M1. Comparative dataset assembly and quality control

We curated a vertebrate comparative dataset comprising, where available: (i) maximum lifespan (MLS), (ii) adult body mass (BM), (iii) typical body temperature (Tb; core temperature for endotherms and typical environmental/active temperature for ectotherms), (iv) mtDNA sequence-derived base-composition features, and (v) aerobic-capacity indices (VO2max, maximal metabolic rate [MMR], resting/basal metabolic rate [RMR/BMR], and factorial aerobic scope [MMR/BMR]).

In the working dataset used for the primary SSP fits, we used 379 mammals with complete data on all four variables as the primary cohort; birds (n = 182) and teleost fishes (n = 247) were analyzed as independent cross-class tests rather than pooled into the primary fit. Lifespan values were drawn from the AnAge database and the primary literature; body mass and temperature estimates were drawn from physiological compilations and species accounts. For ectotherms, where a single 'body temperature' is not well defined, experimental temperature or typical environmental/active temperature was used as a proxy.

Species were excluded where MLS or Tb estimates were judged highly uncertain, where adult BM estimates were inconsistent across sources, or where mtDNA sequence records were incomplete or low confidence. Sample size varies across analyses depending on variable coverage and on the specific model being evaluated.

All variables were harmonized to consistent units and transformed where appropriate (log transforms for MLS and BM). Outlier screening focused on implausible lifespan records, inconsistent adult mass values, and obviously mismatched mtDNA records (e.g., partial genomes or mis-annotated accessions).

#### M2. Scope metrics and proxies

Metabolic scope is defined as reserve capacity: the gap between maximal sustainable power output and basal maintenance requirements. In comparative physiology, this reserve is captured by maximal oxygen consumption (VO2max or MMR) relative to resting consumption (BMR/RMR).

When maximal and resting metabolic measurements were available, Scope was quantified as factorial aerobic scope (MMR/BMR) or as a VO2max-derived reserve index. These measurements are sparse across vertebrates, so to enable full-coverage analysis we additionally used a calibrated Scope proxy. The proxy was trained within the subset of species with direct aerobic-capacity measurements and then applied out-of-sample when Scope_measured was unavailable.

Sensitivity analyses evaluated SSP robustness to alternative Scope constructions, including: (i) use of body-mass-only proxies, (ii) replacement of factorial scope with

VO2max-derived indices, (iii) clade-stratified calibrations to minimize extrapolation error, and (iv) exclusion of proxy-based entries to verify that key coefficient signs persist when only direct Scope measurements are used.

#### M3. mtDNA sequence processing and Stability metrics

Complete mitochondrial genomes were retrieved from GenBank and related sequence repositories. Base composition was computed on the heavy-strand convention to capture replication-associated single-strand exposure during asynchronous replication.

Global Stability metrics included whole-genome GC% and heavy-strand guanine fraction ($G_H$). Gene-level Stability metrics included per-gene GC%, heavy-strand base fractions, and strand-asymmetry indices (e.g., GC skew, AT skew). For additional resolution, genomes were partitioned into functional elements (protein-coding genes, rRNAs, and tRNAs), and where relevant, structural partitions (e.g., tRNA stems vs loops) were used to evaluate whether stability signals concentrate in base-paired regions.

To quantify replication-associated positional effects, genes and windows along the heavy strand were mapped by genomic distance from the origin of heavy-strand replication (OriH). Heavy-strand thymine accumulation ($T_H$) and related compositional gradients were used as proxies for replication-exposure 'scarring'. These positional analyses operationalize the danger-zone concept in which single-stranded exposure time during asynchronous replication increases mutational pressure.

Sequence-processing scripts computed: global mtDNA GC%; strand-specific composition (heavy vs light); distance of each base along the heavy strand from OriH; decile-wise composition across the control region; and per-gene compositional summaries. Intermediate outputs can be provided as supplemental data files to support reproducibility.

### Supplementary Notes

#### Supplementary Note S1. Arrhenius mapping of the Pace term

##### S1.1 Converting the fitted temperature coefficient to an effective activation energy

In endotherms, SSP fits yield a Pace coefficient on the order of −0.081 per degree C (Tb in degrees C). Interpreting MLS as inversely related to an effective rate of scope erosion, an Arrhenius mapping can be used as a consistency check. For a process with rate r(T) proportional to $\exp(-E_a/(R\cdot T))$, ln(r) changes with temperature as $d\ln(r)/dT = E_a/(R\cdot T^2)$. If MLS is proportional to 1/r, then $d\ln(MLS)/dT = -E_a/(R\cdot T^2)$. Evaluated at $T \approx 310$ K (approximately 37°C), a slope of $-0.081\ K^{-1}$ corresponds to an effective activation energy $E_a \approx 0.081\cdot R\cdot T^2 \approx 65$ kJ/mol. This magnitude is compatible with common hydrolytic and deamination chemistry, supporting the interpretation that the Pace term reflects generic thermally activated kinetics rather than organism-specific metabolic programming.

##### S1.2 Why Stability must interact with Pace

The Stability term is encoded by mtDNA base composition, especially heavy-strand guanine (G_H). The mechanistic claim is not that guanine is universally 'good', but that in a replicated, intermittently single-stranded genome, base composition shifts the thermodynamic cost of strand separation and the propensity for single-strand lesions. At higher temperatures, small differences in duplex stability can translate into large differences in time spent in exposed conformations ('breathing'), amplifying replication-associated pressure. Thus, Stability should matter most in warm regimes, whereas in cold regimes, breathing is suppressed and the dominant cost of high-G content may become replication or structural burden.

##### S1.3 Temperature inversion as a mechanistic validation

The above logic yields a strong qualitative prediction: the sign of the Stability term should attenuate and can invert in cold-adapted ectotherms. This is difficult to explain under theories in which base composition is merely a neutral correlate. In the dataset, cold-water fishes show an attenuated or negative association between G_H (or GC%) and longevity, consistent with the predicted inversion. We recommend evaluating this interaction explicitly using models with a Stability × Tb term and reporting the sign of the marginal Stability effect across the ectotherm temperature range.

#### Supplementary Note S2. Robustness checks and sensitivity analyses

This note presents the robustness checks and sensitivity analyses underlying the main-text SSP results. Because comparative datasets are heterogeneous, robustness is assessed along three axes: (i) sensitivity to influential taxa, (ii) sensitivity to variable definition and measurement error, and (iii) sensitivity to phylogenetic non-independence.

##### S2.1 Influence and outlier diagnostics

We recommend reporting leverage and Cook's distance for the primary SSP models, and repeating key fits after excluding high-leverage taxa. Particular attention should be paid to clades known to be longevity outliers (e.g., bats and subterranean rodents) to ensure that SSP captures broad structure rather than being driven by a few iconic points.

##### S2.2 Model comparisons against plausible alternatives

To demonstrate that SSP is not a disguised rate-of-living effect, competing models should include: (i) BM-only models, (ii) BM plus basal metabolic rate (BMR) models, (iii) mass-specific metabolic rate models, and (iv) temperature-only models. The key qualitative expectation is that resting flux should not replace Scope, and that Stability should remain predictive even when size and temperature are included. Models are compared by variance explained and information criteria (AIC/BIC), and predictive generalization is assessed by leave-one-clade-out cross-validation across major vertebrate classes.

#### S2.3 Phylogenetic robustness of the SSP regression

##### *The problem: species are not independent data points*

Ordinary least squares (OLS) regression assumes that every data point contributes independent information. In comparative biology this assumption is violated: a mouse and a rat share recent common ancestry, so their similar body masses, lifespans, and mtDNA compositions are partly inherited from the same ancestor rather than independently evolved. Treating them as two fully independent observations inflates the effective sample size and can make a spurious correlation appear statistically significant. Any comparative regression — including the SSP model — must therefore be tested for robustness to this phylogenetic non-independence.

##### *What phylogenetic generalized least squares (PGLS) does*

PGLS is a regression method that accounts for shared ancestry by modifying the error structure of the model. In OLS, residuals are assumed to be uncorrelated across species; in PGLS, residuals are allowed to covary in proportion to shared evolutionary history. Formally, OLS minimizes $(y - X\beta)^T(y - X\beta)$, treating all residuals equally, while PGLS minimizes $(y - X\beta)^T V^{-1}(y - X\beta)$, where V is a variance–covariance matrix derived from the phylogenetic tree. Species that share a long common history receive higher off-diagonal entries in V, so their residuals are expected to be more similar. The resulting regression coefficients are weighted to extract genuinely independent evolutionary signal rather than counting the same ancestral information multiple times.

The key diagnostic is not whether the coefficients change — some change is expected — but whether their signs and approximate magnitudes survive phylogenetic correction. If a positive OLS coefficient becomes negative under PGLS, the relationship may have been an artifact of clade-level confounding. If it remains positive but shrinks only modestly, the direction of the effect is genuine and the underlying biology is preserved.

##### *Three models of trait evolution: BM, Pagel's λ, and Ornstein–Uhlenbeck*

PGLS requires a model for how traits evolve along the branches of a phylogeny. This model determines the structure of V — in other words, how much phylogenetic covariance to expect between any two species. We evaluated three standard options:

**Pure Brownian motion (BM).** Trait values drift randomly over evolutionary time, accumulating variance proportional to elapsed time. The covariance between two species equals the shared path length from the root of the tree to their most recent common ancestor. BM imposes the full, unattenuated phylogenetic covariance structure on the residuals.

**BM with Pagel's λ.** A one-parameter generalization of BM in which all off-diagonal entries of V are scaled by $\lambda \in [0, 1]$. $\lambda = 0$ recovers OLS (no phylogenetic signal); $\lambda = 1$ recovers pure BM; intermediate values correspond to phylogenetic signal weaker than BM predicts. Estimating λ from the data lets the analysis find the degree of phylogenetic correction best supported by the residuals.

**Ornstein–Uhlenbeck (OU).** This model adds a restoring force: trait values are pulled toward an optimum, so evolutionary "memory" decays exponentially over time rather than accumulating indefinitely. The key parameter is the selection strength $\alpha_{OU}$, which determines the phylogenetic half-life $t½ = \ln 2 / \alpha_{OU}$ — the time required for the evolutionary correlation between two lineages to decay by half. When t½ is much shorter than total tree depth, OU approaches OLS; when t½ is long, OU approaches BM. OU is the canonical model for traits under stabilizing selection within ecological niches.

#### *Dataset, phylogeny, and implementation*

We evaluated the SSP regression $\ln MLS = \alpha \cdot \ln BM + \beta \cdot GC\% - \gamma \cdot Tb + c$ using the primary mammalian dataset restricted to species with complete data on all four variables ($n = 379$). Species-level time-calibrated phylogenies were obtained from the Upham et al. (2019) mammalian supertree (VertLife.org, "MamPhy_BDvr_Completed_5911sp_topoCons_NDexp"), with a sample of 100 trees drawn from the posterior distribution to account for phylogenetic uncertainty. After intersecting the tip set of each tree with the SSP species list, 368 species were present on every tree and were used consistently across all 100 fits. The total depth of the mammalian tree is approximately 200 million years (Myr).

For each tree we fitted three models — BM, Pagel's λ, and OU with fixed root — using the phylolm R package (Ho & Ané 2014), which implements the Ho–Ané linear-time algorithm for Gaussian trait models. Coefficients, log-likelihoods, AIC values, and the fitted λ or $\alpha_{OU}$ were recorded for each tree, and we report medians and 95% ranges (2.5th–97.5th percentiles) across the 100 posterior samples. Because all three models share the same regression design matrix and differ only in the residual covariance structure, their log-likelihoods and AIC values are directly comparable.

#### *Results*

**Coefficient recovery is consistent across models.** All three phylogenetic models recovered the OLS sign pattern and approximate magnitudes of the SSP coefficients. Body mass, GC%, and body temperature retained their expected effect directions (positive, positive, negative) in 100/100 posterior trees under every model. The point estimates under BM, λ, and OU were mutually consistent to within small differences (Table S2).

**Pagel's λ ≈ 0.95, not at the boundary.** Across the 100 trees, λ converged to a median of 0.948 (95% range 0.940–0.960), well below the upper boundary of 1. This indicates substantial but not saturated phylogenetic signal in the residuals: closely related mammals do share correlated SSP residuals, but less so than a pure Brownian-motion drift along 200 Myr of tree depth would predict.

**The λ model is preferred over BM and OU.** Under AIC, the λ model outperforms both alternatives on every tree: median ΔAIC(BM − λ) = +53, median ΔAIC(OU − λ) = +23. In log-likelihood terms, λ beats BM by a median of +27 units per tree (95% range +18 to +113) and beats OU by a median of +12 units per tree (95% range +4 to +72). OU in turn beats BM by a median of +16 log-likelihood units per tree (95% range +12 to +41; $\chi^2 > 24$, 1 d.f., $p < 10^{-6}$ on every tree). All three comparisons are thus decisive in the expected direction: pure BM is rejected, and the Pagel's λ model is the best-fitting member of the family.

**The OU phylogenetic half-life is long but finite.** The median OU selection strength was $\alpha_{OU} = 0.018$ Myr$^{-1}$ (95% range 0.015–0.033), corresponding to a phylogenetic half-life $t½ \approx$ 39 Myr (95% range 21–45 Myr). This is a sizable fraction of the 200-Myr tree depth, indicating that evolutionary correlations among SSP residuals decay on a timescale comparable to major mammalian diversification rather than being confined to the shallow tips of the tree.

#### *Interpretation*

Three conclusions follow, and each strengthens rather than weakens the SSP claim.

First, the SSP coefficients are **robust to phylogenetic correction across a family of covariance models spanning BM, a freely estimated λ intermediate, and OU.** Signs are preserved in 100/100 posterior trees under every model. Magnitudes shrink under correction — by roughly half for β (GC%) and γ (Tb), only slightly for α (ln BM) — but the shrinkage is itself a benign signal: it reflects the fact that deep-clade structure (especially the endotherm body-temperature split) carries some of the univariate variance, leaving less to the within-clade regression once phylogeny is accounted for. The residual information is still sufficient to drive all three coefficients away from zero under every model tested.

Second, the fact that **Pagel's λ ≈ 0.95 rather than 1 or 0** indicates genuine phylogenetic signal in SSP residuals without saturation. This is the expected pattern for life-history traits under stabilizing selection: correlations persist across related lineages but decay more rapidly than unconstrained Brownian drift would imply. The corresponding OU phylogenetic half-life of ~39 Myr — neither short enough to collapse onto OLS nor long enough to approach BM — tells the same story in a different parameterization.

Third, the **model ranking (λ > OU > BM)** places the best-supported covariance structure in the middle of the BM–OLS continuum. This is the regime in which phylogenetic correction has genuine work to do but does not overwhelm the regression signal. The convergence of BM, λ, and OU on the same qualitative answer — with BM the most conservative and λ/OU recovering coefficients closer to OLS — means the SSP relationships are neither artifacts of deep-clade confounding (which BM would have exposed by driving coefficients to zero) nor artifacts of a spuriously unstructured error term (which $\lambda \approx 0$ would have exposed by reproducing OLS exactly).

In summary, the SSP regression is **robust** under every standard phylogenetic model, with the best-fitting model (Pagel's $\lambda \approx 0.95$) recovering $\alpha = +0.138$, $\beta = +0.053$, $\gamma = +0.039$ across all 100 posterior trees. We report OLS as the primary model in the main text for interpretability, with λ-PGLS and OU-PGLS as phylogenetic robustness checks.

#### S2.4 Measurement error and variable harmonization

Temperature is well-defined for endotherms but more variable for ectotherms. We therefore recommend (i) using multiple temperature proxies, (ii) stratifying ectotherms by habitat thermal regime, and (iii) reporting how the Stability effect changes across temperature bins. Similarly, where measured aerobic scope is sparse, report calibration diagnostics for any Scope proxy and show that SSP conclusions are not driven by the proxy construction.

### Supplementary Note S3. Circuit model details and derived predictions

This note provides a more explicit circuit formulation of oxidative phosphorylation and the mapping from circuit parameters to predicted aging phenotypes. The aim is to clarify the minimum assumptions needed for the high-voltage/low-current drift and the rate parity of the two electron-entry routes.

#### S3.1 Minimal Thevenin model and maximum power

Represent the respiratory chain as an effective voltage source V_redox in series with internal resistance R_total feeding an effective load R_load. Current is $I = V\_redox / (R\_total + R\_load)$. Power delivered to the load is $P\_load = I^2 \cdot R\_load = (V\_redox^2 \cdot R\_load) / (R\_total + R\_load)^2$. The maximum power theorem gives P_max at R_load = R_total, yielding $P\_max = V\_redox^2 / (4 \cdot R\_total)$. Therefore, even modest increases in R_total can cause a large collapse in maximal deliverable power, while allowing basal power to be maintained by increasing R_load sharing or by demand suppression.

#### S3.1a Six circuit-predicted consequences of rising OXPHOS impedance

The Thevenin model (S3.1) reduces the respiratory chain to an effective voltage source V_redox in series with internal resistance R_total feeding a load R_load. This minimal representation yields six quantitative consequences of rising R_total that map onto distinct aging phenotypes (Table S3). Crucially, these are not separate phenomena requiring separate explanations—they are mathematically connected outputs of a single underlying lesion.

**1. Ohm's law: high-voltage, low-current drift**

As R_total rises, current $I = V\_redox/(R\_total + R\_load)$ falls while the upstream reduced pool (NADH) accumulates. Membrane potential ($\Delta\psi$) can paradoxically increase at rest because proton pumping continues at reduced flux against a diminished load. The system is overcharged but under-supplied—the defining signature of impedance-driven aging. This drift is distinct from substrate limitation: fuel is abundant, but electrons cannot traverse the chain efficiently. The clinical correlate is that isolated measurements of $\Delta\psi$ or resting ATP may appear normal even when maximal biosynthetic throughput has already substantially declined.

**2. Joule dissipation: inflammaging as wasted power**

In a resistive circuit, power dissipated as heat is P_waste = $I^2 \times R$. When R_total rises 4-fold while the organism maintains basal current at ~50% of youthful levels:

P_waste $\propto (0.5\ I_0)^2 \times 4R_0 = 1.0 \times I_0^2 R_0$

The system produces equivalent dissipative losses to youth despite halved electron flux. In mitochondria, this dissipation appears as electron leakage onto $O_2$ and as thermal waste. The prediction is specific: inflammaging is intrinsic to high-impedance operation, not an independent pathology. It also explains why antioxidant supplementation (addressing the exhaust) shows limited lifespan benefit while exercise (reducing effective R by increasing mitochondrial content, i.e. adding parallel conductance paths) succeeds: exercise addresses the impedance, antioxidants address the symptom.

**3. Kirchhoff's current conservation: obligate ROS**

Total electron input must equal total output. If I_forward (productive flux through Complexes III and IV) is throttled by rising R, then I_leak (non-productive electron escape) must rise to absorb the difference. This is not stochastic damage but an obligate consequence of current conservation. The main leak sites—the Complex I FMN site and the Q-binding pocket—emit superoxide in proportion to the degree of forward-flow throttling. Importantly, this leak is constitutive: it operates even at rest, whenever NADH/$NAD^+$ is elevated. The leak does not require high flux or high metabolic demand—it is a property of the backed-up redox state itself.

The distinction from Consequence 2 is subtle but important: Joule dissipation describes the energetic cost per electron traversing the chain (proportional to $I^2R$), while Kirchhoff conservation describes the fraction of total electron flux that is diverted to non-productive leak (proportional to R_total / [R_total + R_forward]). Both predict rising ROS, but through different mechanisms—one is per-electron inefficiency, the other is current re-routing.

**4. Impedance mismatch: upstream pool depletion**

Maximum power transfer from source to load occurs when R_source = R_load. In young mitochondria, R_total $\ll$ R_load (the chain has excess conductance relative to demand), and power delivery is efficient. As R_total rises, the mismatch grows: source impedance increasingly dominates the circuit, and the $NAD^+$ pool is trapped in reduced form.

The immediate consequence is not ATP shortage but cofactor depletion. $NAD^+$ is the limiting substrate for sirtuins (SIRT1–7), PARPs, CD38, and the >400 other $NAD^+$-dependent enzymes catalogued in the human proteome. Their activities depend on the absolute concentration of $NAD^+$, not on the NADH/$NAD^+$ ratio per se (though the two are coupled). Impedance mismatch thus starves the cell's epigenetic maintenance (sirtuin-mediated deacetylation), DNA repair (PARP), and signaling (CD38/cADPR) machinery before it starves the ATP synthase.

This ordering is critical: the impedance mismatch predicts that the first casualties of rising R_total are the $NAD^+$-consuming housekeeping enzymes, not energy production. Empirical support comes from the observation that $NAD^+$ supplementation (NMN, NR) can restore some age-related functions even when the underlying impedance lesion persists—consistent with a depleted-pool bottleneck rather than a broken-enzyme bottleneck.

**5. RC time constant: metabolic inflexibility**

The inner mitochondrial membrane functions as a capacitor (C) that stores charge as Δψ. The time constant for charging and discharging this capacitor is τ = R_total × C. As R_total rises, τ increases: the system charges more slowly after a discharge event (e.g., an ATP demand transient) and discharges more slowly after a reduction in demand.

Clinically, this predicts metabolic inflexibility—the well-documented inability of aged tissues to switch rapidly between fuel substrates, to recover $VO_2$ after exercise, or to suppress glucose oxidation during fasting. The delayed substrate switching observed in aged skeletal muscle, the prolonged post-exercise oxygen debt in elderly subjects, and the impaired post-prandial metabolic response all fit the prediction of a system with an elevated RC time constant.

Quantitatively, if R_total doubles while C remains approximately constant, τ doubles. A mitochondrion that formerly recovered full Δψ within 100 ms of an ATP burst now requires 200 ms. At the tissue level, this compounds across millions of mitochondria and manifests as the seconds-to-minutes-scale delays measured in whole-organism metabolic flexibility assays.

**6. Maximum power theorem: non-linear frailty**

The maximum deliverable power is $P\_max = V\_redox^2 / (4 \times R\_total)$. P_max falls as 1/R_total, meaning that each incremental rise in impedance produces a proportional loss in maximal power. However, because organisms operate well below P_max under resting conditions, the functional consequence is felt primarily in the maximal-demand regime: the ceiling on what the organism can sustain drops.

The mortality consequence of this declining maximal power—an exponentially contracting reserve, $V(t) \approx V_0 \cdot \exp(-kt)$, feeding the Strehler–Mildvan/Gompertz hazard—is derived in the main text (§2.5).

**Multiplicative power degradation and the proton-circuit brownout**

The six consequences above treat R_total as the sole degrading variable. In reality, three power-delivery variables degrade concurrently as impedance accumulates.

(a) Effective driving voltage (V_redox) depends on which electron-entry mode dominates. When the respirasome carries most flux, V_redox ≈ 1.14 V (NADH → $O_2$). As Complex I impedance rises and the FAD-linked CoQ entry increasingly shares the load, the effective V_redox drops toward ~0.77 V ($CoQH_2$ → $O_2$).

(b) Proton stoichiometry (n_H) falls from ~10 $H^+$ per $2e^-$ (respirasome path) toward ~6 $H^+$ per $2e^-$ (small supercomplex path) as flux shifts.

(c) R_total itself rises as lesions accumulate in mtDNA-encoded subunits.

Power delivered to the proton circuit scales as P ∝ V_redox × n_H / R_total. A numerical example: if V_redox falls from 1.14 to 0.90 V (21% decline), n_H from 10 to 8 (20% decline), and R_total doubles—all plausible mid-trajectory changes—the product falls to (0.90/1.14) × (8/10) × (1/2) = 0.79 × 0.80 × 0.50 = 0.32. A 68% power loss from three modest individual changes. This multiplicative compression explains why proton-circuit-gated loads begin failing well before any single parameter reaches a critical threshold.

**Voltage sag and proton-circuit load shedding.** At rest, demand (R_load) is high relative to R_total, and the proton circuit's voltage (Δψ) appears nearly normal. Under any demand transient—a burst of mitochondrial protein import, a steroidogenic pulse, a translation spike—R_load drops and R_total dominates the circuit. The resulting voltage sag is:

ΔV_sag = V_redox(t) × R_total(t) / [R_total(t) + R_load]

As V_redox drops and R_total rises, the same cellular task produces progressively deeper voltage sags and slower recovery. Proton-gated loads differ in their Δψ threshold requirements. High-threshold loads—mitochondrial protein import (TIM/TOM), Fe–S cluster export (ABCB7), and steroidogenic CYP reactions (CYP11A1 cholesterol side-chain cleavage)—require sustained Δψ above specific minima. When voltage-sag depth exceeds their tolerance, they fail: first intermittently under demand transients, then chronically as resting Δψ itself declines. Low-threshold loads (ATP synthase, passive proton leak) ride through the sag and are preserved until much later.

This thermodynamic triage requires no regulatory coordination. Each load's flux is set by its intrinsic sensitivity to delivered Δψ, so the shedding order is determined by circuit architecture rather than by sensing programs. Buttgereit & Brand (1995) demonstrated exactly this hierarchy experimentally in thymocytes under progressive respiratory inhibition: biosynthesis failed before housekeeping ion cycling and proton leak, with each load showing its own characteristic sensitivity to declining respiratory supply. Aging engages the same hierarchy on a longer timescale.

#### S3.1b Rate parity of the two electron-entry routes

The main text states that the NADH and CoQ electron-entry routes, despite carrying unequal mutational targets, accumulate impedance at the same fractional rate. This note derives that parity and shows why the resulting asymmetry between the $NAD^+$ and CoQ gates is one of threshold, not of aging rate.

**Setup.** Treat each entry route as a series chain of mtDNA-encoded conductive elements. The respirasome route (I–$III_2$–IV) places $n_1 \approx 11$ mtDNA-encoded subunits in series; the CoQ route (Complex II, ETFDH, DHODH, and mGPDH feeding $III_2$–IV) places $n_2 \approx 4$. Let each intact subunit contribute a baseline series resistance $r_0$, so the baseline route resistance is R_p(0) = n_p · $r_0$. All mtDNA-encoded subunits share one mutational environment—the same replication-error rate and the same matrix oxidative load—so each accumulates conductance-degrading lesions at a common per-subunit rate λ, each lesion adding an increment Δr to that subunit's resistance.

**Derivation.** Summed over the chain, the route resistance at age t is

$$R\_p(t) = n\_p \cdot r_0 + n\_p \cdot \lambda \cdot \Delta r \cdot t = n\_p \cdot r_0 \cdot (1 + (\lambda \Delta r / r_0) \cdot t) = R\_p(0) \cdot (1 + \mu t),$$

with $\mu = \lambda \Delta r / r_0$. The subunit count n_p enters both the lesion influx (numerator) and the baseline (denominator) and cancels exactly: μ is independent of target size and series depth. The larger mutational target of the NADH route is matched one-for-one by its larger baseline resistance, so its fractional rate of impedance gain equals that of the shorter CoQ route. Driving voltage enters only through delivered power ($P \propto V\_redox^2/R$), not through the fractional-resistance trajectory, so the higher NADH-route voltage (~1.14 V versus ~0.77 V for the CoQ route; S3.1a) likewise leaves μ unchanged.

**Consequence: a fixed ratio.** Because both routes obey the same (1 + μt) law,

$$R_1(t) / R_2(t) = R_1(0) / R_2(0) = \text{constant for all t.}$$

The two impedances remain locked in fixed proportion across the entire trajectory. Whichever gate is read out first therefore reaches its reporting threshold first because of where that threshold sits, not because its underlying conductance erodes faster. The $NAD^+$ gate is constitutive not because it escapes demand—proliferative aspartate synthesis loads it too—but because it is under load at rest: the $NADH/NAD^+$ couple carries the trunk flux of maintenance catabolism and is continuously consumed non-redoxly by sirtuins, PARPs, and CD38, a drain the recyclable CoQ pool never bears. The CoQ gate is demand-revealed because its one biosynthetic reader, DHODH, carries negligible flux until proliferation recruits de novo pyrimidine synthesis, leaving an over-reduced CoQ pool biosynthetically silent at rest. The two are thresholds on one synchronously degrading system—one loaded at baseline, one latent until demand—so the asymmetry between them is one of visibility, not rate, exactly as the main text states. The $NAD^+$ gate and the CoQ gate are accordingly not a chronological ordering of independent lesions but two readouts of a single parallel erosion.

**Boundary of validity.** Parity holds while lesion accumulation is approximately linear and the two routes remain electrically independent upstream of the shared $III_2$–IV segment. Once that shared segment is itself significantly damaged (the late, shared-segment phase), both routes lose their common downstream conductance, the (1 + μt) scaling breaks down, and the trajectory passes into the late-stage divergence and Δψ volatility of late-stage decompensation.

#### S3.2 Redox-gated biosynthetic bottlenecks

Matrix $NAD^+$/NADH back-pressure inhibits multiple nodes in central carbon metabolism, including PDH, IDH3, α-KGDH, and MDH2. Beyond energy, these nodes gate precursor availability: citrate export (lipid synthesis), α-ketoglutarate flux (amino acid and epigenetic substrates), oxaloacetate/aspartate production (nucleotide synthesis), and one-carbon throughput. Because these are branch-gated processes, biosynthetic flux can fail while ATP remains buffered, yielding a ‘regenerative scope’ failure mode.

##### *S3.2.1 The four-node bottleneck predicts a constrained metabolite signature*

Because PDH, IDH3, α-KGDH, and MDH2 sit at branch points, their redox inhibition creates a stereotyped pattern that is often misread as heterogeneous ‘mitochondrial dysfunction’. In the impedance/back-pressure framing, the pattern is expected to be reproducible across tissues and species, with variation primarily in compensation capacity.

A minimal signature consistent with upstream NADH back-pressure includes:

- PDH gate: increased lactate production and/or alanine transamination (pyruvate diverted to cytosolic NADH disposal).
- IDH3 gate: citrate accumulation and export (cataplerosis), increasing cytosolic acetyl-CoA availability for lipid or cholesterol synthesis.
- α-KGDH gate: reduced succinyl-CoA throughput with relative α-ketoglutarate/glutamate retention and altered one-carbon handling.
- MDH2 gate: constrained oxaloacetate and aspartate availability, limiting nucleotide synthesis and cell-cycle/anabolic throughput.

These are directionality claims. They are best evaluated with paired reserve and pressure measures (e.g., VO2max plus lactate:pyruvate) and with isotope tracing into citrate, aspartate, nucleotides, and fatty acids where feasible.

##### *S3.2.2 Carrier-linked biosynthesis creates oncology-adjacent dependencies*

Several biosynthetic reactions are obligatorily coupled to the respiratory chain through shared carrier pools. This coupling means that conductance loss can constrain biosynthesis even when ATP appears adequate. A canonical example is dihydroorotate dehydrogenase (DHODH), which requires oxidized CoQ as an electron acceptor to sustain de novo pyrimidine synthesis.

In proliferative systems, this coupling produces well-known dependencies (e.g., respiration dependence of aspartate supply and DHODH activity). MSTA predicts a parallel, non-malignant phenotype in aging tissues: reduced proliferative and reparative capacity (immune expansion, epithelial repair, hematopoietic output) driven by carrier-level back-pressure rather than by primary lesions in each biosynthetic enzyme.

The practical implication is interpretive: ‘cancer-like’ redox programs (lactate handling, lipogenic flux, nucleotide stress) may appear as pressure-management signatures in aging without implying that the tissue is executing a growth program. This motivates explicit cross-domain tests rather than rhetorical analogies. The full set of these intermediate- and carrier-linked dependencies — each withdrawn input, the machine it stalls, and the domain that fails — is catalogued in Table S5.

##### *S3.2.3 Redox relief valves: a minimal taxonomy*

When forward electron flow becomes resistive, cells can temporarily preserve cofactor availability by diverting electrons into alternative sinks (‘relief valves’). These can be organized by time scale, compartment, and mechanism:

- Rapid cytosolic NADH sinks: lactate formation (LDH) and related shunts that preserve glycolytic throughput.
- Matrix/export sinks: ketogenesis and reductive packaging of equivalents into exportable metabolites (context dependent).
- NADPH-consuming sinks after NADH-to-NADPH pressure transfer: fatty-acid synthesis/desaturation, mevalonate/cholesterol flux, and defensive reductive programs.

In MSTA, these sinks are compensatory but costly: they spend carbon, ATP, and/or membrane potential to lower redox pressure. An explicit treatment of citrate export, lipogenesis, and related cataplerotic programs is provided in Supplementary Note S5.1.1. The $NAD^+$/NADPH Fork framework (Supplementary Note S7) provides a systematic catalog of 72 opposing enzyme systems whose equilibria shift directionally under cofactor imbalance, generating specific aging phenotypes.

##### *S3.2.4 ROS as the primary pressure valve*

The most primitive electron disposal mechanism is ROS generation itself. When Complex I impedance throttles forward electron flow, electrons leak directly onto $O_2$, generating superoxide. This is commonly interpreted as collateral damage, but the downstream stoichiometry reveals adaptive logic: each leaked electron triggers an antioxidant cascade that consumes multiple NADPH equivalents.

The sequence is as follows. Superoxide is dismutated to $H_2O_2$ by superoxide dismutase (SOD). $H_2O_2$ is then reduced by glutathione peroxidase (GPx), oxidizing 2 GSH to GSSG. Glutathione reductase regenerates GSH at the cost of one NADPH. In parallel, the thioredoxin/peroxiredoxin system imposes additional NADPH demand via thioredoxin reductase. The net effect is an “electron multiplier”: one electron escaping the chain burns ≥2 NADPH molecules during cleanup, each regenerating $NADP^+$ that siphons pressure from the coupled NADH pool through NNT (NADH + $NADP^+$ → $NAD^+$ + NADPH, $\Delta\psi$-driven).

This resolves the long-standing paradox of simultaneous oxidative stress (high ROS) and reductive stress (high NADH/$NAD^+$) in aged tissues. These are not contradictory states but causally linked: ROS is the exhaust of the cell’s attempt to vent reductive backpressure through the NADPH pool. The cost—protein carbonylation, lipid peroxidation, DNA adducts—is the thermodynamic price of pressure relief. ROS damage is thus real but secondary: a consequence of the valve’s operation, not the primary aging lesion.

Within the valve hierarchy, ROS venting is the most primitive and "always-on" mechanism, operating constitutively and scaling with impedance. Downstream valves (lipogenesis, cholesterol flux, cortisol reactivation, prostaglandin rerouting) are slower, tissue-specific, and regulated, but share the same fundamental logic: consuming NADPH to drain the NADH backlog. The complete taxonomy of these directional Fork systems is provided in Supplementary Note S7 and Table S6.

#### Supplementary Note S4. Extended comparative section (avian longevity, ectotherm inversion, and captive tests)

This note expands comparative observations used to motivate or test the SSP constraint structure. The focus is on specifying tests that could refute MSTA rather than on exhaustive review.

##### S4.1 Captive-longevity tests (the caged-parrot prediction)

MSTA distinguishes intrinsic scope/stability constraints from extrinsic ecological hazards. Under protected captivity (low predation, controlled nutrition, veterinary care), realized lifespan should approach the intrinsic ceiling. If avian longevity is driven strongly by intrinsic SSP parameters, captive birds should realize a larger fraction of their predicted MLS than similarly protected mammals. This can be tested using existing zoological and companion-animal datasets with relatively low experimental burden.

##### S4.2 Torpor and hibernation as within-lineage Pace modulation

Hibernation and daily torpor reduce body temperature for extended periods, effectively reducing the time-integrated Pace term. Multiple biological clocks slow during torpor and resume upon arousal. In SSP terms, torpor is expected to increase effective lifespan by reducing cumulative thermal exposure without requiring changes in Stability. Within MSTA, torpor is therefore predicted to preserve scope by reducing the kinetics of conductance erosion and by allowing episodic recovery of redox balance. Comparative tests can examine whether hibernators deviate upward from size-and-temperature predictions when torpor duration is incorporated as an effective temperature integral.

#### Supplementary Note S5. Regenerative scope and the biosynthetic blockade

This note expands the 'regenerative scope' concept by focusing on central carbon metabolism. It motivates why biosynthetic and regenerative failure can precede overt ATP failure in a conductance-limited system.

##### S5.1 Cataplerotic escape and the appearance of compensation

Cells can partially evade the blockade by rerouting carbon. Examples include increasing glycolytic flux and converting pyruvate to lactate to regenerate cytosolic $NAD^+$, increasing glutaminolysis to feed α-ketoglutarate, or relying on fatty acid oxidation and ketones.

These routes can maintain ATP and some TCA turnover, but they are not equivalent to restoring scope. They often increase reduction pressure at the CoQ node (especially $FADH_2$-heavy entry) and can worsen acceptor-side instability if discharge remains constrained.

This logic also explains why some metabolic signatures that resemble ‘Warburg-like’ shifts can appear in non-proliferative aging tissues: they reflect redox management under conductance limitation rather than a deliberate growth program.

##### *S5.1.1 Citrate export and de novo lipogenesis as a pressure valve (and a cancer-like parallel)*

A specific and clinically relevant form of cataplerotic escape is citrate export followed by de novo lipogenesis. When IDH3 and α-KGDH are inhibited under high $NADH/NAD^+$, citrate can accumulate and be exported to the cytosol, where ATP-citrate lyase regenerates acetyl-CoA. This route is often interpreted purely as ‘energy storage’. In the impedance model, it has an additional thermodynamic role: it functions as an ATP- and NADPH-consuming sink that can lower $\Delta\psi$ and relieve matrix back-pressure.

Conceptually, lipogenesis can vent pressure through three coupled sinks: (i) carbon disposal (cataplerosis of citrate that cannot be fully oxidized), (ii) reductant disposal (NADPH demand that draws on reducing equivalents via transhydrogenation or malic-enzyme-linked cycling), and (iii) mechanical relief (ATP demand that engages ATP synthase, increasing proton re-entry and lowering $\Delta\psi$, which reduces the thermodynamic load on Complex I proton pumping).

This framing provides a mechanistic route from mitochondrial conductance loss to stereotyped clinical phenotypes such as visceral adiposity, steatosis, and dyslipidemia, including situations where caloric intake is not increased. In sedentary aging, the loss of external ATP sinks (muscle work) can shift the burden of $\Delta\psi$ relief toward internal sinks, making lipogenesis and related remodeling programs more dominant. The $NAD^+$/NADPH Fork framework (Supplementary Note S7) systematizes this logic: the “Fat Fork” (β-oxidation [HADH; $NAD^+$] vs. fatty acid synthesis [FASN; NADPH]) and the “TCA Flow” fork (IDH3 [$NAD^+$] vs. IDH1/2 [$NADP^+$]) both illustrate how cofactor imbalance forces carbon toward storage pathways regardless of caloric input.

A closely related sink is mevalonate/cholesterol flux, which is also ATP- and NADPH-intensive and can therefore participate in pressure management while simultaneously remodeling membranes, steroidogenic substrate pools, and signaling lipids. These pathways overlap with canonical oncology metabolic programs (lipogenesis and mevalonate dependence), suggesting an explicit, testable bridge: aging tissues may partially recapitulate ‘cancer-like’ redox-sink strategies as survival or stabilization programs rather than as proliferation programs.

Operational predictions include: (i) increased citrate export and fractional de novo fatty-acid synthesis under conditions of elevated lactate:pyruvate (pressure), (ii) coupling between pressure-relief interventions (exercise, conductance restoration) and reduced de novo lipogenesis, and (iii) phase dependence: aggressive promotion of $FADH_2$-heavy entry without conductance repair can worsen acceptor-side instability despite transient energetic benefit.

#### Supplementary Note S6. Discriminating tests versus alternative narratives

This note outlines discriminators between MSTA and closely related narratives. Many frameworks acknowledge mitochondrial decline; the value of MSTA is in the specific constraint structure (SSP) and in the circuit-level predictions (pressure, phase, and bypass logic).

##### S6.1 Distinguishing MSTA from 'mitochondrial dysfunction' as a generic descriptor

Generic mitochondrial-decline descriptions are often compatible with many mechanisms. MSTA makes more specific claims: (i) reserve collapses before basal ATP, (ii) mtDNA-linked stability interacts with temperature in a way that can invert in cold regimes, (iii) conductance loss is central and can be represented as rising internal resistance, and (iv) biosynthetic blockades at redox-gated nodes are early and causal rather than secondary.

##### S6.2 Distinguishing MSTA from oxidative damage/rate-of-living narratives

Rate-of-living and oxidative damage narratives predict that higher flux and higher temperature should uniformly shorten lifespan. MSTA predicts that high flux can coexist with long lifespan if conductance and stability preserve scope, and that the effect of mtDNA composition should not be uniformly beneficial (temperature inversion). The strongest discriminator is therefore the sign structure: a positive Stability term in warm endotherms but an attenuated or negative term in cold-adapted ectotherms. The ROS pressure-valve logic (S3.2.4) provides an additional discriminator: MSTA predicts that ROS generation is a secondary consequence of the cell's attempt to vent reductive backpressure, not the primary aging lesion. This explains why antioxidant interventions consistently fail to extend lifespan: they address the exhaust rather than the blockage.

##### S6.3 Distinguishing MSTA from signaling-centric 'hyperfunction' narratives

Signaling-centric theories emphasize late-life overactivation of growth pathways. MSTA does not deny signaling contributions; rather, it predicts that many late-life signaling shifts are responses to a shrinking energetic budget. A discriminator is ordering: if conductance-first interventions restore reserve and normalize downstream signaling, this supports MSTA; if signaling manipulation reliably restores reserve without affecting conductance or redox pressure, this challenges MSTA. The hallmarks-collapse observation (Main Text Discussion 3.2) provides a strong positioning statement: six of twelve recognized hallmarks of aging emerge as downstream consequences of rising ETC impedance under the MSTA framework, suggesting that MSTA does not compete with the hallmarks taxonomy but provides the upstream constraint that orders and connects them.

#### Supplementary Note S7. The $NAD^+$/NADPH Fork framework: directional shifts in opposing enzyme systems

This note provides the mechanistic derivation and extended catalog for the $NAD^+$/NADPH Fork framework introduced in Main Text Section 2.6.3. While the consilience matrix (Table S4) documents precursor/product dissociations at gated steps, the Fork framework addresses a complementary phenomenon: metabolic nodes governed by opposing enzyme pairs that snap directionally when cofactor ratios shift.

##### S7.1 The "double-lock" mechanism

The primary biophysical lesion in aging—rising Complex I impedance—creates NADH backpressure (elevated NADH/$NAD^+$). This is coupled to the NADPH pool through three mechanisms: (i) nicotinamide nucleotide transhydrogenase (NNT), which runs forward using $\Delta\psi$ (NADH + $NADP^+$ → $NAD^+$ + NADPH), transferring pressure from the NADH pool to the NADPH pool; (ii) the isocitrate shuttle, where mitochondrial IDH3 ($NAD^+$) stalls while cytosolic IDH1/2 ($NADP^+$) continue, generating NADPH from exported citrate-derived isocitrate; and (iii) the malic enzyme shuttle, where mitochondrial ME2 ($NAD^+$) stalls while cytosolic ME1 ($NADP^+$) diverts malate to NADPH generation. The net result is a "double-lock": simultaneous $NAD^+$ depletion and NADPH elevation from a single upstream event.

Under the double-lock, the $NAD^+$-dependent arm of each pair stalls as its substrate is depleted while the NADPH-dependent arm accelerates as its substrate accumulates, forcing the equilibrium in a single direction independent of upstream signaling—thermodynamic compulsion rather than enzyme regulation. Because the cofactor imbalance equilibrates across compartments through the shuttle systems, many such opposing-enzyme systems shift in concert, generating the coordinated endocrine, inflammatory, and signaling syndrome of aging. The complete catalog of 72 verified opposing systems is provided as Supplementary Data (Table S6 Complete NAD+ NADPH Fork catalog - 72 verified opposing enzyme systems.xlsx); the systematic Rhea-database curation behind it is documented in Table S7.

### Supplementary Tables

#### Table S1. Comparative vertebrate dataset: maximum lifespan, body mass, body temperature, and mitochondrial DNA base composition

Note: Provided as Supplementary Data (AnAge_Based_Table.csv). This is the source dataset for the Scope–Stability–Pace (SSP) regression and all comparative analyses. Each row is one vertebrate species (n = 2,739: 1,375 Aves, 1,018 Mammalia, 346 Teleostei) and reports taxonomy (Class, Order, Family, scientific and common name); maximum lifespan

(MLS) with derived lnMLS, initial mortality rate (IMR), and mortality-rate doubling time (MRDT); adult body mass (BM) and metabolic rate; core or ambient body temperature (Tb) with its literature reference; NCBI accession and heavy-strand mtDNA base composition (genome length, A/C/G/T counts and fractions, GC% and AT%, GC and AT skew); and nuclear-genome size and GC% for mammals. MLS, BM, and basal metabolic rate were drawn from AnAge (build 14); mtDNA GC% from MitoAge; mammalian and avian Tb from Clarke & Rothery and primary sources; teleost ambient temperature was curated from FishBase, with the per-row assignment method recorded. The canonical mammal-only SSP fit uses the 379 mammals with complete data on all four predictors (BM, GC%, Tb, MLS).

#### Table S2. SSP coefficients across 100 posterior trees (medians; 95% ranges in brackets). n = 368 species on every tree.

| Coefficient | OLS | BM-PGLS | λ-PGLS | OU-PGLS |
|---|---|---|---|---|
| α (ln BM) | +0.149 | +0.127 [+0.122, +0.137] | +0.138 [+0.136, +0.140] | +0.137 [+0.133, +0.143] |
| β (GC%) | +0.104 | +0.060 [+0.053, +0.110] | +0.053 [+0.051, +0.055] | +0.068 [+0.063, +0.100] |
| γ (Tb) | +0.091 | +0.021 [+0.012, +0.028] | +0.039 [+0.037, +0.041] | +0.030 [+0.020, +0.039] |
| $R^2$ | 0.69 | 0.62 | 0.64 | 0.65 |

#### Table S3. Circuit-predicted consequences of rising R_total

| Principle | Circuit prediction | Observed aging phenotype |
|---|---|---|
| Ohm's law (V = IR) | High upstream voltage (NADH), low current (flux) at high R | Elevated NADH/$NAD^+$, hyperpolarized resting Δψ, reduced maximal respiration |
| Joule dissipation ($P = I^2R$) | Energy wasted as heat/ROS even at reduced flux | Inflammaging, chronic low-grade oxidative stress |
| Current conservation (Kirchhoff) | I_in = I_forward + I_leak; throttled forward flow forces rising leak | Obligate superoxide at Complex I FMN and Q-site, even at rest |
| Impedance mismatch | Source R ≫ load R → inefficient power transfer and upstream pool depletion | $NAD^+$ depletion, sirtuin/PARP substrate limitation, impaired DNA repair |
| RC time constant | τ = R × C; high R → slow charge/discharge kinetics | Metabolic inflexibility, delayed post-exercise and post-prandial recovery |

| Principle | Circuit prediction | Observed aging phenotype |
|---|---|---|
| Maximum power theorem | P_max = $V^2/(4R)$; P_max falls inversely with R | Non-linear frailty acceleration from modest further cellular decline |

#### Table S4. Matrix-localized precursor/product dissociations gated by the mitochondrial NADH/$NAD^+$, CoQ, and Δψ state

Note: Reactions are restricted to the mitochondrial matrix, where reductive stress concentrates—the matrix sits far more reduced than the cytosol even at rest (free $NAD^+$/NADH ≈ 8 versus ≈ 725 in cytoplasm; Williamson, Lund & Krebs, 1967). Each step is throttled as its gating currency—the matrix NADH/$NAD^+$ couple, the CoQ pool, or Δψ/matrix NADPH—is depleted. Entries are candidate dissociations of differing testability rather than uniformly validated biomarkers, and fall into three classes. Thermodynamic equilibrium readers (‡: β-hydroxybutyrate:acetoacetate and glutamate:α-ketoglutarate, which equilibrate with the matrix NAD pool and read its poise directly; lactate:pyruvate reports the cytosolic pool) are pool- and network-independent and the most robust. Boundary accumulators—fed by a non-gated source and drained only through a gated step (e.g., citrate, branched-chain amino acids, 4-hydroxynonenal, long-chain acylcarnitines)—have a robust direction. Interior-node ratios, whose substrate and product are each flanked by gated steps so that their steady-state direction is set by anaplerosis, cataplerosis, and the contracting NAD(H) pool, are listed as mechanistic predictions, not validated readouts. β-hydroxybutyrate:acetoacetate is developed as the cleanest single test in §3.9.

| # | System / pathway | Substrate (↑ under reductive stress) | Product (↓) | Matrix enzyme / gate | Gating currency | Ratio biomarker |
|---|---|---|---|---|---|---|
| 1 | Glycolytic entry (PDH) | Pyruvate (→ lactate, alanine) | Acetyl-CoA entry to TCA | Pyruvate dehydrogenase complex | Matrix $NAD^+$ | Lactate : pyruvate (cytosolic spillover) |
| 2 | TCA — isocitrate step | Citrate / isocitrate | α-Ketoglutarate flux | IDH3 ($NAD^+$) | Matrix $NAD^+$ | Citrate : α-KG |
| 3 | TCA — malate step | Malate | Oxaloacetate (→ citrate synthase) | MDH2 | Matrix $NAD^+$ | Malate : OAA |

| | | | | | | |
|---|---|---|---|---|---|---|
| 4 | Glutamate anaplerosis | Glutamate | α-Ketoglutarate (anaplerotic) | GLUD1/2 | Matrix $NAD(P)^+$ | Glutamate : α-KG ‡ |
| 5 | Ketone-body redox | 3-Hydroxybutyrate | Acetoacetate | BDH1 | Matrix $NAD^+$ | β-OHB : acetoacetate ‡ |
| 6 | Glycine cleavage / one-carbon | Glycine | 5,10-$CH_2$-THF (1-C units) | GCS (GLDC/AMT/GCSH/DLD) | Matrix $NAD^+$ | Glycine : serine |
| 7 | BCAA catabolism | BCAAs / branched-chain α-ketoacids | Branched-chain acyl-CoA flux | BCKDH | Matrix $NAD^+$ | BCAA : C5-acylcarnitine |
| 8 | Aldehyde detoxification | Reactive aldehydes (4-HNE, acetaldehyde) | Carboxylic acids (cleared) | ALDH2 | Matrix $NAD^+$ | 4-HNE adducts (rising) |
| 9 | GABA shunt | Succinic semialdehyde (→ GHB) | Succinate (shunt anaplerosis) | ALDH5A1 (SSADH) | Matrix $NAD^+$ | GHB : succinate |
| 10 | Fatty-acid β-oxidation | Long-chain acylcarnitines / 3-OH-acyl-CoA | Complete β-oxidation (acetyl-CoA) | HADH/HADHA ($NAD^+$) + ETF:QO (CoQ) | Matrix $NAD^+$ + CoQ | LC-acylcarnitines : C2 |
| 11 | Proline catabolism | Proline | Glutamate (via P5C) | PRODH (CoQ) + ALDH4A1/P5CDH ($NAD^+$) | Matrix $NAD^+$ + CoQ | Proline : glutamate |
| 12 | TCA — succinate step | Succinate | Fumarate | SDH / Complex II (FAD→CoQ) | Matrix CoQ | Succinate : fumarate |
| 13 | Sulfide clearance | Hydrogen sulfide ($H_2S$) | Thiosulfate / persulfides | SQOR (FAD→CoQ) | Matrix CoQ | $H_2S$ : thiosulfate |

| | | | | | | |
|---|---|---|---|---|---|---|
| 14 | Choline → betaine | Choline | Betaine | CHDH (FAD→CoQ) | Matrix CoQ | Choline : betaine |
| 15 | Mitochondrial steroidogenesis | Cholesterol / cholesteryl esters | Pregnenolone & downstream steroids | CYP11A1/11B/27 via FDXR–FDX1 | Matrix NADPH + Δψ | Cholesterol : pregnenolone |
| 16 | Matrix protein import | Cytosolic precursor proteins | Mature matrix / IMM proteins | TIM23 / PAM presequence translocase | Δψ | Precursor : mature |
| 17 | Fe-S cluster supply | Labile iron (matrix) | Fe-S–loaded proteins | ISCU assembly → ABCB7 export | Δψ / matrix redox | Aconitase activity : labile iron |
| 18 | Heme completion | Protoporphyrin IX + labile iron | Functional heme | Ferrochelatase (ALAS upstream) | Δψ (IMM) | Zn-protoporphyrin : heme |

#### Table S5. Carrier- and intermediate-linked biosynthetic dependencies of cataplerotic Krebs-cycle output

Note: Provided as Supplementary Data (Table S5 Carrier- and intermediate-linked biosynthetic dependencies of cataplerotic output.xlsx). For each cataplerotic Krebs-cycle intermediate withdrawn as the $NAD^+$-gated dehydrogenases stall (§2.6; Note S3.2), the table records the biosynthetic machine deprived of substrate (Missing_Input → Stalled_Machine), the stoichiometric reaction that consequently fails (Hidden_Cost_Equation), and the physiological domain that loses output (Failed_Domain) — for example, oxaloacetate withdrawal stalls GOT2 aspartate transamination (nucleotide synthesis), citrate synthase (lipid entry), the purine nucleotide cycle (ammonia detoxification), and N-linked glycosylation (glycoprotein stability). The catalog substantiates the carrier- and intermediate-linked dependencies of Note S3.2.2, by which conductance loss constrains biosynthesis through shared intermediate and carrier pools even when ATP appears adequate.

#### Table S6. Complete $NAD^+$/NADPH Fork catalog: 72 verified opposing enzyme systems

Note: Provided as Supplementary Data (Table S6 Complete NAD+ NADPH Fork catalog - 72 verified opposing enzyme systems.xlsx). This table catalogs 72 verified human metabolic nodes governed by opposing $NAD^+$- and NADPH-dependent enzymes, organized into five tiers reflecting proximity to the primary lesion and clinical impact; under the double-lock (simultaneous $NAD^+$ depletion and NADPH elevation; Supplementary Note S7), these equilibria shift directionally from the $NAD^+$-dependent toward the NADPH-dependent direction.

#### Table S7. Fork-catalog curation: systematic Rhea-database classification of $NAD^+$/NADPH-coupled reactions

Note: Provided as Supplementary Data (Table S7 Fork-catalog curation - Rhea-database classification of NAD+ NADPH reactions.xlsx). Documents how the Fork catalog (Table S6) was derived: every Rhea-database reaction turning over an NAD/NADP cofactor was classified by direction — $NAD^+$-dependent oxidation (NAD(+)→NADH; stalls under MSTA), NADPH-dependent reduction (NADPH→NADP(+); continues), or $NADP^+$-dependent oxidation (electron acceptor; not directly antagonistic) — and grouped into twelve physiological categories (steroids, bile acids, prostaglandins, aldehydes, sugars/polyols, proline/glutamate, quinones, glutathione, folate, retinoids, fatty acids, amino acids). The spreadsheet provides the full classified reaction set (787 reactions with Rhea ID, equation, and EC/enzyme), per-category counts, and the classification rules; the directional antagonistic pairs distilled from it are the 72 opposing systems of Table S6.

#### Table S8. Curated human NAD(P)(H)-interacting proteome (306 proteins, three panels)

Note: Provided as Supplementary Data: Table S8 Curated human NAD(P)(H)-interacting proteome (306 proteins, three panels).xlsx. Proteins are retained only on direct molecular UniProt evidence—an annotated NAD/NADP/NADH/NADPH binding site, a catalytic reaction that turns over the cofactor, or an NAD(P) cofactor; holoenzyme reactions are not inherited across structural or accessory subunits (Complex I collapses to NDUFV1), and proteins that receive electrons via a partner reductase (cytochrome P450s, heme oxygenases) or that bind S-adenosylmethionine (SAM methyltransferases) are excluded. Panel S8a lists 236 redox-cofactor binders across 16 structural fold families—the set underlying §2.6.2; panel S8b lists 49 $NAD^+$-consuming/cleaving enzymes (sirtuins, PARPs, CD38/BST1, SARM1, TIR-domain NADases) and panel S8c lists 21 biosynthesis, transport, and sensor proteins—the $NAD^+$-depletion axis developed in §3.6 and Supplementary Note S7.

#### Table S9. Annotated human $NAD^+$/NADH enzyme catalog with reactions and stalling consequences

Note: Provided as Supplementary Data (Table S9 Annotated human NAD+ NADH enzyme catalog with reactions and stalling consequences.xlsx). For each human enzyme or complex that turns over the $NAD^+$/NADH couple, the table lists the gene(s), EC number, balanced reaction, and the predicted physiological consequence when the reaction stalls under $NAD^+$ depletion — the functional-annotation layer behind the ordered load-shedding hierarchy of §2.6. Entries span $NAD^+$-dependent dehydrogenases, $NAD^+$ consumers (PARPs, sirtuins, CD38, SARM1), $NAD^+$ biosynthesis, Complex I, and NADH sensors/regulators. These are the mechanistically characterised subset of the redox-cofactor binders catalogued in Table S8; the two differ in scope because Table S8 is a UniProt binding-evidence census whereas this table is a reaction-and-consequence annotation.
